# A systems approach uncovers drivers of the multifocal emergence of an astrocyte state contributing to cognitive decline in Alzheimer’s disease

**DOI:** 10.1101/2025.02.24.639868

**Authors:** Natacha Comandante-Lou, Tsering Lama, Sarah E. Heuer, Kevin Chen, Jinho Kim, Jinglong Zhang, Bin Hu, Shuhui Liu, Swagata Dey, Kristen J. Brennand, Julie Schneider, Lisa L. Barnes, Bin Zhang, Minghui Wang, Hongyan Zou, Roland H. Friedel, Terrence L. Fisher, Vikas Mishra, Leslie Balch, Yiyi Ma, Aiqun Li, Masashi Fujita, David Bennett, Vilas Menon, Hans-Ulrich Klein, Ya Zhang, Elizabeth E. Evans, Maurice Zauderer, Tracy Young-Pearse, Mariko Taga, Philip L. De Jager

**Author notes:** Correspondence, Lead Contact: Philip L. De Jager, MD PhD.

## Abstract

Diverse cellular states have been mapped in Alzheimer’s disease (AD) pathology, but their links to cognitive decline and the tissue signals driving their emergence remain unclear. We assembled a unified single-nucleus atlas from 867 diverse brains. Meta-analysis identified new cell-state associations with AD and prioritized a non-canonical reactive *SLC38A2^high^SMTN^high^CACNA1D^high^*astrocyte (Ast10) as a key link between neuropathologies and cognitive decline. To identify regulators of Ast10 emergence, we modeled ligand-receptor relationships. Independent datasets and two spatial modalities validated these drivers and revealed Ast10’s multifocal distribution throughout the neocortex. Mechanistic validation in mouse and human astrocytes confirmed *PLXNB1*, *JAM2* and *SEMA4D* as regulators. In cerebrospinal fluid from a SEMA4D-blocking antibody trial, Ast10 markers are pharmacologically modifiable, and their reduction tracks with cognitive stabilization. Ast10 may contribute to cognitive decline through disrupted presynaptic neurotransmitter release and may be a convergence point for AD and other neurodegenerative pathologies, a promising target for enhancing resilience to dementia.

## Introduction

Alzheimer’s Disease (AD) changes the cellular composition of the brain^1,2^. The frequency of many different cell states of various cell types, including neuronal, glial and vascular subpopulations has been associated with AD^1,3–11^; yet, major questions remain regarding the origins of these various AD-associated cell states and how they interact with one another to create cellular communities that may drive the trajectory towards AD^1^. In particular, characterizing these cellular changes to understand which cell subpopulations to target to preserve cognition in the face of AD pathology remains a major challenge since most studies focus on pathologic outcome measures^12,13^.

To elucidate the sequence of changes in cell subpopulations leading to AD, we previously built a cell atlas of 1.65 million single-nucleus transcriptomes from the dorsolateral prefrontal cortex (DLPFC) of 437 participants in the Religious Orders Study or the Rush Memory and Aging Project (ROSMAP)^14,15^. These participants are without dementia at study entry and agree to brain donation; as a result, they represent a wide spectrum of aging brain states found in older individuals. Such interindividual variability allows us to reconstruct the trajectory of brain aging and to dissect the cellular and molecular traits leading to the accumulation of common neuropathologies, such as amyloid and tau that define AD^1^, as well as to subsequent cognitive decline.

Here, we expanded our previous human DLPFC single-nucleus atlas by generating new single-nucleus data and integrating them with two other existing datasets, creating a unified atlas from 867 individuals, nearly doubling the size our previous study^1^. All participants are from ROSMAP or related RUSH studies of diverse participants. This larger atlas enabled a systematic meta-analysis of the cellular change associated with AD traits, increasing the statistical power to detect AD associations with rare cell subsets that would have otherwise been missed. This effort also interrogated ancestrally diverse individuals to explore whether the prioritized cell subpopulations have a similar role among different human populations. This analysis uncovered some new and validated cell states associated with AD-related traits, including a neurovascular cell subtype which unsorted single nucleus RNA sequence (snRNAseq) data typically under-sample.

While these results provide a valuable Resource, we wanted to go beyond associations: we aimed to not only prioritize cell states but also identify candidate intercellular signals that may drive cells towards those disease-associated states. Here, we present a systematic, data-driven approach to prioritize such cell-state transition signals. Glial cells are very plastic, responding to systemic and local cues; however, the local tissue signals regulating these transitions remain poorly understood. We elected to explore the signals driving the emergence of a non-canonical reactive *SLC38A2^high^SMTN^high^CACNA1D^high^*astrocyte state (Ast10), given that it is strongly associated with cognitive decline and mediates part of the relationship between amyloid and tau proteinopathies and cognitive impairment^1^. Since some individuals maintain cognitive performance despite advanced AD pathology, cell states that connect neuropathology to cognitive decline may illuminate mechanisms of cognitive vulnerability and resilience. Identifying signals that promote Ast10 emergence could therefore inform therapeutic targets to limit the pathogenic transition toward this astrocyte state and enhance cognitive resilience in AD.

To identify the ligand-receptor pairs regulating the Ast10 state, we elected to focus on microglia and astrocytes as potential sender cells. This choice was motivated by the fact that these two cell types are deeply interconnected and by our prior study^1^, which showed that Ast10 co-enriches with the lipid-associated microglial state Mic13 (*APOE^high^GPNMB^high^*) in AD, together with growing evidence that specific astrocyte and microglial states co-occur across different neurodegenerative diseases^16–23^. Other neuroglial cell signaling is relevant to disease and will be studied in future analyses. Using the microglial and astrocytic transcriptomes from the 437 human aging cortices in our Discovery study^1^, we systematically inferred the upstream regulatory molecular interactions associated with Ast10 signature and their abundance. This approach narrowed hundreds of possible ligands expressed by either microglia or astrocytes as sender cells and associated receptors to a dozen of high confidence candidates with converging evidence at the donor, pathway and single-cell levels. We then tested these predictions across independent DLPFC snRNA-seq datasets and spatial transcriptomic profiles that assessed whether predicted ligands colocalized with Ast10 astrocytes in human tissue. A transcriptomic assessment of 754,760 spatially registered cells validated the prioritized ligands and uncovered the multifocal, patchy distribution of the Ast10 state throughout the cortex. We next examined two validated ligands and a top-ranked receptor mechanistically using mouse and induced-pluripotent stem cell model systems, human primary astrocytes, and data repurposed from *in vivo* human perturbation in a clinical trial of pepinemab, a SEMA4D-blocking antibody (SIGNAL-AD, NCT04381468). The SIGNAL-AD trial evaluated the safety and tolerability of pepinemab (data not shown, see **Methods**), as well as exploratory proteomic biomarker outcome measures. Altogether, our study provides a prioritization framework using human post-mortem data to systematically identify cell-cell communication signals that promote expansion of a glial state involved in the critical transition from AD pathologies to cognitive decline. Such transition could be exploited therapeutically to enhance resilience to AD and perhaps other pathologies by restoring homeostasis of the brain cellular communities.

## Results

### An expanded human DLPFC snRNA-seq atlas prioritizes Ast10 as a robust cognition-associated astrocyte state in AD

To refine our understanding of the cellular basis of AD, we assembled a staged meta-analysis of four single-nucleus RNA-seq datasets from the dorsolateral prefrontal cortex (DLPFC), comprising 867 brains from older individuals. Our previous study, which we refer to as *Columbia University Irving Medical Center 1 (CUIMC1)*, associated the frequencies of 91 cortical cell subtypes with quantitative measures of three AD-related quantitative traits across 437 participants^1^. Here, we expanded this analysis to nearly double the size of *CUIMC1* by integrating three additional DLPFC snRNA-seq datasets, increasing the power of detecting robust AD-associated cellular states across independent cohorts and ancestrally diverse participants.

All participants have data produced from the same brain region, the dorsolateral prefrontal cortex, and they are all participants in one of five studies based at Rush University: the Religious Order Study (ROS)^14^, the Memory and Aging Project (MAP)^15^, the Minority Aging Research Study (MARS)^24^, the Rush Clinical Core^25^, or the Latino Core Study^26^ (See **Methods** for detailed description of cohorts). All participants undergo the same ante- and post-mortem characterization^24,26–28^, allowing us to combine results across datasets for our three primary outcome measures: the burden of amyloid and tau proteinopathies^29^ as well as the person-specific slope of cognitive decline derived from longitudinal assessments using 19 neuropsychological tests^30–32^. **Supplementary Table 1** presents the demographic and clinicopathologic characteristics of the four sets of participants.

One of the datasets being integrated has not been reported previously, and we refer to it as the *CUIMC2* dataset. It consists of 240 ROSMAP participants; 219 of these participants are not found in the *Discovery* dataset (*CUIMC1*) and were retained for independent analysis (**Figure 1A**). The second dataset was generated at the *Massachusetts Institute of Technology (MIT)* (n=427 ROSMAP participants^3^); 132 of these participants are not represented in *CUIMC1* or *CUIMC2* datasets and are used in our analysis. To begin evaluating whether disease-associated cellular change patterns generalize beyond predominantly European-ancestry cohorts, we also included snRNAseq data generated by the Accelerating Medicines Partnership for AD Diversity project as our third independent set (*Diversity* dataset). Our Data Availability section includes the Synapse ID’s to access all of the data used in this manuscript. We focused the analysis on the 79 participants who are African American (69%), Hispanic White (25%), or American Indian or Alaska Native (5%) based on their self-reported race and who were not already sampled in one of the other three datasets. As described above, only one instance of each participant is used in our meta-analyses (See **Methods**). Notable differences among the four datasets include a significantly higher percentage of female participants (74%) in the *Diversity* dataset and a significantly lower percentage of participants with a diagnosis of AD per the National Institute of Aging’s Reagan Criteria^33^ (45%) in the *MIT* dataset. There was no significant difference in the slope of cognitive decline (*P* = 0.41) across the four datasets (**Supplementary Table 1, Extended Data Figure 1A - E**).

**Figure 1.**
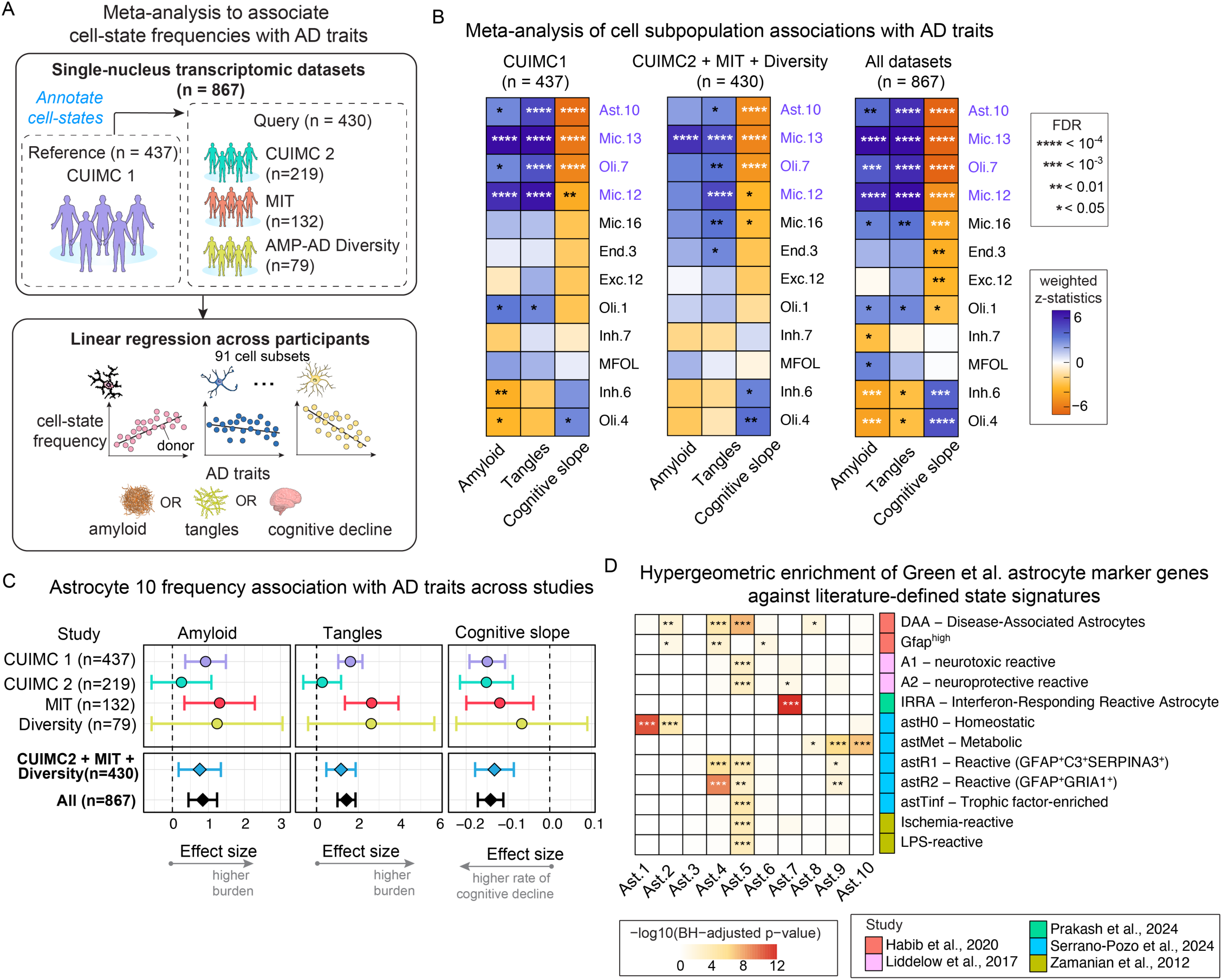
Meta-analysis of DLPFC single-nucleus transcriptomic datasets reveals cell subpopulations associated with AD traits in 867 older individuals. **(A)** Meta-analysis strategy for linking cell-state frequencies to three AD traits (Aβ, tau tangles, and cognitive decline). The Replication set includes three single-nucleus RNA sequencing (snRNAseq) datasets (*CUIMC2*, *MIT*^3^, *Diversity*). Cells were annotated by their cell-state using the Discovery (*CUIMC1*) dataset^1^ as a reference, and associations were tested via linear regression, followed by fixed-effect meta-analysis. **(B)** Meta-analysis results for the Replication set (left) and all participants (Discovery + Replication, right). Heatmaps display effect sizes, ranging from negative (orange) to positive (purple). Positive effect sizes on amyloid and tangles correspond to higher pathology burdens, while negative effect size of cognitive slope corresponds to a higher rate of cognitive decline. Asterisks indicate significance: ****FDR < 10⁻⁴, ***FDR < 10⁻³, **FDR < 10⁻^2^, *FDR < 0.05. **(C)** Forest plots of Ast10 frequency associations with AD traits across individual studies and meta-analyses (bold), shown for Replication (n = 430) and all participants (n = 867). Dots represent effect sizes; lines indicate 95% confidence intervals. **(D)** Hypergeometric enrichment of Green et al. astrocyte subtype marker genes in literature-defined astrocyte state signatures. Heatmap shows enrichment of marker genes from Green et al. astrocyte subtypes Ast1-Ast10 across literature-defined astrocyte signatures. Color intensity indicates -log10 Benjamini-Hochberg-adjusted p value from hypergeometric tests. Asterisks denote adjusted p value thresholds: \**P* < 0.05, ** *P* < 0.01, *** *P* < 0.001, **** *P* < 0.0001. Row annotations indicate the source study for each literature-defined signature.

Armed with these datasets, we first integrated and annotated cells in the three additional datasets using the *CUIMC1* cell-state taxonomy as a reference, allowing us to test all 91 previously defined cortical type and their subtypes^1^ across the expanded atlas (**Extended Data Figure 2-5**; Methods). This represents a broader cell-state-level analysis than our prior bulk RNA-seq-based validation using CelMod^2^, which could only infer the frequencies of 62 cell states. For each dataset, we modeled the association between cell-state frequency and three quantitative measures of AD traits available for these cohorts: (1) an immunohistochemical measure of neocortical amyloid-β (Aβ) (using one of the three monoclonal antibodies against Aβ1-40 or Aβ 1-42: 4G8, 6F/3D and 10D5^29,34,35^), (2) an immunohistochemical measure of neocortical tau (using the AT8^29^ antibody against a phospho-tau epitope), and (3) the slope of cognitive decline prior to death (**Figure 1A**; See **Methods** for full variable descriptions). We then performed staged meta-analyses to test for evidence of association between the frequency of each cell subtype and the AD traits, using a linear regression model while controlling for sex, age, post-mortem interval, education, and library quality.

The first meta-analysis includes the three independent data not used in the *CUIMC1* analysis (n = 430 non-overlapping participants from the three datasets). This new effort recapitulates the top results previously reported in the Discovery analysis^1^, revealing the same top four cell subtypes that are significantly associated with two or more AD traits (based on False Discovery Rate (*FDR*) <0.05): lipid-associated microglial subtypes (Mic13, Mic12), a stress-response oligodendrocyte subtype (Oli7) and the Ast10 astrocytic subtype (**Figure 1B**). To maximize the statistical power and assemble all of our data, we also performed a second meta-analysis using all participants (n = 867). This analysis summarizes all available data and highlights some previously unreported associations, including the role of a microglial state Mic16 (marked by upregulation of the senescence-related gene *SERPINE1*)^1^ and an endothelial state End3 (upregulated genes associated with extracellular matrix organization e.g., *ADAMTS9*)^1^ with faster cognitive decline (**Figure 1B**). We also calculated a measure of heterogeneity (*I*^2^)^36^ among the four datasets and found that Mic16 exhibits the most heterogeneity in effect among the studies (*I*^2^ = 85%) (**Supplementary Table 1**). One example of a cell subtype with a consistent effect (*I*^2^ = 0) across datasets is Ast10 in relation to the rate of cognitive decline (**Figure 1F, Supplementary Table 1**). Its effect size is consistent in all studies; only the Diversity dataset is not significant on its own for this cell state, possibly due to its smaller sample size and the fact that Ast10 is a relatively rare astrocytic state.

Importantly, our analyses begin to provide some evidence that their results may be relevant to participants who are not of European descent (**Figure 1C**). In particular, a higher frequency of Mic13 (an *APOE^high^GPNMB^high^* microglial subtype) consistently has the most significant associations with increased amyloid (*FDR* = 2.7 ×10^−25^) and tau burden (*FDR* = 5.8 ×10^−16^), whereas Ast10 shows one of the strongest associations with faster cognitive decline (*FDR* = 4.4×10^−15^) and less strong association with pathology **(Extended Data Figure 1F**, **Supplementary Table 1**).

Whereas the *APOE^high^GPNMB^high^* Mic13 state has been relatively well described in AD^37,38^ and its association is therefore expected, Ast10 represents an understudied state despite its significant associations with AD traits, especially with cognitive decline. To ensure that Ast10’s robust associations across studies were not due to any technical artifacts, we repeated the meta-analysis on the same four datasets adjusting for additional technical variables (total number of nuclei or astrocytes, sequencing depth, and percent mitochondrial genes) combined with donor-level variables (age, sex, years of education, post-mortem interval); Ast10’s associations with AD traits remained unchanged (**Extended Data Figure 1G**). Given the relatively continuous nature of glial state transcriptional heterogeneity^1^, we assessed how different Ast10-annotation confidence cutoffs (0-0.9)^39^ may affect its associations (**Methods**). Applying a stricter confidence cutoff retained fewer Ast10 nuclei per donor and thus widened the confidence intervals; nevertheless, Ast10’s associations with cognitive decline, amyloid, and tau each remained significant at every cutoff from 0 to 0.9, with effect sizes comparable to or larger than the primary estimates (**Extended Data Fig. 1H**).

To statistically infer the causal relationships of the prioritized glial states (**Figure 1B**) in AD, we previously used structural equation modeling (SEM) to test various positioning of the key glial states along the canonical AD cascade (Aβ ◊ tau ◊ cognitive decline)^1^. In our best fit model on *CUIMC1*, Ast10 lies downstream of Mic13 and tau pathology and upstream of cognitive decline^1^ (**Extended Data Figure 1I**). To assess its reproducibility, we applied the prespecified model to the independent CUIMC2 and MIT datasets, achieving Comparative Fit Index (CFI, ranged 0-1) of 0.94 (**Extended Data Figure 1I**), validating our previous model (CFI = 0.997)^1^. We similarly tested different alternative SEM configurations by swapping Ast10’s position with that of Mic13 and tau; these alternative models however perform much more poorly than our prioritized model (CFI = 0.41-0.45, **Extended Data Figure 1J).** Together, these analyses confirm the robustness of Ast10’s association to AD and reinforce our statistical prioritization of Ast10 as a candidate convergence node downstream of disease-associated microglia (Mic13) and tau pathology, and upstream of cognitive decline.

To determine whether Ast10 represents any known reactive astrocyte program or a distinct AD-associated state, we compared its marker signature against literature-defined astrocyte states. Ast10 was transcriptionally distinct from the more commonly described neurotoxic reactive/A1^16,40^ and disease-associated astrocyte^8^ signatures (**Figure 1D**). Altogether, given our interest to understand why some individuals are resilient to the presence of amyloid and tau proteinopathies, we elected to further explore the interactions between astrocyte and microglial subpopulations that may drive astrocytes towards the Ast10 state, which is an underexplored cellular state immediately upstream of cognitive decline in our model.

### Inference of glial intercellular communications that drive Ast10 transcriptional programs

To uncover the cell-cell signaling that drive the emergence of Ast10, we focus on microglia and astrocytes, given Ast10’s strong association with Mic.13 in our modeling. We start our analysis with the snRNA-seq profiles of 81,255 microglia and 223,063 astrocytes isolated from the 437 donors of the *CUIMC1* dataset; these nuclei are partitioned into 16 transcriptionally distinct microglial and 10 astrocytic states. Within these two cell types alone, there are 315 expressed ligands and 286 receptors that are possible candidates; thus, a systematic approach is required to prioritize the top candidates that will be tested for a possible effect on driving astrocyte transition to Ast10.

**First**, we performed a system-wide cell-cell communication inference to prioritize the upstream ligand-receptor pairs that may regulate the Ast10 signature based on literature curated signaling networks^1^ (**Figure 2A**; **Supplementary Table S2**). **Second**, we used an orthogonal data-driven approach to further prioritize these inferred ligands by using multivariate modeling to predict each donor’s Ast10 frequency based on the combinatorial expression of these ligands. **Third,** to validate these ligands and receptors, we used a series of approaches: (1) *in silico* validation using two of the replication datasets, (2) spatially registered profiling of human tissue at the transcript and protein levels, (3) genetic and ligand perturbations of human and murine astrocytes, and (4) cerebrospinal fluid proteomic data derived from patients receiving a ligand-blocking antibody.

**Figure 2.**
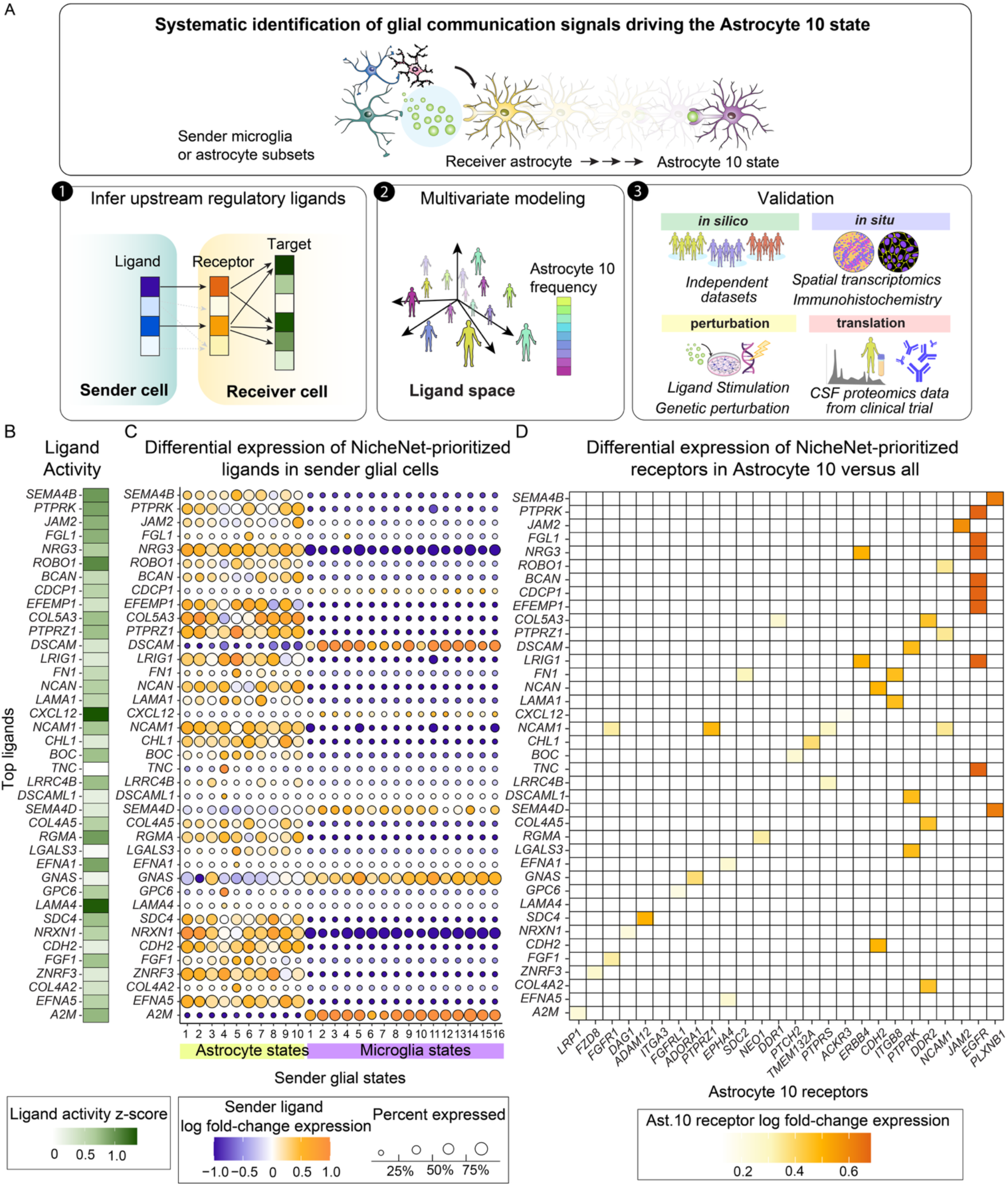
Systematic identification of cell-cell communication signals regulating the *SLC38A2^high^SMTN^high^CACNA1D^high^* Astrocyte 10 state. **(A)** Overview of our systematic approach to identify microglial and/or astrocytic cell-cell signals regulating Astrocyte 10 (Ast10). **(1)** NicheNet^41^ prioritizes ligand-receptor pairs based on their expression and how well their downstream signaling activities recapitulating the Ast10 transcriptional signature. **(2)** Partial Least Squares Regression (PLSR) models predict Ast10 frequency per donor using expression patterns of prioritized ligands or receptors. **(3)** Prioritized signals are then evaluated through orthogonal validation strategies: replication in independent single-nucleus RNA-seq datasets; spatial transcriptomics and immunohistochemistry to test whether candidate ligand-receptor pairs are enriched near Ast10 in human brain tissue; perturbation experiments, including ligand stimulation and genetic depletion of candidate receptors in model systems; and analysis of cerebrospinal fluid proteomics from a phase II clinical trial targeting one of the prioritized pathways. **(B)** Ligand activity z-scores from NicheNet for the top 100 sender-ligand-receptor interactions. A high z-score indicates that a ligand’s predicted target genes are enriched for Ast10 signature genes. A positive z-score reflects above-average activity relative to all other ligands analyzed. **(C)** Differential expression of the top ligands across all analyzed astrocytic and microglial sender states. Color indicates log fold-change (logFC) in ligand expression relative to other sender populations; circle size represents the percentage of cells expressing each ligand. **(D)** Differential expression of the receptors for top-ranked ligands from (B). Color denotes logFC of receptor expression in Ast10 compared to other astrocytic and microglial subsets.

To systematically identify relevant upstream interactions between astrocyte receptors and ligands expressed by microglial and/or astrocytic subpopulations, we used NicheNet^41^, which computationally infers ligand-receptor pairs that are not only differentially expressed but also functionally interacting based on prior knowledge of the ligand’s signaling and gene-regulatory networks^41^. Specifically, we prioritized ligand-receptor pairs based on three key criteria: (1) the degree to which their downstream signaling activity recapitulates the gene expression signature of Ast10 cells (**Figure 2B**), (2) differential expression of receptors in Ast10 compared to all microglial and astrocytic subsets (**Figure 2D**) and (3) differential expression of ligands in a given sender cell subset versus all glial subsets (**Figure 2C**). Based on these criteria, we prioritized 46 ligand-receptor pairs (consisting of 39 ligands and 26 receptors). Among the top ligand-receptor interactions, many are associated with signaling processes such as axon guidance (e.g., *SEMA4B/D – PLXNB1*), cell adhesion (e.g., *JAM2 – JAM2*, *ROBO1 – NCAM1*), and remodeling of the extracellular matrix (e.g., *COL5A3 – DDR*1). Most of the top ligand-receptor interactions involve astrocyte subsets (including Ast10 itself) as senders (**Figure 2C**), suggesting that astrocytes as a cell type may self-modulate the frequency of different cell-states through paracrine and/or autocrine signaling^42^. There are also several microglial-specific signals that are associated with cell guidance or adhesion and neural circuit formation (e.g., *DSCAM, SEMA4D*), G-protein signaling (e.g., *GNAS*) and a gene associated with AD susceptibility and zinc disequilibrium, *A2M*^43,44^. Together, this systemic inference of ligand-receptor pairs and their activities prioritize a set of cell-cell interactions that may drive astrocytes towards the Ast10 state.

### Glial state-specific ligand and receptor gene expression predict variable Astrocyte 10 frequency across aging and AD human brains

Ligand-receptor pairs prioritized by NicheNet represent candidates whose downstream pathway and target genes are consistent with the Ast10 signature gene set, based on literature curated databases. To independently confirm the relevance of these literature-inferred ligand-receptor pairs, we next questioned to what extent they may predict the empirical donor-to-donor variability in Ast10 frequency among ROSMAP participants. In most individuals, Ast10 represents <5% of all astrocytes (**Figure 3A**); with such distribution being consistent across independent datasets. Notably, the enrichment of Ast10 is especially pronounced in participants with a clinical diagnosis of AD (**Figure 3A**). To identify the signaling cues underlying such variability, we used a multivariate approach, partial least-squares regression (PLSR)^45^ to model the relationship between prioritized ligand expression patterns and Ast10 frequency across 171 *CUIMC1* participants who have Ast10 cells (**Figure 3B**). PLSR was chosen because it reduces high-dimensional, correlated ligand expression data into a small number of interpretable principal components that maximally capture the covariance between ligand expression profiles and Ast10 frequency (**Figure 3B**). In this model, each donor is treated as an independent observation and is represented by a vector of ligand summary scores to predict donor-level Ast10 frequency.

**Figure 3.**
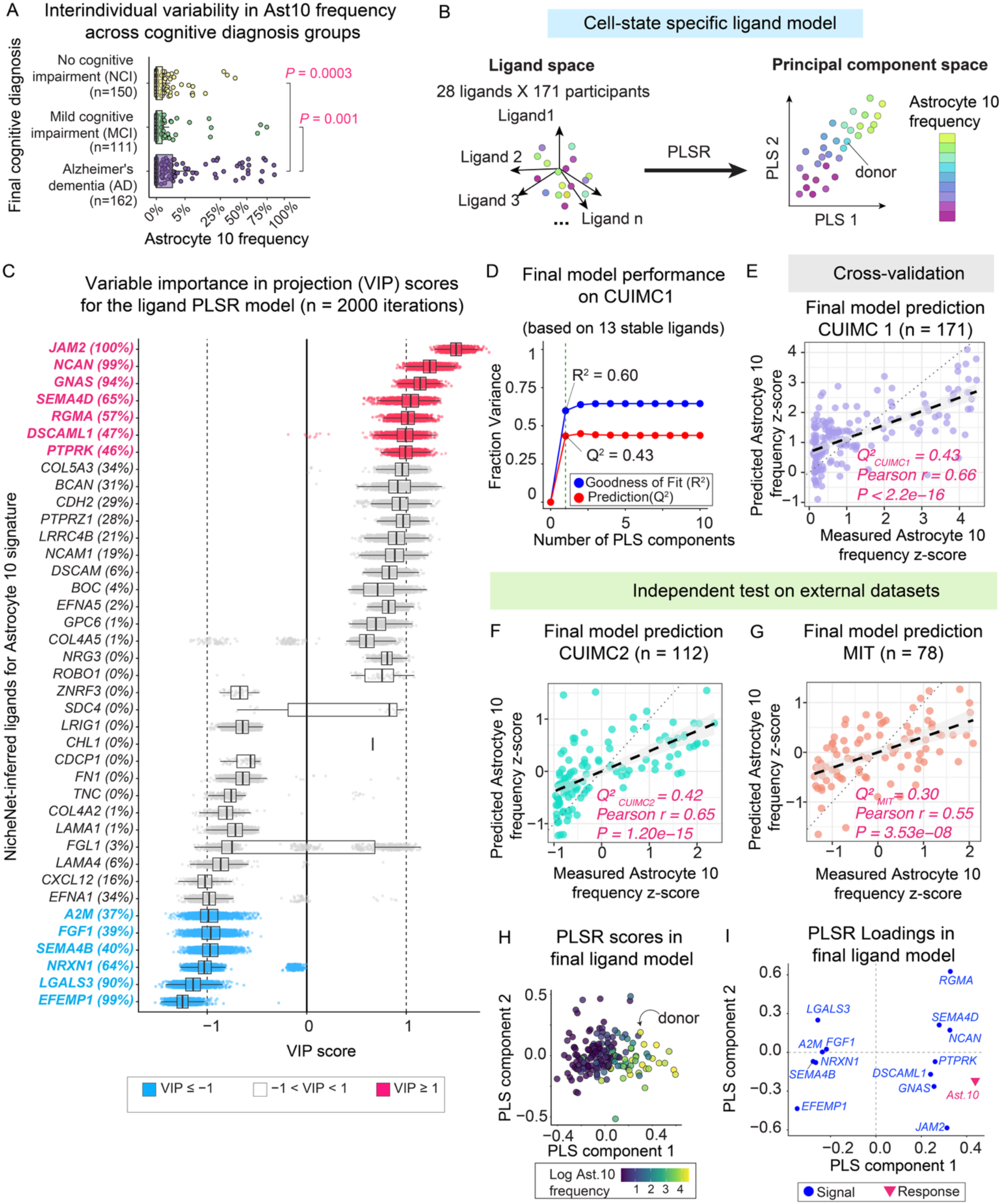
Expression of prioritized ligands in specific glial states predict variation in Ast10 frequency across participants. **(A)** Astrocyte 10 (Ast10) frequency varies across participants in the Discovery cohort, grouped by final consensus cognitive diagnosis: No cognitive impairment (NCI), mild cognitive impairment (MCI), and Alzheimer’s dementia (AD). Clinical diagnoses were determined by neurology experts based on all available clinical data at the time of death^120^. *P*-values: two-sided Wilcoxon signed-rank test. **(B)** Partial Least Squares Regression (PLSR) modeling predicts Ast10 frequency per donor based on prioritized ligand expression. The model input consists of 28 ligand summary scores, each summarizing ligand expression across specific glial states (see Methods). PLSR projects the high-dimensional ligand space into a principal component space, capturing covarying ligands that predict Ast10 variation. Representative principal components (PLS 1 and PLS 2) are shown. **(C)** Stability and importance of ligands in the PLSR model, quantified by Variable Importance in Projection (VIP) scores across 2,000 iterations of repeated random partitioning of training and validation sets. In each iteration, a random subset of donors was used for model training and the remaining donors for cross-validation (see Methods). The magnitude of the VIP score reflects each ligand’s contribution to Ast10 frequency prediction, while its sign indicates positive or negative association with Ast10 frequency. Ligand stability is defined as the proportion of iterations in which |VIP≥ 1. Thirteen ligands selected for the final model based on stability ranking are highlighted. **(D)** Performance of the final PLSR model based on the 13 stable ligands, evaluated by variance explained (R^2^) and variance predicted (Q^2^) using leave-one-out cross-validation, plotted against the number of PLS components. The first PLS component which achieve the optimal performance was highlighted **(E)** Leave-one-out cross-validation performance evaluated by comparing predicted versus empirical (measured) Ast10 frequency z-scores in the *CUIMC1.* **(F-G)** Independent tests of the final model with external datasets not used in model training. Predicted versus empirical Ast10 frequency z-scores in the *CUIMC2* **(F)** and *MIT* **(G)** snRNAseq datasets. Predictions were made using the ligand PLSR model trained on the *CUIMC1* Discovery dataset. **(H)** Donor-level PLSR scores for the first two components of the final ligand model, colored by log-transformed Ast10 frequency. **(I)** PLSR loadings (the first two PLS components) of the final PLSR model for each ligand predictor and the response variable (Ast10 frequency).

For each ligand, we calculate its summary score by averaging its pseudobulk expression across specific microglial or astrocytic subtypes, where the expression of that ligand significantly correlates with Ast10 frequency (**Extended Data Figure 5A**; see **Methods**). To prevent data leakage, we learned these ligand aggregation rules exclusively within each training set and applied them unchanged to held-out individuals of the validation set during each iteration of the repeated training and validation cycle (see **Methods; Extended Data Figure 5A**).

To identify ligands with robust associations with Ast10 frequency, we assessed ligand stability across 2000 iterations of random partitioning of the *CUIMC1* Discovery cohort into training and validation sets (**Extended Data Figure 5A**). In each iteration, we trained a PLSR model and computed Variable Importance in Projection (VIP) scores, which quantify each ligand’s contribution to model predictions (see **Methods**). We ranked ligands by their selection frequency, defined as the proportion of iterations in which the ligand has significant contribution (|VIP| ≥ 1) (**Figure 3C)**.

To define a stable ligand set, we incrementally added ligands according to this stability ranking and evaluated predictive performance as a function of model complexity **Extended Data Figure 5B)**. Leave-one-out cross-validated Q^2^ exhibited a broad plateau from approximately 13 to 28 ligands (Q^2^ ≈ 0.40-0.44; **Extended Data Figure 5B**), indicating limited gains in generalization with additional features. We therefore selected the top 13 ligands (highlighted in **Figure 3C**) within this plateau as the final PLSR model, corresponding to a parsimonious model size with stable cross-validated performance.

Using these 13 ligands, we trained a final PLSR model on all 171 *CUIMC1* donors. The first PLS component maximized explained (R^2^ = 0.60) and cross-validated predicted (Q^2^ = 0.43) variance in Ast10 frequency under leave-one-out cross-validation (**Figure 3D, E**). To further assess the robustness of the final model based on these 13 ligands, we used the trained model to *independently* predict Ast10 relative proportions across non-overlapping individuals from the *CUIMC2* and *MIT* datasets. Importantly, these datasets did not in any way inform our final model, providing independent evaluations of our model. The *Diversity* dataset did not have a sufficient number of individuals with Ast10 and was therefore not considered in this replication effort. In *CUIMC2* (**Figure 3F**), the ligand model retained most of its performance, predicting 42% of the variance (Q^2^ = 0.42) in Ast10 frequency. In the *MIT* dataset (**Figure 3G**), predictive accuracy was reduced (Q^2^ = 0.30); however, predicted and observed Ast10 frequencies remained strongly correlated (Pearson’s *R* = 0.55, *P* = 3.5×10^−8^), suggesting the model captures biologically relevant signal despite cross-data generation site technical variation. Thus, we have demonstrated that our PLSR ligand model based on these 13 ligands is robust, yielding reproducible results in two additional independent test datasets that do not overlap with the set of participants from which the model was derived.

The high predictability of the ligand PLSR model supports a strong association between NicheNet-prioritized ligands and the Ast10 transcriptional program. Importantly, it also indicates a significant link between signaling cues from specific glial states and astrocyte state compositions, explaining the substantial interindividual variability in Ast10 relative proportions (**Figure 3A**).

Accordingly, donors with low to high Ast10 relative proportions can be ordered based on their PLSR scores along the first PLSR component (PLS 1) (**Figure 3H**). The interpretability of PLSR enables us to then dissect the PLSR components in molecular terms based on their loadings, which describe the contribution of each *signal* or *response* variable to a given PLSR component. Consistent with the scores plot, the *response* variable (i.e., Ast10 frequency) is highly and positively weighted along PLS 1 (**Figure 3I**). Loadings of the *signal* variables reveal two distinct groups of ligands that contribute to PLS 1 in opposite directions (**Figure 3I**). Among the 13 stable ligands in the final model, axon guidance and cell-cell adhesion genes (e.g., *JAM2*, *NCAN*, *SEMA4D*) predict Ast10 enrichment (**Figure 3C, I**). In contrast, another group of significant ligands predict Ast10 state depletion, including genes associated with extracellular matrix organization (e.g., *EFEMP1*), synaptogenesis (*NRXN1*) and injury-induced astrocyte reactivity (*LGALS3*^46^, *FGF1*^47^) (**Figure 3 C, I**). Together, analysis of the ligand PLSR model generates hypotheses on the signal-response relationships between combinatorial patterns of specific ligands and the enrichment of Ast10 cell subpopulation.

Next, we assessed whether astrocyte receptor expression likewise predicts Ast10 frequency across donors (**Extended Data Figure 5C**). We reason that receptors that are highly expressed in astrocytes from donors with high Ast10 frequency are potential candidates for mediating signaling pathways that drive astrocytes toward transitioning into the Ast10 state. Applying the same PLSR framework, we identified a stable set of 10 receptors whose expression optimally predicted Ast10 frequency (Q^2^ = 0.65) in *CUIMC1* using the first two PLS components (**Extended Data Figure 5D-F**). Consistent with the ligand model, independent testing showed strong generalization to *CUIMC2* (Q^2^=0.40; **Extended Data Figure 5G**) and significant predictive correlation in *MIT* (Q^2^ = 0.25; Pearson’s *R* = 0.52, *P* = 1.3×10^−7^; **Extended Data Figure 5H**). Individuals with higher Ast10 frequency show elevated PLS1 and PLS2 scores (**Extended Data Figure 5I**), driven by coordinately upregulated receptors (**Extended Data Figure 5J**). These findings show that astrocyte population-level expression of these top 10 receptors is sufficient to predict donor-level enrichment of the Ast10 transcriptional program. Consistently, among the most significant receptors (based on their VIP score stability across iterations), we find *JAM2* (one of our ligands which is reported to homodimerize^48^), and *ITGA3* (for which NicheNet annotations^41,49^ suggest a potential JAM-family interaction^50,51^), *PLXNB1* (*SEMA4D* and *SEMA4B* receptor), *ADORA1* (*GNAS* receptor), *PTPRK* (*DSCAML1* and *LGALS3* receptor) and *EGFR* (*EFEMP1* receptor) whose corresponding ligands are also significant contributors to the ligand model (**Extended Data Figure 5J, Figure 3C**).

Together, these results reveal that the Ast10 subpopulation is associated with cell-cell interactions involving specific microglial states or other astrocytic states. The gene expression of these ligands and receptors in specific microglial or astrocytic states are highly predictive of the relative abundance of the Ast10 subpopulation across hundreds of individuals from different datasets.

### Prioritized ligand pathway activities are associated with Astrocyte 10 signature at the single-cell level

While PLSR established donor-level associations between prioritized ligand-receptor pairs and Ast10, we next used a complementary single-cell approach to assess whether signaling pathways downstream of the 13 significant ligands identified by the PLSR model (*JAM2, NCAN, GNAS, SEMA4D, RGMA, DSCAML1, PTPRK, EFEMP1, LGALS3, NRXN1, SEMA4B, FGF1, and A2M*) show coordinated activities with the Ast10 transcriptional program across individual astrocytes. Specifically, we quantified the mutual information between genes downstream of these ligands and the Ast10 signature across all astrocytes in the *CUIMC1* dataset, leveraging cell-to-cell variability within the astrocyte population. We reasoned that if these prioritized ligands are functionally active, expression of genes within their downstream pathways should be informative of an astrocyte’s Ast10 signature (regardless of it being up- or down-regulated by the ligand) (**Extended Data Figure 6A**). To quantify such information flow at the single-cell level, we used a metric called conditional-Density Resampled Estimate of Mutual Information (DREMI) that is designed to capture the non-linear associations among genes in single-cell data^52,53^ (See **Methods**). For each ligand-pathway gene (excluding Ast10 signature genes themselves), that NicheNet used to connect the prioritized ligands to downstream Ast10 signature genes based on literature-curated signaling pathways, we calculated a DREMI score between its expression and the Ast10 signature score across individual astrocytes (see **Methods** for signature score calculation).

As expected, most of the prioritized ligands’ cognate receptors as well as their downstream signaling mediators and transcription regulators have significantly higher DREMI scores compared to random permutations of the data, indicating that they share certain levels of mutual information with the Ast10 signature within the same cell (**Extended Data Figure 6B**). In particular, top receptors predicted by the PLSR model (**Extended Data Figure S5C**), *JAM2* and *PLXNB1*, have high DREMI scores of 1.17 and 1.09 (among top 0.5% and 1% of all tested receptors, See **Methods**) respectively (**Extended Data Figure 6C**). Notably, given its strong predictability of Ast10 frequency as a ligand (**Figure 3C**) and as a receptor (**Extended Data Figure S5C**), *JAM2* appears to play a role in both juxtacrine signaling from sender cells to Ast10 and intrinsic signaling within Ast10 itself. As predicted by the PLSR model, ligand *FGF1* is associated with decreasing Ast10 frequency (**Figure 3C**), its downstream signaling mediator, *PRKCA*^54^, also decreases with Ast10 signature with a high DREMI score of 1.18 (among top 0.2% of signaling mediators). In summary, the pathway analysis provides evidence that our results are robust to the analytic method used to generate them, supporting the activities of the prioritized ligands in regulating Ast10 state at the single-cell level.

### Prioritized ligands spatially co-localize with Ast10-high regions *in situ*

To validate the group of ligands prioritized by NicheNet and PLSR, we repurposed spatial transcriptomic data (Visium platform) from 32 sections of frozen dorsolateral prefrontal cortex obtained from 17 individuals selected from our Discovery cohort^55^ (**Figure 4A**). Among them, 15 donors had a clinical diagnosis of AD dementia and advanced AD pathology (Braak Stages 4-5), while 2 donors were cognitively unimpaired and showed no pathological signs of AD. (see **Supplementary Table 7** for demographic and clinicopathologic characteristics). This version of Visium captures RNA in an array of 55 µm spots, and the RNA is subsequently reverse transcribed to cDNA and sequenced in a spatially registered manner^56^. We focused on the subset of highly stable ligands that are predicted to upregulate the Ast10 signature (i.e., VIP scores ≥ 1 in at least 50% of model iterations) and asked whether the expression of these ligands in the vicinity of a given spot can predict the expression level of the Ast10 signature of that spot (**Extended Data Figure 7A**). To operationalize this analysis, we assessed data found within a radius of 300 μm from the center of the target spot (corresponding to 3 spot-to-spot distances in the Visium array). The target spot itself was *excluded* from the neighborhood definition to avoid confounding the correlation assessment. We selected a neighborhood radius of 300 μm to encompass sender cells that signal through both immediate cell-cell contact (< ∼150 μm, based on an average human astrocyte territory size diameter of 142.6 μm^57^) and diffusion-mediated signaling of secreted factors (∼100-300 μm^58^).

**Figure 4.**
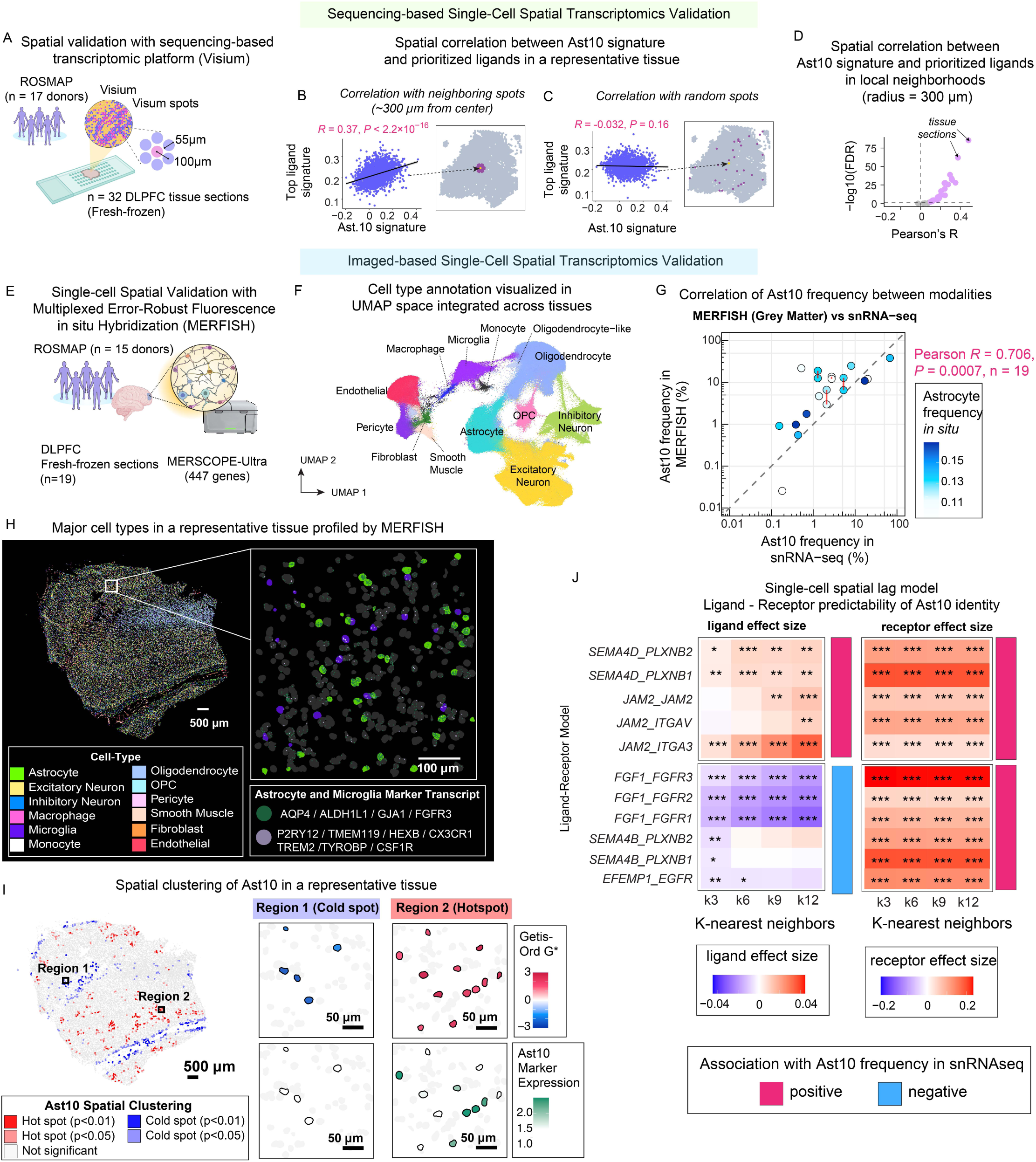
Prioritized ligands preferentially predict nearby Ast10 signature in spatial transcriptomic data. **(A)** Sequencing-based spatial transcriptomic profiling using the Visium platform. 32 fresh-frozen DLPFC brain tissue sections from across n = 17 ROSMAP donors were analyzed. Visium spots are 55µm in diameter with 100µm center-to-center spacing. **(B)** Pearson correlation between Ast10 signature scores at each Visium spot and the mean expression of prioritized ligands across neighboring spots within a 300 µm radius, excluding the focal spot itself. Each point represents one spot in the representative tissue section. Inset: spatially registered tissue section highlighting an example focal spot (yellow) and the neighboring spots (purple) used to compute the neighborhood ligand expression. **(C)** Control analysis corresponding to (B), using randomly selected spots rather than spatial neighbors. Pearson’s correlation between the Ast10 signature score at each Visium spot and the mean expression of prioritized ligands across randomly selected spots from the same tissue section. Inset: the same tissue section, highlighting the focal spot (yellow) and the randomly selected spots (purple) used for comparison. **(D)** Volcano plot summarizing within-slide Pearson’s correlations in Visium data, as described in Extended Data Figure 7A. Each point represents one tissue sample and shows the Pearson’s correlation statistics computed across spots between the focal-spot Ast10 state signature and the neighborhood ligand signature score. The ligand signature was defined using prioritized Ast10-predictive ligands with significantly positive VIP scores in at least 50% of training iterations. For each focal spot, the ligand signature score was averaged across neighboring spots within a 300 µm radius, excluding the focal spot itself. Ast10 state signature was derived from differentially upregulated marker genes (see Methods). Samples with significant correlations (false discovery rate; *FDR* < 0.05) are highlighted. **(E)** Imaging-based spatial transcriptomic profiling at single-cell resolution using MERFISH. 19 fresh-frozen DLPFC tissue sections from n = 15 donors of the CUIMC1 cohort were profiled, using a pre-designed 447-gene panel. **(F)** Integrated UMAP of nuclei across all 19 MERFISH tissue sections, calculated from transcripts assigned by nuclei segmentation. Each data point represents a nucleus, colored by cell type annotation derived from *de novo* clustering and canonical cell-type marker expression. **(G)** Concordance of Ast10 frequency measured by MERFISH (grey matter regions) and snRNA-seq across matched donors (n = 19 sections). Connected points indicate sections from the same donor. Points are colored by the proportion of astrocytes detected by MERFISH. **(H)** Representative tissue section profiled by MERFISH, with each cell colored by their cell type annotation. Insets: highlight microglia (purple) and astrocytes (green) in the zoom-in regions showing expression of their corresponding canonical marker transcripts. **(I)** Same representative tissue as in (H), showing the spatial pattern of Ast10 quantified using local Getis-Ord Gi* z-scores, which assess whether the focal astrocyte’s spatial neighborhood is enriched for high or low Ast10 scores relative to the tissue-wide distribution. Significant positive Gi* values (hotspots: Gi* ≥ 1.96, P ≤ 0.05) indicate spatial clustering of astrocytes with high Ast10 scores, whereas significant negative values (coldspots: Gi* ≤ -1.96, P ≤ 0.05) indicate clustering of astrocytes with low Ast10 scores. Inset: top row, Gi* z-scores for individual astrocytes within the zoomed-in coldspot (Region 1) and hotspot (Region 2); bottom row, the same regions with individual astrocytes colored by their Ast10 scores, calculated as the average expression of the top Ast10 markers *SLC38A2, SMTN, SLC39A11, CACNA1D, MT1E.* **(J)** Spatial lag modeling effect sizes for prioritized ligand-receptor pairs. Models quantify their spatial colocalization with Ast10 scores across multiple neighborhood scales. For each ligand-receptor pair, Ast10 scores in a target astrocyte are modeled as a function of the average ligand expression in the *k*-nearest microglia or astrocytes and receptor expression in the target astrocyte, adjusting for other technical covariates (see Methods). *k* ranges from 3 to 12 neighbors (approximately 59-126 µm). For visualization, ligand-receptor pairs are grouped by whether they are positively or negatively correlated in Ast10 frequency in snRNAseq data analysis (Figure 3, Extended Data Figure 5). ***, *FDR* < 0.001; **, *FDR* < 0.01; *, *FDR* < 0.05)

We observed significant and positive correlations between the top ligand signature within the 300 μm neighborhood and the Ast10 signature (representative sample shown in **Figure 4B**, statistics of all samples shown in **Figure 4D**). In contrast, no significant correlations were observed when associating each spot’s Ast10 signature with the ligand signature in randomly located spots (**Figure 4C**). Importantly, for most samples, we did not observe significant and positive correlations when associating the prioritized ligands with the upregulated signatures of other astrocyte subtypes (**Extended Data Figure 7B**), suggesting that the prioritized ligands are specifically predictive for the Ast10 signature. Interestingly, Ast3 (*PINK1^high^RPL13^high^YWHAG^high^*) appears to be significantly depleted in the target spot, suggesting that Ast10 and Ast3 may be somewhat mutually exclusive in regions with high Ast10 ligand expression. Sensitivity analyses across different neighborhood sizes (ranging from 100-500 μm radii) demonstrated that the prioritized ligands remained significantly and specifically associated with Ast10 at all tested distances (**Extended Data Figure 7C**), indicating robustness of these Ast10-predicting ligands across biologically relevant spatial scales.

To further explore the spatial relationship between individual prioritized ligands and Ast10 signature genes, we used MISTy^59^, an explainable machine learning framework to analyze the importance of each ligand in predicting each Ast10 signature gene. Specifically, we used MISTy to estimate the contribution of ligands within the local neighborhood (radius of 300 μm) on a given Ast10 signature gene, while adjusting for the intrinsic contribution from other Ast10 markers within the same spot. We found that many of the ligands in the PLSR model are significantly predictive (based on MISTy importance scores > 0; see Methods for detailed explanation of the metric) of most of the Ast10 signature genes (**Extended Data Figure 7D**). To assess the significance of the MISTy importance scores across the prioritized ligand set from the PLSR model (**Figure 3C**), we performed a permutation analysis. Specifically, we randomly selected 13 ligands from the full set of possible ligands 100,000 times, calculating the mean total importance score for each set (defined as the sum of importance z-scores across all Ast10 signature genes). The prioritized ligands in the PLSR model exhibited significantly higher average total importance scores compared to most randomly sampled ligand sets (empirical *P-*value = 8.3 × 10^−4^, **Extended Data Figure 7E**).

To further validate the Ast10-predicting ligand-receptor pairs at a *single-cell* spatial resolution *in situ*, we performed Multiplexed Error-Robust Fluorescence In-Situ Hybridization (MERFISH^60^, using the second-generation MERSCOPE-Ultra platform). This imaging-based approach enables highly quantitative, sensitive, and specific measurement of individual transcripts at single-cell resolution. Guided by our computational predictions, we designed a 447-gene panel (**Supplementary Table 8**), including 66 Ast10 differentially expressed genes identified in *CUIMC1* and 165 additional genes capturing candidate Ast10-driving ligands, receptors, and downstream pathways (**Figure 4E**). We profiled 19 tissue sections from 15 *CUIMC 1* donors spanning different points along the cellular trajectories leading to AD (**Extended Data Figure 8A, B**; See **Supplementary Table 8** for donor demographics). In total, we profiled 754,760 cells (71% cells from grey matter, 27% cells from white matter), including 97,387 astrocytes (**Figure 4F, Extended Data Figure 8C,** representative tissue shown in **Figure 4H**, see Methods for cell-type annotation approach).

We first asked whether Ast10 state captured by snRNAseq can also be detected in intact human tissue. To identify Ast10 cells *in situ*, we scored each astrocyte based on the mean expression of five top Ast10 markers (*SLC38A2, SMTN, SLC39A11, CACNA1D, MT1E*). In the Discovery snRNA-seq dataset (*CUIMC1*), these markers individually distinguished Ast10 from other astrocyte states with area under the receiver operating characteristic curve (AUC) values of 0.61-0.77, while their combined averaged expression achieved an AUC of 0.88 (**Extended Data Figure 8D,E**), demonstrating that this five-gene average expression effectively captures the snRNA-seq-defined Ast10 state. We next compared Ast10 frequencies derived from MERFISH with matched snRNA-seq data from the same individuals (see **Methods**). Ast10 frequencies were highly correlated between modalities in both grey matter (Pearson’s *R* = 0.71, *P =* 0.0007; **Figure 4G**) and in white matter (Pearson’s *R* = 0.9, *P =* 2.8×10^−7^; **Extended Data Figure 8F**), providing cross-modality support that Ast10 is not solely a snRNA-seq-specific artifact and that the selected markers identify the Ast10 state *in situ*.

To quantitatively characterize the spatial organization of Ast10 *in situ*, we computed, for each astrocyte, a local Getis-Ord Gi* z-score^61^, which assesses whether the focal astrocyte’s spatial neighborhood is enriched or depleted for astrocytes with higher Ast10 scores relative to the tissue-wide distribution. Significantly positive Gi* values (hotspots, Gi* ≥ 1.96, *P* ≤ 0.05) indicate spatial clustering of astrocytes with higher Ast10 scores, whereas significantly negative values (coldspots, Gi* ≤ -1.96, *P* ≤ 0.05) indicate clustering of astrocytes with lower Ast10 scores (representative tissue shown in **Figure 4I**). Grey matter (GM) and white matter (WM) exhibited markedly distinct Ast10 patterning. Grey matter regions showed significantly greater enrichment of Ast10 in the local microenvironments (GM: median Gi* = 0.34 ± 0.30, WM: median Gi* = -0.92 ± 0.43; paired Wilcoxon test *P* = 7.63×10^−6^) and a higher fraction of Ast10 hotspots (GM: 0.12 ± 0.06, WM: 0.007 ± 0.008; *P* = 7.63×10^−6^) than in the white matter (**Extended Data Figure 8G)**. These results suggest that grey matter provides a more permissive microenvironment for Ast10 induction and that Ast10 cells seem to cluster into microscopic foci in the neocortex. Notably, despite these territorial differences, Ast10 frequencies in grey and white matter regions within the same tissue sections were strongly correlated (Pearson’s *R*= 0.84, *P* < 0.0001, **Extended Data Figure 8H**), suggesting coordinated tissue-level regulation alongside local region-specific factors driving Ast10 accumulation.

Focusing on grey matter astrocytes, we next quantified how local ligand and receptor expression relates to Ast10 scores at the single-cell level. We modeled each grey-matter focal astrocyte’s Ast10 score as a function of (i) its own receptor expression, (ii) the average ligand expression of its nearest astrocyte and microglial neighbors (a spatial lag; *k* = 3-12 neighbors; corresponding approximately to ∼59-126 μm; **Extended Figure 8I**) and (iii) technical covariates, with a random intercept per tissue section (**Methods**). This analysis confirms significant positive spatial lag effects for *JAM2* and *SEMA4D*, reflecting their elevated expression in neighboring cells surrounding Ast10-high astrocytes. In contrast, additional ligands, including *EFEMP1*, *SEMA4B*, and *FGF1*, exhibited significant negative spatial lag effects, or local depletion near Ast10 as predicted by the donor-level PLSR (**Figure 4J; Extended Data Figure 8J**). The spatial-lag effects of these ligands were largely consistent across neighborhood sizes. Notably, top receptors prioritized by the PLSR model to predict Ast10 enrichment at the donor level (e.g., *PLXNB1, JAM2, ITGA3, PTPRK, EGFR*; **Figure 4J; Extended Data Figure 5D**) showed robust positive associations with Ast10 scores across spatial scales, confirming the co-expression of these receptors with Ast10 markers in the target cell at the single-cell level *in situ*.

Taken together, these results derived from different datatypes (snRNAseq, spatially registered RNAseq data and *in situ* hybridization) suggest that we have identified a set of robust ligands expressed in microglia or astrocytes that putatively regulate the Ast10 state. These ligands are prioritized and validated *in silico* and *in situ* based on several orthogonal approaches: 1) their ability to recapitulate the Ast10 signature given by prior evidence of the signaling pathways, 2) their predictability of Ast10 frequencies across individuals in several independent single-nucleus datasets, and 3) their spatial colocalization with Ast10 signature in brain tissue. Importantly, the patchy distribution of Ast10-enriched foci lends a moth-eaten appearance to the AD neocortex (**Figure 4I**). Across these several approaches, we found that molecules involved in tight junctions and cellular adhesion, such as *SEMA4D-PLXNB1*, *JAM2-JAM2* and *JAM2-ITGA3* ligand-receptor pairs consistently showed up as significant contributors to the Ast10 state. Therefore, we elected to focus on these ligand-receptor pairs in further experimental validation.

### Ast10 marker colocalizes with *PLXNB1* in human tissue at the protein level

Since *PLXNB1* is one of the top receptors associated with the Ast10 transcriptional program *in situ*, we next asked whether PLXNB1 protein signal is similarly co-expressed with an Ast10 marker in human tissue at the single-cell level. To this end, we used immunofluorescence (IF) to measure the expression of PLXNB1, Ast10 marker SLC39A11 and astrocyte cell-type marker GFAP, capturing 40 images at 20x magnification of a formalin-fixed paraffin-embedded (FFPE) DLPFC sample derived from a donor with AD. SLC39A11 was selected as a marker for the Ast10 signature due to the lack of specific antibodies against other upregulated transcriptional markers (e.g., *SLC38A2, SMTN, CACNA1D*) for IF on human brain sections. Nevertheless, SLC39A11 is significantly upregulated in Ast10 compared to all of the other astrocyte states and achieved one of the highest AUC scores of 0.77 in snRNAseq data, confirming its specificity in distinguishing Ast10 from other astrocyte states (**Extended Data Figure 8D**, **Supplementary Table S2**).

For image analysis, the images were segmented to identify individual GFAP+ astrocytes using a custom pipeline implemented in CellProfiler^62^; the resulting cell-by-mean fluorescence intensity feature matrix was used in downstream analyses (See Methods, **Figure 5A**). To determine whether PLXNB1 protein expression level in a given astrocyte is predictive of its expression of SLC39A11 (used as a marker protein for the Ast10 signature), we fit a mixed effects model controlling for the fixed effect of GFAP levels and the random effect across images (See Methods). We observed a strongly positive and significant effect size (*P* < 2×10^−16^) of PLXNB1 in predicting SLC39A11 expression level across individual astrocytes, confirming the association between astrocytic PLXNB1 receptor and the Ast10 signature in human tissue at the protein level (**Figure 5B, C**). Interestingly, GFAP expression is moderately but inversely correlated with SLC39A11, suggesting that the Ast10 signature and the classic reactive astrocyte state, characterized by elevated GFAP levels, may capture distinct astrocytic states^8,16,63^. (**Extended Data Figure 9A**).

**Figure 5.**
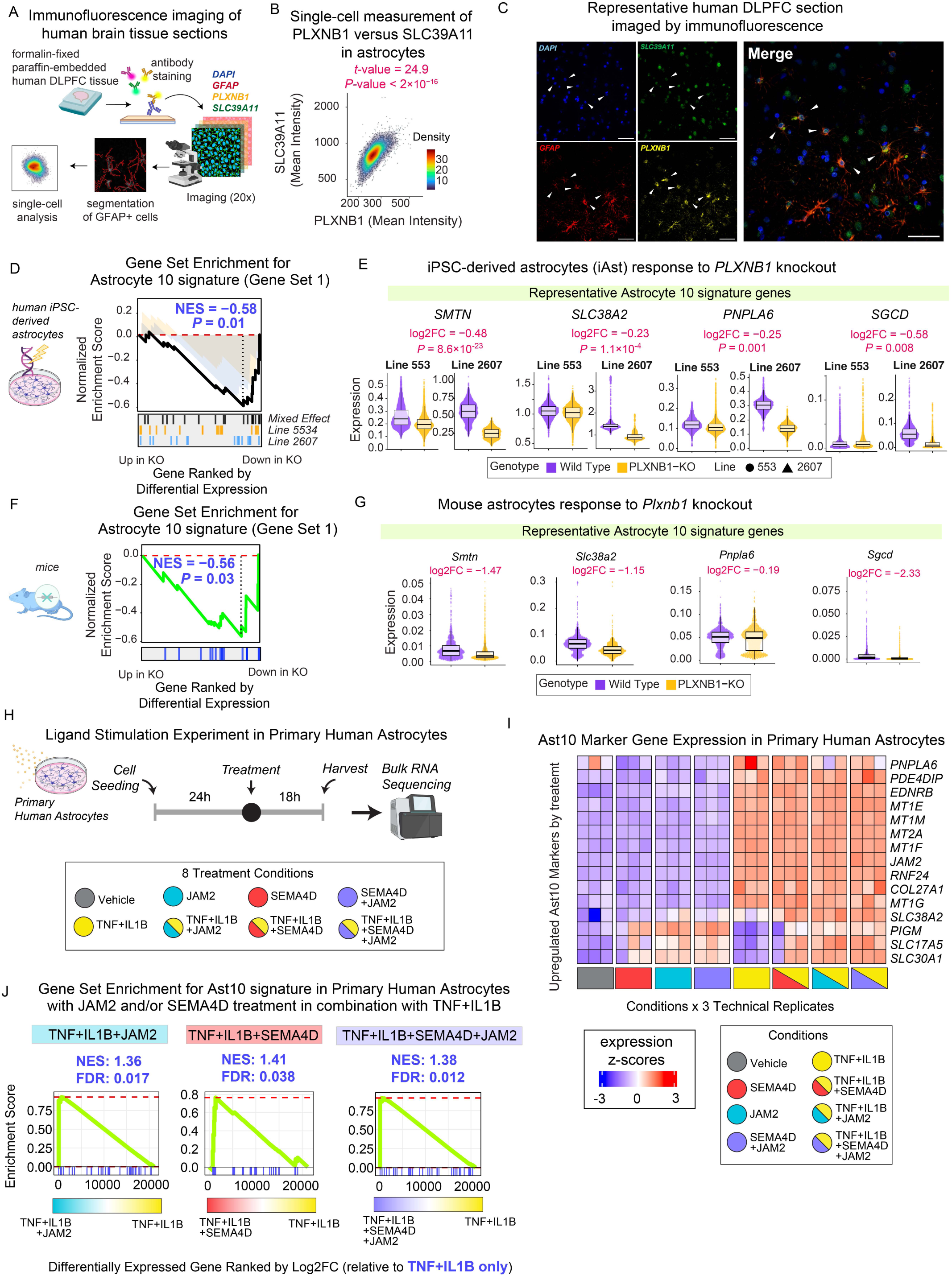
Validation of JAM2 and SEMA4D/PLXNB1 signaling in the emergence of Ast10. **(A)** Schematic of the immunofluorescence approach to validate PLXNB1 coexpression with the Ast10 marker SLC39A11 in human dorsal lateral prefrontal cortex (DLPFC) tissue from an AD donor. The FFPE section was stained for DAPI (nuclei), GFAP (astrocytes), SLC39A11 (Ast10 marker), and PLXNB1. Forty 20x images were systematically collected across cortical layers. Astrocytes were identified via DAPI and GFAP segmentation, and mixed-effect modeling assessed single-cell covariation between PLXNB1 and SLC39A11. **(B)** Single-cell density plot showing SLC39A11 versus PLXNB1 expression, quantified by mean fluorescence intensity. Each point represents a GFAP+ astrocyte across 40 images. Effect size and significance for PLXNB1 predicting SLC39A11 levels were determined using a mixed-effect model (see Methods). **(C)** Representative tissue section showing DAPI, GFAP, PLXNB1 and SLC39A11 staining. Astrocytes coexpressing high levels of PLXNB1 and Ast10 marker SLC39A11 (GFAP^high^SLC39A11^high^PLXNB1^high^ cells) are marked by white arrows. Scale bars: 50 μm. **(D)** Gene set enrichment analysis (GSEA) of the Ast10 signature (Gene Set 1) in hiPSC-derived astrocytes (iAst) comparing *PLXNB1*-knockout (KO) versus wild-type cells. Genes were ranked by differential expression (Wald statistic) using Nebula (see Methods), from those upregulated in KO (left) to those downregulated in KO (right). The running enrichment score and corresponding *P* value were derived from a mixed-effects model summarizing differential expression across both lines. Horizontal bars indicate the positions of Ast10 signature genes in the ranked lists from the mixed-effects model and from individual lines 553 and 2607. NES, normalized enrichment score. **(E)** Representative Ast10 signature gene expression at the single-cell level measured in iAst with *PLXNB1* knockout (*PLXNB1*-KO) or without (Wild Type) across two independent donor lines (line 553, 2607). Log2 fold-change and *P* values shown were estimated from the mixed effect differential expression analysis using Nebula. **(F)** GSEA of the Ast10 signature (Gene Set 1) in mouse astrocytes comparing *Plxnb1*-knockout (KO) versus wild-type cells. Genes were ranked by differential expression (Wald statistic) estimated using Nebula (see Methods). The running enrichment score and corresponding *P* value were calculated from the ranked gene list. Horizontal bars indicate the positions of Ast10 signature genes within the ranked list. NES, normalized enrichment score. Sample size: *n* = 1 per genotype. **(G)** Representative Ast10 signature gene expression at the single-cell level measured in mouse astrocytes with *Plxnb1* knockout (*Plxnb1*-KO) or without (Wild Type). Log2 fold-change shown were estimated from the differential expression analysis using Nebula. **(H)** Ligand stimulation experiment in primary human astrocytes derived from a donor to validate prioritized Ast10-predictive ligands, JAM2, SEMA4D and their combination (SEMA4D+JAM2), in the presence or absence of inflammatory cytokines (TNF+IL1B). Cells were seeded for 24 hours, treated for 18 hours, then harvested for bulk RNA-seq. **(I)** Z-scored expression of Ast10 marker genes differentially upregulated by any treatment conditions. Each column represents a technical replicate of each condition. **(J)** Gene set enrichment analysis of the Ast10 signature in primary human astrocytes treated with TNF + IL1B alone or together with JAM2, SEMA4D, or JAM2 + SEMA4D. Genes were ranked relative to TNF + IL1B alone using a signed composite statistic, calculated as the Wald statistic × -log10(*P*-value). Normalized enrichment scores (NES) and false discovery rates (FDR) are shown.

### Altered PLXNB1 expression reduces Ast10 signature in human iPSC-derived astrocytes and in an in vivo mouse model

To validate the role of *PLXNB1* in regulating the Ast10 signature in a human cellular model, we accessed previous validated^64^ human induced pluripotent stem cell (hiPSC) lines from two control donors (CD1: 553 and CD2: 2607), in which CRISPR/Cas9 editing introduced frameshift deletions within the coding sequence of the *PLXNB1* exon 3^64^, and compared them with their respective isogenic controls (**Extended Data Figure 9B)**. Successful editing was confirmed by Sanger DNA sequencing^64^ (See Methods). Both isogenic pairs of hiPSCs were differentiated into astrocytes by viral induction of *SOX9* and *NFIB* expression^65^ (**Extended Data Figure 9B**; See Methods). We then performed scRNA-seq to profile transcriptome-wide changes in the hiPSC-derived astrocytes (iAst) due to *PLXNB1* perturbation.

We first evaluated the similarity between the Ast10 transcriptional program in the *baseline* iAst models (iAst Wild Type) and that observed in postmortem human astrocytes. To achieve this, we performed a gene-gene correlation analysis, comparing the averaged coexpression patterns of Ast10 markers in the baseline iAst models with those in the snRNAseq dataset (**Extended Data Figure 9C**). In the baseline iAst models, Ast10 markers were grouped into two distinct coexpression gene modules: Gene Set 1 included genes mainly related to ion transport (e.g., *SLC38A2, SLC30A1*) and extracellular matrix or cytoskeleton remodeling (e.g., *SMTN, SGCD*); whereas Gene Set 2 mainly included metallothionein genes (e.g., *MT1E*). However, in postmortem snRNAseq data, these two gene modules are strongly correlated with one other (**Extended Data Figure 9C**). This suggests that what appears to be a single Ast10 signature in postmortem human astrocytes is actually composed of two distinct transcriptional programs whose regulations can be disconnected *in vitro*. Thus, we evaluated the two components of the Ast10 signature separately in subsequent analyses.

At the gene set level, Gene Set 1 of the Ast10 transcriptional program showed significant enrichment among genes that are downregulated upon *PLXNB1* knockout (Normalized Enrichment Score, NES= -0.58 *P* = 0.01), with concordant trends observed in both donor lines as expected (**Figure 5D).** Representative Gene Set 1 members, such as *SMTN* (log2 fold-change, log2FC = -0.48), *SLC38A2* (log2FC = -0.23), *PNPLA6* (log2FC = -0.25), and *SGCD* (log2FC = -0.58), were consistently downregulated in *PLXNB1*-KO iAst across both donor lines (**Figure 5E)**. By contrast, Gene Set 2, mainly composed of metallothionein genes, showed no significant changes following *PLXNB1* knockout (NES = 0.58, *P* = 0.08, **Extended Data Figure 9E**).

Since the human cellular model does not fully capture the cellular context in the brain, we also tested the regulatory role of *PLXNB1* for Ast10 signature *in vivo*, by repurposing a published single-cell RNAseq dataset collected from mouse cortical tissues to compare reference (wild-type) and *Plxnb1* knock-out (*Plxnb1*^−/-^) animals^64^. To computationally identify the astrocytes from this dataset, we annotated individual cells by their major cell types, using our snRNAseq Discovery data as reference (See Methods). Consistent with our human *PLXNB1-KO* iAst findings, *Plxnb1* knockout significantly decreased astrocyte expression of the Ast10 Gene Set 1 (NES = -0.56, *P* = 0.03; **Figure 5F**) but not Gene Set 2 (NES = 0.86, *P* = 0.002, **Extended Data Figure 9G**). Representative genes decreased in *PLXNB1-KO* iAst showed a consistent downregulation in *Plxnb1*^−/-^ murine astrocytes (**Figure 5G**): *Smtn* (log2FC = -1.47), *Slc38a2* (log2FC = -1.15), *Pnpla6* (log2FC = -0.19), and *Sgcd* (log2FC = -2.33). These results validate the regulatory role of astrocytic *PLXNB1* in maintaining the Ast10 transcriptional state *in vivo*.

Altogether, these findings demonstrate that *PLXNB1* drives at least a part of the main transcriptomic signature of Ast10 consistently across two model systems; it is likely that the complete signature only emerges with the activation of several receptors and the convergence of multiple different signals. We therefore evaluated the effect of a second ligand, JAM2, in addition to that of SEMA4D which works through PLXNB1.

### JAM2 and SEMA4D augment the Ast10 transcriptional response in a reactive cytokine context

Having validated our ability to reduce Ast10 by genetically depleting the Ast10-driving receptor *PLXNB1*, we next tested whether the Ast10 transcriptional signature could be induced *in vitro* by treating primary human astrocytes with the top-ranked ligands *SEMA4D*, *JAM2*, or their combination. While JAM2 is not known to directly interact with SEMA4D, these signaling molecules converge on RhoA and cellular remodeling pathways. Therefore, we also tested their combination which may exert synergistic or redundant effects on Ast10 induction. In addition, we profiled their effect in an experimental paradigm simulating reactive astrocytes using a cytokine cocktail of TNF (5 ng/ml) and IL1B (1 ng/ml) (TNF+IL1B) (**Figure 5H**)^66,67^. The latter condition addresses the possibility that the effect of our prioritized ligands is context-dependent^66,68^.

Primary human astrocyte cells were harvested at 18 h after stimulation^66^; RNA was extracted and underwent bulk RNA-sequencing. Although treatment with SEMA4D and/or JAM2 alone modestly upregulated several Ast10 markers (**Figure 5I**), gene set enrichment analysis of differentially expressed genes relative to vehicle revealed no significant enrichment of the entire Ast10 program (FDR =0.07-0.44; **Extended Data Figure 9H**). TNF+IL1B treatment alone induced a broader set of Ast10 signature genes, particularly metallothionein genes (**Figure 5I, Extended Data Figure 9I**) but likewise did not significantly enrich the Ast10 program as a whole (FDR = 0.08; **Extended Data Figure 9H**). In contrast, co-treatment with TNF+IL1B and SEMA4D, JAM2, or their combination significantly amplified the Ast10 transcriptional program relative to TNF+IL1B alone (JAM2: *NES* = 1.36, *FDR* = 0.017; SEMA4D: *NES* = 1.41, *FDR* = 0.038; JAM2 + SEMA4D: *NES* = 1.38, *FDR* = 0.012; **Figure 5J**). Together, these results indicate that inflammatory signaling alone induces partial Ast10-like features as part of a generic reactive astrocyte response, whereas JAM2 and SEMA4D provide additional, context-dependent cues that selectively amplify the Ast10 transcriptional program.

### Ast10 has a pathology-independent effect on cognitive decline and may exert its effect through presynaptic dysfunction

The significant, robust association between Ast10 frequency and cognitive decline across multiple independent studies and our prior structural equation modeling suggest an important, potentially causal role of Ast10 in AD-related cognitive decline^1^. However, while AD pathology is a major contributor to aging-related cognitive decline, many other factors contribute to this process, and more than half of the variance in cognitive decline remains unexplained after all known risk factors are accounted for^69^. To understand the extent to which other factors may explain the association between Ast10 frequency and cognitive decline, we deployed a linear regression model that includes all known aging-related proteinopathies (amyloid, tau, Lewy bodies, TAR DNA-binding protein 43), hippocampal sclerosis, multiple measures of neurovascular pathology, and other factors (such as the level of education and age at death) that are known to be associated with cognitive decline (**Extended Data Figure 10A**). We then added an additional term for the frequency of Ast10 (**Figure 6A**) which remains significantly associated (*P* = 3.1 × 10^−6^) with the slope of cognitive decline after all listed variables are accounted for. Thus, we can say that, beyond its role in AD, Ast10 captures the consequences of other, as yet undiscovered, processes that contribute to aging-related cognitive decline.

**Figure 6.**
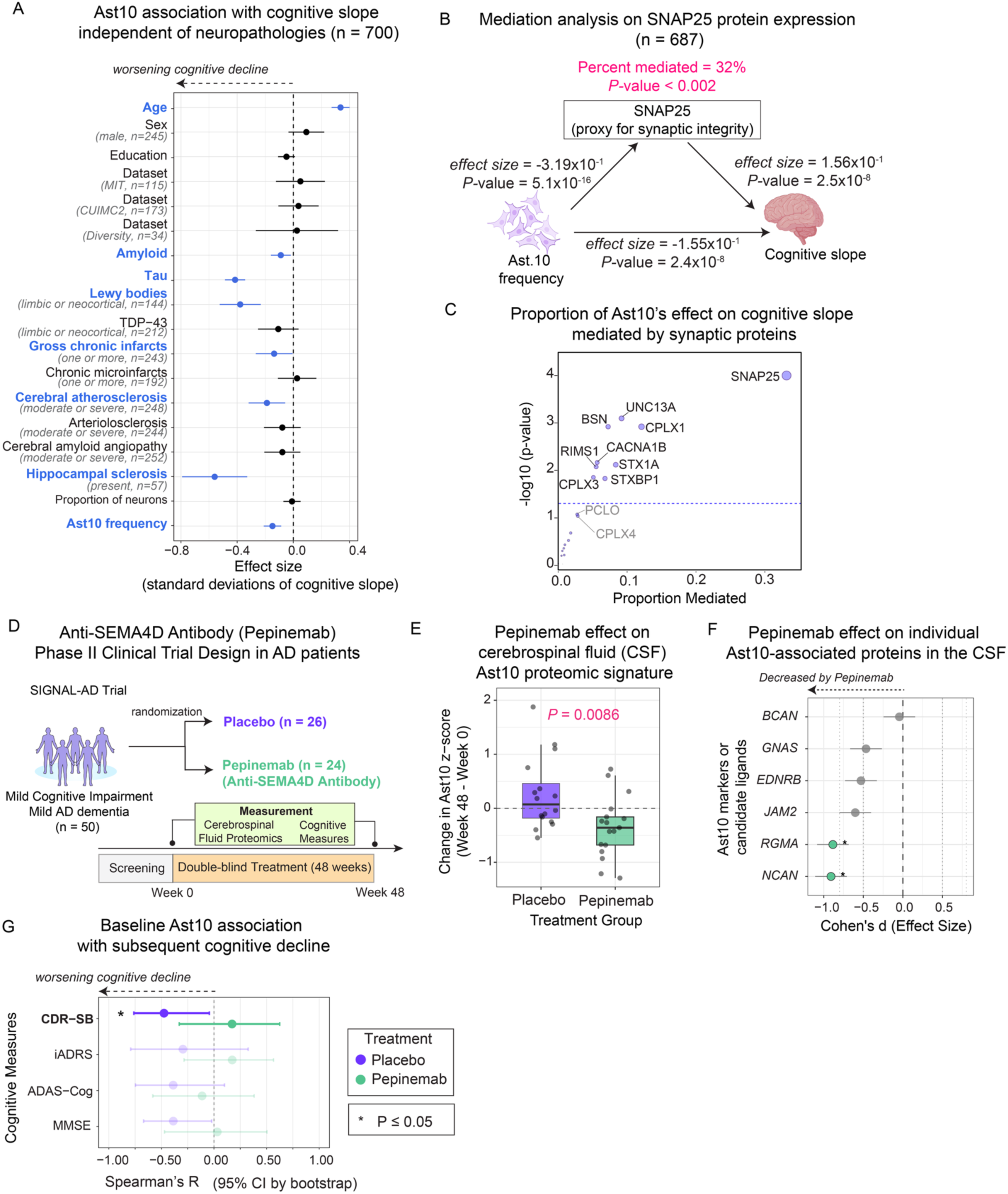
Ast10 is associated with cognitive decline through partial synaptic protein mediation and is reflected in a pepinemab-responsive CSF signature in an AD clinical trial. **(A)** Multivariate regression model predicting cognitive decline based on Ast10 frequency, neuropathologies, and demographics. Effect sizes (dots) and 95% confidence intervals (error bars) are shown; significant variables (blue) are highlighted. A total of *n* = 700 samples across all snRNAseq datasets (Discovery + Replication) with complete measurements of all variables were analyzed. **(B)** Mediation analysis testing SNAP25 protein expression as a mediator of the Ast10-cognitive decline association in 687 ROSMAP DLPFC samples. SNAP25 significantly (*P* < 0.002) mediated 32% of the association. *P* value derived from 5000 bootstrap iterations. **(C)** Mediation analysis testing other synaptic proteins of the SNARE complex as a mediator of the Ast10-cognitive decline association. *P* values derived from 5000 bootstrap iterations. Dot size scales with proportion mediated. **(D)** Schematic of the Pepinemab SIGNAL-AD Trial, which enrolled 50 participants (confirmed amyloid positive, with mild cognitive impairment) for a randomized, double-blind treatment of either placebo or Pepinemab for 48 weeks. Outcome assessments included pre- and post-treatment Olink proteomics profiling of the cerebrospinal fluid (CSF) and cognitive measures. **(E)** Comparison of the change in Ast10 proteomic signature (Ast10-zscore) between treatment groups after 48 weeks (n= 16 placebo and n=17 pepinemab). Ast10-zscore was derived from aggregating cerebrospinal fluid proteomic measurements of six Ast10-predicting ligands or markers that were available on the Olink panel. (see Methods). These six markers include: BCAN, GNAS, EDNRB, JAM2, RGMA, NCAN. *P*-value: two-sided unpaired Student’s t-test. **(F)** Forest plot showing Cohen’s d effect sizes for the changes in individual Ast10-predicting protein (Week 48 minus Week 0), comparing placebo versus pepinemab-treated groups. Each point represents the standardized mean difference between treatment groups for a single protein. Error bars indicating 95% confidence intervals. Negative values indicate greater decreases (or smaller increase) in Pepinemab relative to Placebo control. Green points: FDR < 0.05 (Benjamini-Hochberg correction); gray points: not significant. Significance: *FDR < 0.05. **(G)** Forest plot showing Spearman’s correlations between the baseline Ast10 proteomic signature and cognitive change at 48 weeks, across four cognitive measures in the placebo versus pepinemab-treated groups. Negative correlation coefficient indicates greater cognitive decline. Error bars indicating 95% confidence intervals. \**P* ≤ 0.05.

To begin probing the mechanism linking Ast10 to cognition, we hypothesized that Ast10 may reduce synapse integrity, ultimately leading to cognitive decline. Synaptosomal-associated protein 25 (*SNAP25*) is a core component of the presynaptic vesicle fusion machinery and is strongly associated with cognitive decline in ROSMAP. SNAP25 protein level has previously been used as a proxy for synaptic integrity in the aging brain^70,71^. Leveraging matching bulk DLPFC proteomic data from a subset of the ROSMAP participants in our analyses (n = 687 have snRNAseq and proteomic data from the same brain region, DLPFC), we first tested whether *SNAP25* protein expression mediates the effect of Ast10 on cognitive decline. Higher levels of *SNAP25* protein, as previously reported^72^, is associated with slower cognitive decline (*P* = 2.5 × 10^−8^). Since Ast10 frequency is significantly associated with both cognitive decline (*P* = 2.4 × 10^−8^) and SNAP25 expression (*P* = 5.1 × 10^−16^) in these ROSMAP participants (**Figure 6B**), we asked whether the association between Ast10 frequency and SNAP25 expression was better supported in the model where SNAP25 is statistically downstream (i.e., the mediator) or upstream of Ast10 (i.e. the exposure). Direction dependence analysis shows that SNAP25 is better explained as a mediator (**Extended Data Figure 10B**). We then test for mediation, which shows that SNAP25 mediates 32% (*P* < 0.002) of the association between Ast10 frequency and cognitive decline (**Figure 6B**).

To assess whether this effect extends beyond a single synaptic marker, we expanded this analysis to additional components of the presynaptic neurotransmitter release machinery, including core SNARE proteins (STX1A, STX1B, VAMP1, VAMP2) and their regulators (CPLX1-4). Several of these proteins also showed significant mediation effects, with SNAP25 exhibiting the largest estimated proportions mediated, followed by UNC13A, CPLX1 and STX1A (**Figure 6C, Extended Data Figure 10B**), suggesting that disrupted presynaptic vesicle fusion may be one mechanism linking Ast10 enrichment to cognitive decline. Together, these results support a model in which Ast10 cells may contribute to impaired synaptic release function and subsequent neuronal dysfunction. This hypothesis connects with several of the characteristics of the Ast10 signature, such as genes involved in extracellular matrix remodeling (e.g., *JAM2*, *EFEMP1, NCAN*, **Figure 3C**), which have been implicated in perisynaptic nets that contribute to the maintenance of synapses, and PLXNB1-SEMA4D signaling, which has been reported to regulate inhibitory synapses^73^.

### Ast10 biomarker signature in cerebrospinal fluid is reduced by anti-SEMA4D blocking antibody in patients with early-stage AD

To validate our predicted ligands in a patient context and assess the therapeutic relevance of Ast10, we next analyzed cerebrospinal fluid (CSF) proteomic profiles (Olink Explore 5000 panel) from a clinical trial (SIGNAL-AD, NCT04381468) of pepinemab, a SEMA4D blocking antibody. In this randomized, double-blind, placebo-controlled clinical trial, 50 individuals (who fulfilled entrance criteria for being positive for amyloid proteinopathy using PET scan and having signs of mild cognitive impairment or mild AD), (see **Methods**) were treated with either pepinemab or placebo for 48 weeks by intravenous infusion between July 2021 and June 2024. A CSF sample and cognitive measures were collected at baseline and at end of the treatment period of 48 weeks. (**Figure 6D**).

Using this dataset, we first asked whether Ast10-associated proteins detectable in CSF were modulated by pepinemab treatment. Six Ast10-related proteins were measured by the Olink panel, including top ligands predictive of Ast10 enrichment (JAM2, BCAN, NCAN, GNAS, RGMA) and Ast10-upregulated signature genes (JAM2, EDNRB, BCAN). For each participant, we computed an Ast10 CSF summary score based on these proteins (see **Methods**) and compared longitudinal changes between treatment groups. Importantly, all analyses were performed blinded to treatment assignment. Ast10 CSF signature levels were significantly reduced after 48 weeks of pepinemab treatment and to a greater extent than in the placebo group (t-test, *P* = 0.0086; **Figure 6E**; see **Extended Figure 10B** for individual patient trajectories). Among these six Ast10-associated proteins, RGMA and NCAN individually showed significantly greater reduction in the pepinemab group than in placebo (**Figure 6F)**.

Next, we examined whether this CSF-derived Ast10 proteomic signature was associated with cognitive decline, and whether pepinemab treatment altered this relationship. Cognitive decline was quantified as the average longitudinal change in each of four clinical measures (CDR-SB, iADRS, ADAS-Cog, MMSE, see **Methods** for metrics detail). In the placebo group, higher baseline Ast10 CSF scores were associated with greater subsequent cognitive impairment (**Figure 6G**). Notably, worsening in CDR-SB score is significantly associated with baseline Ast10 proteomic signature in the placebo group (Spearman R = -0.48, *P* = 0.05, n=17), while trends were observed in other cognitive measures: ADAS-Cog13 (Spearman R = -0.39, *P* = 0.09, n=20) and MMSE (Spearman R = -0.39, *P* = 0.075, n=22). In contrast, these associations were not observed in the pepinemab-treated group **Figure 6G**. Together, these findings suggest that SEMA4D blockade with pepinemab reduces an Ast10-associated proteomic signature in the CSF and disrupts the association of Ast10 with cognitive decline, supporting a model in which SEMA4D-dependent Ast10 expansion contributes to cognitive decline in AD.

## Discussion

To date, an increasing number of disease-associated cellular states of every major cell type has been reported to be involved in different aspects of AD^1,3–11^. However, there remains a major gap in our knowledge when it comes to the origins of these cell subpopulations and their inter-cellular coordination over the course of brain aging and disease progression. Our work and other reports have proposed that these disease-associated cellular states often appear in concert^1,2,16,55,64,74–78^, suggesting that their origins are likely orchestrated by cell-cell interactions or through responses to shared microenvironmental signals.

In this study, we substantially expanded the human DLPFC single-nucleus **Resource** for aging brains and performed a staged meta-analysis across four independent datasets comprising 867 brains. This analysis confirmed the robustness of the core glial cell-state architecture associated with AD-related traits and identified an uncommon microglial state (Mic16) and the vascular unit such as the endothelial cell subtype End3 (**Figure 1B**). These analyses also present an initial assessment suggesting that the findings may be relevant among non-European individuals, although much more work is needed to fully address this issue. Here, we elected to pursue the characterization of Ast10 as an exemplar, developing an approach to identify the intercellular signals that lead to the differentiation of a prioritized cell state.

After analyzing 4.2 million nuclei collected from 867 brains, we identified an *SLC38A2^high^SMTN^high^CACNA1D^high^*astrocyte state, Ast10, as one of the cell subpopulations (among the 91 tested) most strongly associated with cognitive decline, the ultimate target for therapeutic development as many individuals with evidence of AD proteinopathy are cognitively unimpaired^3,79^: they are resilient. In our earlier work, Ast10 defined the terminus of a trajectory of brain aging that leads to AD^1^, suggesting that blocking its development may help to prevent dementia in the setting of amyloid and tau pathology. Ast10 is characterized by a distinct transcriptional signature associated with intracellular ion transport, extracellular matrix organization and oxidative stress. Key marker genes in this signature include ion transporters such as *SLC38A2*, *SLC39A11* and *CACNA1D*, cytoskeleton components such as *SMTN* and *SGCD*, as well as metallothionein genes like *MT1F* and *MT1G*. This signature stands apart from the more commonly described Disease-Associated Astrocyte (DAA) state, which typically exhibits an acute inflammatory response profile, including *GFAP*, *SERPINA3*, and *OSMR*^8,16,63^. Although the origins and specific functions of the Ast10 state have not been studied, a partially overlapping astrocyte signature (e.g. *SLC38A2*, *MT1G*, *CACNA1D*) was upregulated in AD samples from a smaller, independent study^5^. Another independent snRNAseq study that sampled the spatiotemporal progression of AD across 32 participants highlighted astMet, a late-stage AD-associated astrocyte subset^80^. Notably, astMet includes both stress-response markers enriched in Ast9 (e.g, *BAG3, HSPB1*) and metallothionein markers (e.g., MT1G, MT1E) that mark Ast10 (**Figure 1D**), suggesting that astMet may reflect a broader reactive astrocyte program encompassing multiple states resolved in our atlas. Together, these studies further support an important role of astrocytes in AD, and our large sample size and deeply phenotyped participants have enabled us to resolve the role of the rare Ast10 subtype in the slope of cognitive decline, going beyond evaluations of pathologic features and summary diagnoses.

Focusing on Ast10 as an important but understudied cell state, we performed a systematic analysis to identify the microglial or astrocytic signals that may drive the emergence of the Ast10 state. We designed our computational framework to provide convergent, multi-scale support for signaling hypotheses underlying Ast10 emergence: (1) NicheNet uses literature-curated ligand-receptor-target relationships to prioritize ligands that could plausibly regulate the Ast10 transcriptional program. Such an approach is distinct from common cell-cell communication inference approaches which typically only prioritize ligand-receptor interactions based on the cell-type/subtype specificity of expression alone^81–83^; (2) Multivariate modeling with PLSR further confirmed that expression of the prioritized ligands in specific sender populations predict donor-level variation in Ast10 frequency. We have also illustrated how PLSR can yield specific hypotheses that can be validated experimentally (i.e. *PLXNB1, SEMA4D, JAM2*); (3) DREMI addresses a different question at single-cell resolution by quantifying whether variation in downstream pathway components of the ligands is coordinated with cell-to-cell variation in the Ast10 transcriptional program across astrocytes, revealing that this program exists along a continuum rather than as a strictly discrete state. By interrogating distinct features of the snRNAseq data at different biological scales while converging on similar candidates, our approach prioritizes robust ligands that are unlikely artifacts of any single computational method. Such an approach can be repurposed to explore the paracrine regulation of our other cell subpopulations that are associated with AD traits (**Figure 1B**). Finally, we validated our proposed prioritized regulators in two independent datasets, CUIMC2 and MIT (**Figure 3F,G**).

A key validation effort in this study is the use of two complementary spatial transcriptomic approaches (imaging-based and sequencing-based approaches) to provide direct, independent evidence for the role of prioritized ligands in regulating the Ast10 signature profiled in human tissue while preserving its spatial context. We first analyzed the sequencing-based (Visium) spatial transcriptomic data from 32 DLPFC tissue sections obtained from 17 donors, the majority of whom exhibited advanced AD pathology. The prioritized ligands in the local neighborhood were significantly and specifically predictive of the Ast10 signature at the center of the neighborhood, compared to other astrocytic states, providing independent spatial support for their potential role in driving the Ast10 state in AD brains (**Figure 4A-D**). Furthermore, the transcriptome-wide nature of the sequencing-based spatial profiling enabled an unbiased assessment of the prioritized ligand set’s significance compared against all ligands detected in the tissue (**Extended Figure 7E**). Interestingly, this analysis also identified additional ligand sets with high importance (**Extended Figure 7E**), potentially expressed in other cell types not included in our initial NicheNet analysis, which warrants future investigation. While these analyses establish spatial association at the neighborhood level, they do not resolve whether these signaling relationships are indeed engaging with Ast10 cells at the single-cell level or how they are organized *in situ*. We therefore generated and analyzed a MERSCOPE Ultra dataset measuring 447 genes using V2 chemistry and the V2 gene probe panel across 19 DLPFC sections from 15 ROSMAP donors with matched donor-level snRNA-seq profiles, spanning both previously defined brain aging trajectories (**Figure 1A)**. To our knowledge, this represents one of the largest imaging-based, single-cell-resolution spatial transcriptomic datasets of the adult human DLPFC with matched snRNA-seq, as well as one of the first reported applications of MERSCOPE Ultra V2 chemistry to postmortem human brain tissue. This resource enabled us to map Ast10 astrocytes, candidate ligands, and candidate receptors in the same tissue sections at cellular resolution.

MERFISH analysis revealed that Ast10 astrocytes are not uniformly distributed across tissue, but instead form spatially organized clusters, particularly within grey matter, suggesting that Ast10 emergence is shaped by localized microenvironmental signals rather than uniform tissue-wide cues. Despite this focal enrichment, we found that Ast10 frequencies were strongly correlated between grey and white matter across individuals, suggesting that global tissue-level factors establish overall Ast10 burden while local signaling niches govern focal accumulation. Together, these observations highlight the need for future studies to test whether focal Ast10 clustering is spatially associated with local neuropathological features, and to determine whether donor-level and local influences arise from distinct mechanisms or represent manifestations of a shared underlying process. At the single-cell level, spatially lagged modeling confirmed that predicted Ast10-driving ligands such as *SEMA4D* and *JAM2* are enriched in the immediate neighborhoods of Ast10-high astrocytes, whereas ligands predicted to negatively associate with Ast10, such as *EFEMP1*, *SEMA4B*, and *FGF1*, are locally depleted. Moreover, top receptors prioritized by the PLSR model, including *PLXNB1* and *JAM2*, showed robust positive associations with Ast10 scores within the same cell, providing direct *in situ* confirmation that these receptors are co-expressed with the Ast10 program rather than indirectly correlated at the donor level. The spatial enrichment of *JAM2* around Ast10 cells and its co-expression with the Ast10 program within the target astrocytes suggest that *JAM2* may represent both a component of the intrinsic Ast10 transcriptional program and a candidate juxtacrine signal that could influence an astrocyte’s transition to the Ast10 state. For *PLXNB1*, we went one step further and offered a third tissue-based assessment, this time at the protein level: PLXNB1 and a predicted single gene proxy for the Ast10 signature, SLC39A11, are correlated in individual astrocytes at the protein level (**Figure 5A-C**). Altogether, these findings demonstrate that Ast10 is embedded within spatially structured signaling microenvironments that are consistent with the ligand-receptor interactions inferred from snRNAseq analyses. These spatial analyses support local, non-cell-autonomous associations between prioritized ligand-receptor axes and the Ast10 program and identify specific signaling relationships whose molecular engagement can now be tested experimentally. Methodologically, this study highlights the power of integrating multi-modal data with rigorous statistical methodology and downstream multi-modal validation. First identifying disease-relevant cell subpopulations and their signaling regulators through well-powered single-cell transcriptomics and then situating them within the tissue context using spatial approaches.

To move beyond correlative inference, we leveraged targeted perturbations to mechanistically test our hypothesis that our predicted ligands lead astrocytes to adopt an Ast10 state. We prioritized two of them, JAM2 and SEMA4D, given their strong and consistent prioritization across single-nucleus transcriptomic and spatial analyses. Using primary human astrocytes, both of these ligands, are validated as proteins that lead to the engagement of the Ast10 state in astrocytes. Interestingly, their individual effects are incomplete at baseline but significant in the proper context: that of an inflammatory response (TNF+IL1B)^68^, highlighting the contextual nature of the signaling that we have uncovered in our modeling of snRNAseq data. Further, these observations suggest that Ast10 is distinct from a generalized astrocyte reactivity state induced by the TNF+IL1B paradigm that has been studied previously^16,66,68^: in this framework, JAM2 and SEMA4D act as context-dependent modulators that bias reactive astrocytes toward a more specialized Ast10 state that is related to AD and other processes leading to synaptic loss and cognitive decline.

Genetic ablation of *PLXNB1, the SEMA4D* receptor, in mice and iAst further validates our prioritized signals: both model systems show a reduced Ast10 signature (**Figure 5D-G**). *PLXNB1* has been associated with various central nervous systems disease contexts including AD^64,68,73^. We previously identified *PLXNB1* as a key node of a gene module from bulk neocortical RNAseq data that is significantly associated with aging-related cognitive decline in the ROSMAP cohort^84^. Another recent study suggests that *PLXNB1* plays a critical role in regulating cell-cell distancing of microglia and astrocytes around amyloid plaques^64^. Our findings therefore elaborate our understanding of *PLXNB1’s* role, implicating it also in the more distal processes of synaptic loss and cognitive decline and not simply the accumulation of amyloid proteinopathy. Further, it places PLXNB1 within a larger framework in which JAM2 receptors and other convergent signals lead astrocytes to a specialized state that is involved in synaptic response to AD pathology and other processes (**Figure 6B,C**).

Many of the significant predictors for Ast10 from the ligand PLSR model and spatial analysis were associated with axon guidance, synapse assembly, or synaptic function, including *SEMA4D, RGMA, NRXN1* and *SEMA4B.* This enrichment of synapse-related molecules aligns with the potential role of Ast10 in modulating synaptic integrity and is further supported by our mediation analysis with *SNAP25* and other SNARE proteins (**Figure 6B,C**). Additionally, our results also revealed a specific extracellular matrix (ECM) profile associated with the Ast10 state, adding to existing evidence that suggest an important role of astrocyte-ECM interactions in neurodegeneration^85,86^. In particular, our PLSR model shows that the enrichment of Ast10 in the cortex is associated with increased levels of the lectican family of chondroitin sulfate proteoglycans, such as neurocan (*NCAN*) and, to a lesser extent, with brevican (*BCAN*) (**Figure 3C**), which are key components of the perineuronal ECM^87^. In contrast, fibronectin (*FN1*) that makes up the basement membrane ECM surrounding the vasculature^88^ is not significantly predictive of Ast10 frequency and trends towards Ast10 depletion (**Figure 3C**). These results suggest that Ast10 may preferentially reside around neurons, possibly interacting with the perisynaptic ECM to alter synaptic functions. Similar spatial analysis with MERFISH data as described in this study will be needed to reveal whether Ast10 spatially localize with interacting neuronal subsets to test this hypothesis. The exact role of such lectican-rich ECM microenvironment in AD remains unclear. While lectican may serve as a protective barrier for neurons^89^, it may also mediate neuroinflammatory responses^88,90^ or inhibit synaptic plasticity^91,92^. These seemingly opposite roles of lecticans likely depend on their versatile post-translational modifications. Since astrocytes are a primary source of lecticans, future studies should investigate the potential mutual influence between Ast10 and surrounding ECM, and how such interactions may impact synaptic function in AD.

Altogether, these results suggest that the increased frequency of Ast10 may lead to disruption of the established peri-synaptic nets and subsequent loss of synapses by neurons in AD. However, we also show that Ast10 explains a significant amount of the variance in aging-related cognitive decline that is not explained by known risk factors such as AD proteinopathies. Thus, Ast10 may also play a key role in other, as yet unknown, processes that contribute to loss of cognition in older age (**Figure 6A**).

Our analyses of the pepinemab trial data are interesting in several ways: (1) it demonstrates that the Ast10 signature is measurable in CSF and may be responsive to therapeutic intervention, opening up its use a relevant biomarker for AD and perhaps other conditions. (2) It provides key validation that blocking SEMA4D *in vivo* in humans reduces the presence of the Ast10 state and may be related to slower cognitive decline. This analysis provides an interesting mechanistic insight into pepinemab which is being developed for AD and Huntington Disease based on an hypothesis that inhibiting SEMA4D can preserve or restore neuronal function by modulating astrocyte reactivity^93^. Our results therefore support Ast10 as a potential mediator of pepinemab’s mechanism of action.

## Limitations of the study

Several limitations should be considered when interpreting these findings. Our computational analyses of cross-sectional data from autopsy specimens are inherently correlative and cannot, on their own, establish causality. However, convergence of evidence across donor-level, single-cell, and spatial analyses helps generate mechanistic hypotheses, prioritizing ligand candidates (from hundreds of ligands to a dozen) across complementary biological scales. We further provide mechanistic support through perturbation experiments showing that selected JAM2, SEMA4D and PLXNB1 regulate features of the Ast10 program in model systems, while the pepinemab analyses offer orthogonal human evidence that modulation of the SEMA4D pathway is accompanied by changes in an Ast10-related CSF signature. The CSF signature is best interpreted as a protein signature that captures the broader molecular environment linked to the Ast10 state; we cannot attribute the production of the proteins in the Ast10 signature uniquely to astrocytes. These proteins may come from both astrocyte-intrinsic and/or other cellular sources. However, the CSF protein signature’s reduction following SEMA4D blockade provides human *in vivo* evidence that Ast10-associated signaling is pharmacologically modifiable. Although this study prioritized microglia and astrocytes, it is likely that other cell types expressing the same ligands (e.g., vascular endothelial cells also express *JAM2*) or other ligands may contribute to Ast10 emergence. Our study thus provides a systematic, scalable framework that can be generalized to studying other cell type/subtype contributors to disease-associated cell states; in future work, we will expand the approach to all cell types. Overall, these results strengthen the case for Ast10-related signaling as a biologically meaningful and potentially actionable pathway. Future longitudinal and perturbational studies in human-relevant systems will be needed to determine how directly these other prioritized ligand pathways regulate Ast10 *in vivo* and to define the causal mechanisms and functions linking Ast10 to cognition, as suggested by the mediation analyses. Altogether, our study highlights that the Ast10 state may be a key nexus for multiple different pathophysiological processes that lead to synaptic loss and cognitive dysfunction, making it and its regulators promising targets for therapeutic development to enhance resilience not just to AD but also other dementing pathophysiologies.

## Methods

### Participants

Human molecular profiles and clinicopathological traits used in this study are measured from participants from the following five community-based cohort studies of aging and dementia. All five studies are conducted by the Rush Alzheimer’s Disease Center (RADC) and share a large common core of data collected by the same team^26^. All five were approved by an Institutional Review Board of Rush University Medical Center. All participants enrolled without known dementia and agreed to annual clinical evaluation. All participants signed informed and repository consents and brain donors signed an Anatomic Gift Act. Below are summaries of the study designs and demographics:

#### Religious Orders Study (ROS)^14^

The Religious Orders Study (ROS) is a longitudinal epidemiologic clinical-pathologic cohort study focused on aging and Alzheimer’s disease. Participants include older Catholic nuns, priests, and brothers from diverse groups across the US. Enrollment began in 1994, with participants agreeing to annual clinical evaluation, cognitive testing, and a subset have annual blood draw. All participants agree to brain donation and sign an Anatomic Gift Act.

#### Rush Memory and Aging Project (MAP)^15^

The Rush Memory and Aging Project (MAP) is a longitudinal epidemiologic clinical-pathologic cohort study, emphasizing common chronic conditions of aging, cognitive decline, and Alzheimer’s risk. It includes older men and women recruited from retirement communities and senior housing facilities in Chicagoland and northeastern Illinois. Enrollment began in 1997, and all participants undergo annual clinical evaluation, cognitive testing, and annual blood draw. All participants agree to organ donation (e.g., brain, spinal cord, nerve and muscle) and sign an Anatomic Gift Act.

#### Minority Aging Research Study (MARS)^24^

The Minority Aging Research Study (MARS) is a longitudinal epidemiologic cohort study focusing on cognitive decline and Alzheimer’s risk in older African Americans. Recruitment began in 2004, exclusively enrolling older African American men and women who undergo annual clinical evaluation, blood draw, and brain donation is optional.

#### Rush Clinical Core^25^

The Rush Clinical Core is a longitudinal epidemiologic clinical-pathologic cohort study examining the transition from normal aging to mild cognitive impairment (MCI) to early dementia. Recruitment of the current cohort began in 2008. Participants, primarily older African Americans from the Chicago area, undergo annual clinical evaluation, blood draw, and brain donation is optional.

#### Latino CORE Study (LATC)^26^

The Latino CORE Study (LATC) is a longitudinal epidemiologic cohort study investigating cognitive decline and Alzheimer’s risk in older Latinos. Recruitment started in 2015, enrolling older Latino/Hispanic adults who undergo annual clinical evaluation, blood draw, and brain donation is optional.

### AD phenotype measures

We focused on the following two continuous measures of AD pathological traits, which have higher statistical power than discrete classifications.

#### Rate of cognitive declin

estimated person-specific rate of change in the global cognition variable over time. Global cognition is assessed annually using 21 cognitive tests, 17 of which are shared across studies and used to derive summary measures for five cognitive domains: episodic memory, semantic memory, working memory, visuospatial ability, and perceptual speed. The global cognition score is computed by averaging the standardized scores of these 17 tests. The methods for assessing cognition are detailed in previous publications^30–32^. The person-specific slope of cognitive decline was then measured by modeling the annual global cognitive scores using a linear mixed effects model, controlling for age at baseline, sex and years of education^32^.

#### Aβ and tau pathology burden

Pathological measures were collected (as described previously in detail^26,28,29^). Briefly, percent area of cortex occupied by Aβ or abnormally phosphorylated tau-positive neurofibrillary tangles, which were measured on fixed autopsy tissue by immunohistochemistry. Aβ immunostaining was performed with one of three monoclonal antibodies: 4G8 (1:9000; Covance Labs, Madison, WI), 6F/3D (1:50; Dako North America Inc., Carpinteria, CA), or 10D5 (1:600; Elan Pharmaceuticals, San Francisco, CA). Paired helical filament (PHF) tau proteins were detected using the phosphorylated tau-specific antibody AT8 (Innogenetics, San Ramon, California, USA; 1:1000). Immunostaining was conducted as previously described on 6 or 20 µm paraffin-embedded tissue sections collected from 8 brain regions (the hippocampus, entorhinal cortex, anterior cingulate cortex, midfrontal cortex, superior frontal cortex, inferior temporal cortex, angular gyrus, and calcarine cortex), followed by conversion to or new image analysis for quantification^29,34,35^. Global measures for Aβ and tau burdens were summarized by averaging the measurements obtained from the eight brain regions. In the association analysis, we used the square-root transformed burden measures for better statistical properties.

### Other neuropathological measures

Other neuropathologies were also measured using light microscopy or immunohistochemistry as previously described^94^. Presence of TAR DNA-binding protein 43 (TDP-43) cytoplasmic inclusions in neurons and glia were assessed in eight brain regions (amygdala, entorhinal cortex, hippocampus CA1, hippocampus dentate gyrus, anterior temporal pole cortex, midtemporal cortex, orbital frontal cortex, and midfrontal cortex) using a phosphorylated monoclonal TAR5P-1D3 antibody (pS409/410)^95^. TDP-43 pathology was staged into four categories: none, amygdala only, amygdala + limbic, and amygdala + limbic + neocortical. Lewy body disease was evaluated in four stages (none, nigral-predominant, limbic-type, neocortical-type and also amygdala predominant) across seven regions (substantia nigra, anterior cingulate cortex, entorhinal cortex, amygdala, midfrontal cortex, superior or middle temporal cortex, inferior parietal cortex) using α-synuclein immunostaining with monoclonal antibody LB509^96^. Cerebral amyloid angiopathy (CAA) pathology was analyzed in four neocortical regions (midfrontal, midtemporal, parietal, calcarine cortices) using one of three monoclonal anti-amyloid-β antibodies: 4G8, 6F/3D, or 10D5^97^. Amyloid-β deposition in meningeal and parenchymal vessels was scored from 0 to 4, and the scores across regions were averaged to obtain a continuous composite score for CAA^97^. Additionally, large vessel cerebral atherosclerosis and small vessel arteriolosclerosis were graded based on visual and histological examination of arterial vessels after paraformaldehyde fixation and H&E staining^98,99^. Chronic cerebral gross infarctions were examined on 1cm slabs of brain and confirmed on H&E. Microinfarcts were assessed in a minimum of nine brain regions stained with hematoxylin and eosin (H&E)^100^. Hippocampal sclerosis was identified by neuronal loss and gliosis in CA1 or subiculum regions stained with H&E^101^.

### Generation of single-nucleus RNA & ATAC-seq dataset (CUIMC2 dataset)

#### Nuclei Isolation

Nuclei were isolated with EZ PREP buffer (Sigma, Cat #NUC-101). Tissue samples cut into pieces < 0.5 cm or cell pellets were homogenized using a glass dounce tissue grinder (Sigma, Cat #D8938) (25 times with pastel A, and 25 times with pastel B) in 2 ml of ice-cold EZ PREP and incubated on ice for 5 minutes, with additional 2 ml ice-cold EZ PREP. Nuclei were centrifuged at 500 x g for 5 minutes at 4°C, washed with 4 ml ice-cold EZ PREP and incubated on ice for 5 minutes. After centrifugation, the nuclei were washed in 4 ml Nuclei Suspension Buffer (NSB; consisting of 1x PBS, 0.01% BSA and 0.1% RNAse inhibitor (Clontech, Cat #2313A)). Isolated nuclei were resuspended in 100-500uL NSB depending on pellet size, filtered through a 35 μm cell strainer (Corning, Cat # 352235). Nuclei were counted using the Nexcelom Cellometer Vision 10x objective and an AO/PI stain. 20µl of the AO/PI was pipet mixed with 20ul of the nuclei suspension and 20µl was loaded onto a Cellometer cell counting chamber of standard thickness (Nexcelom catalog number: CHT4-SD100-002) and counted with the dilution factor set to 2. Determine volume needed for 250,000 nuclei from each donor to create a 3-plex donor pool. Pooled nuclei were centrifuged at 500 x g for 5 minutes at 4°C. Nuclei pellet was resuspended in 50-70uL NSB depending on pellet size, filtered through a 35 μm cell strainer. Nuclei were counted with AO/PI again. A final target of 21,500 nuclei was used for 10x Single Cell Multiome ATAC + Gene Expression experiments.

#### Single-nucleus RNA & ATAC-seq profiling

After nuclei isolation the 10x multiome protocol (Chromium Next GEM Single Cell Multiome ATAC + Gene Expression Reagent Bundle, PN-1000283) was followed according to manufacturer’s instructions (10X Genomics, USA) and can be accessed at https://cdn.10xgenomics.com/image/upload/v1666737555/support-documents/CG000338_ChromiumNextGEM_Multiome_ATAC_GEX_User_Guide_RevF.pdf.

Briefly, following transposition GEMs were generated by combining barcoded Gel Beads, transposed nuclei, a Master Mix that includes reverse transcription (RT) reagents, and Partitioning Oil on a Chromium Next GEM Chip J (10× Genomics; PN-2000264). Incubation of the GEMs in a thermal cycler for 45 minutes at 37°C and for 30 minutes at 25°C generates full-length cDNA from poly-adenylated mRNA for gene expression (GEX) library and a Spacer sequence that enables barcode attachment to transposed DNA fragments for Assay for Transposase-Accessible Chromatin (ATAC) library. This was followed by a quenching step that stopped the reaction. Next, GEMs were broken, and pooled fractions were recovered. Silane magnetic beads were used to purify the first-strand cDNA from the post GEM-RT reaction mixture. Barcoded transposed DNA and barcoded full-length cDNA from poly-adenylated mRNA were pre-amplified by PCR and the products were used as input for both ATAC library construction and cDNA amplification for gene-expression library construction. Libraries were pooled and sequenced together on a NovaSeq 6000 with S4 flow cell (Illumina, San Diego, CA). The scATAC-seq libraries were sequences as follows: Read 1N, 50 cycles; i7 Index, 8 cycles; i5 Index, 24 Cycles; Read 2N, 49 cycles. The snRNA-seq libraries were sequenced as follows: Read 1, 28 cycles; i7 Index, 10 cycles; i5 Index, 10 Cycles; Read 2, 90 cycles.

### Single-nucleus RNA & ATAC-seq data processing

#### Alignment and technical artifact adjustment

Through the application of CellRangerArc (v2.0.2), the raw sequence reads were aligned to human genome hg38. To adjust for technical artifacts from background RNA and random barcode swapping, the *remove-background* function from CellBender (v0.3.1) was conducted over the RNA matrices from CellRanger. In short, CellBender models from each library’s gene expression matrix levels of empty droplets (low UMI count/droplet with high droplet counts) and removes empty/contaminated droplets. The parameters set for CellBender *remove-background* include: epochs=150, fpr=0.01, expected-cells=5000, and total-droplets-included set to the number of cells in the library from the CellRanger gene expression matrix.

#### Demultiplexing

Demultiplexing was performed using the demuxlet software (v1.11). Given each library contains 3 or 4 individuals, and each individual has their own WGS VCF file from ROSMAP, we were able to identify which cell belonged to which study sample. Succinctly, BAM files with filtered VCF were taken from the output of CellRanger, popscle generates a pileup file required for demuxlet, and demuxlet assigned scores to identify corresponding cells to study samples.

#### Normalization and clustering

The normalization pipeline for the RNA gene expression matrices began with the *NormalizeData* method from Seurat (v4.4.0), followed by *FindVariableFeatures* (selection.method = ‘vst’, nfeatures=2000), *ScaleData*, dimensional reduction with *RunPCA* (npcs=30), creation of k-NN neighbor with *FindNeighbors* (dims=1:30), and cluster detection with *FindClusters* (resolution = 1.5, algorithm = 1).

#### Cell Type Classifications

Cell types were annotated with weighted ElasticNet linear regression trained off prior atlas of human ROSMAP dorsolateral prefrontal cortex tissue (n=24)^2^ where 7 major cell types are present (excitatory neurons, inhibitory neurons, vascular cells, oligodendrocytes, astrocytes, microglia, and oligodendrocyte precursor cells). Weights were applied as the inverse of the number of nuclei per cell type to preserve classifications of smaller cell types like the vascular cells. Cross validation was performed to optimize regulation parameters using *glmnet* (*cv.glmnet*) (v4.1) with an alpha=0.25, nfolds = 10, family = ‘multinomial’.

### Single-nucleus RNA & ATAC-seq data quality control

#### Low quality cell assignment and removal

Given biological differences in the numbers of genes expressed (nGene) and the number of UMIs (nUMI) between different cell types, we optimized seven different thresholds for each cell type where cells with nGene or nUMI under the thresholds would be labeled as low quality. To do so, we first picked 6 out of the 61 libraries (RA006, RA007, RA010, RA018, RA035, RA039) controlling for sex, age at death, and AD prevalence. We then manually identified clusters within each library that are low quality, judging by the overlap of cell type classifications, a high mitochondrial gene expression, and a low nGene/nUMI relative other clusters. Afterwards, we derived a harmonic mean using precision/recall to identify nUMI/nGene thresholds per cell type: excitatory neurons=2331/1068, inhibitory neurons=857/397, vascular cells=146/493, oligodendrocytes=776/490, astrocytes=528/442, microglia=291/150, and oligodendrocyte precursor cells=667/554.

#### Doublet removal

To account for doublets across samples, demuxlet doublet identification based off of WGS-VCF files of each sample was applied. Additionally, to address doublets across different cell types (e.g. Excitatory Neuron-Microglia doublet), we utilized Gilad Green’s customized DoubletFinder’s *DoubletFinder_v3* method (https://github.com/GreenGilad/DoubletFinder) where pN=0.5, pK=75/(1.5*(#nuclei/library), nExp=0, sct=T, labels=cell type information for each library. Thus any nuclei classified as either a demuxlet doublet or DoubletFinder doublet were subsequently removed.

#### Mitochondrial gene expression

Following guidelines from the 10X genomic multiome page, we filtered out cells possessing a mitochondrial RNA percent expression outside of 2 standard deviations within each library (https://www.10xgenomics.com/analysis-guides/common-considerations-for-quality-control-filters-for-single-cell-rna-seq-data).

### Cell state annotations transfer across datasets

To replicate cell subset associations with AD pathologies using independent snRNAseq datasets, we first annotated the cellular states of individual cells from each cell type separately, based on the cell-state definitions from Green et al^1^. For each cell type, we mapped cells from each strata of the replication dataset as query onto our reference CUIMC1 Discovery dataset, using the Seurat query mapping pipeline^39^. Briefly, the reference and query data were first normalized, and the top variable genes were identified using the *SCTransform* function while regressing out percent of mitochondrial reads. A set of mapping anchors (pairs of cells that encode the relationships between the reference and query) was identified and scored using the *FindTransferAnchors* function with default parameters. These anchors were then used to transfer the cell-state annotations from the reference to the query data using the *TransferData* function. In this pipeline, we varied three key parameters in this pipeline, while keeping other parameters as default: the number of top variable genes (*variable.features.n*, ranged 1000 – 4000), the number of principal components for finding anchors (*npcs, ranged 10* –*100)*, and the number of neighbors to consider when weighing anchors (*k.weight*, ranged 10 – 50). The parameters set for each cell type in each dataset are detailed in **Supplementary Table 1**.

### Cell-state frequency calculation

We grouped the cell subpopulations into seven broad cell types as previously defined^1^: astrocytes, microglia, inhibitory neurons, excitatory neurons, oligodendrocytes, oligodendrocyte precursor cells (OPCs) and vascular niche, which includes endothelial cells, arteriole cells, venule cells, pericytes, smooth muscle cells (SMC) and fibroblasts. We excluded rare subpopulations such as macrophages, monocytes, erythrocytes, CD8^+^ T cells, NK cells and neutrophils due to the low abundances. Here, we define the frequency (or relative proportion) of a cell subpopulation as the number of cells of the *cell-state* of interest divided by the total number of cells of the same *cell-type*. For each donor, the frequency of subpopulation *s* is calculated as:

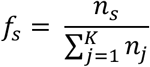

where *ns* is the number of cells for cell-state *s* and *K* is the total number of cell-states within the cell-type to which cell-state *s* belongs. Hence, the cell-state frequency here reflects the empirical probability of a cell of a given cell-type acquiring the transcriptional state of interest.

### Statistical analysis associating AD traits and cell-state frequencies

For each dataset, statistical associations between AD traits and cell subset frequencies were tested using linear regression models as previously described^1^: 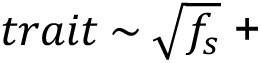 covariates, where *fs* is the frequency of cell subpopulation *s* (defined in the previous section). The frequencies were square-root transformed to better capture the subtle changes of the less abundance cell subsets. To adjust for potential confounding variables, we included the following variables as covariates: 1) participants characteristics (sex, age, post-mortem interval, education), and 2) snRNA-seq library quality metrics (cell count and median of gene count for each participant). Results from the linear regression models were corrected for multiple hypothesis testing by calculating the false-discovery rate within each AD trait using the Benjamini-Hochberg correction method.

To integrate the statistical associations across datasets, we performed fixed-effect meta-analyses using the *metagen* function from the meta (v8.0-1) package^102^ in R (v4.4.2). The meta-analyses were performed using an inverse-variance based approach by summing the effect size (*β_i_*) weighted by the inverse of standard error (*SE_i_*) of each study *i*. Mathematically, the summarized z-statistics for each association was calculated as:

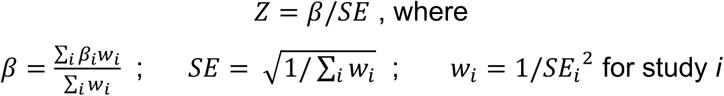

Meta-analysis results were then corrected for multiple-hypothesis testing using the Benjamini-Hochberg method within each AD trait. To assess the heterogeneity of statistical associations across studies, we calculated the *I*^2^ statistics^36^

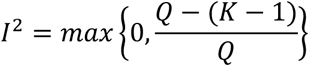

where K is the number of studies and Q is the Cochran’s Q^103^:

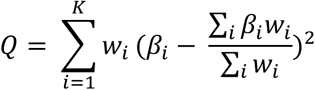

### Identification of cell state signatures

To identify signature genes for a specific cell state, we performed pairwise differential expression (DE) analysis, comparing the gene expression of the cell state of interest against each of the other cell states within the same cell type. The input for the DE analysis was a pseudobulk expression matrix, generated by aggregating UMI counts for each gene across cells in each cell state for each individual. Lowly expressed genes were filtered using the *filterByExpr* function from the edgeR package (v3.40.2) with default parameters. To account for the mean-variance relationship across genes, we applied the voom function from the limma package (v3.54.2), which generated a set of precision weights. These weights were subsequently used to fit a linear model for each gene with the *lmFit* function. The statistics and fold changes of the linear model fit were computed using the *eBayes* function (with trend = FALSE and robust = FALSE).

To ensure specificity of the signature genes, we identified genes that were significantly differentially expressed (adjusted *P* < 0.05) in the same direction (either upregulated or downregulated) across *all* significant pairwise comparisons involving the cell state of interest. Genes exhibiting inconsistent regulation (upregulated in some comparisons and downregulated in others) were excluded. To further refine the selection of uniquely differentiating genes, we retained only those genes with at least one pairwise comparison showing a significant log-fold change (LFC) magnitude greater than 1, while ensuring that no comparisons had an LFC magnitude below 0.25. The final list of signature genes used for downstream analyses is provided in **Supplementary Table S2**.

### Hypergeometric enrichment of astrocyte subtype signatures against literature-defined astrocyte states

To assess transcriptional correspondence between the astrocyte subtypes defined by Green et al. and previously reported astrocyte states, we performed pairwise gene-set enrichment analyses using one-sided hypergeometric tests. To maximize subtype specificity, marker genes for Green et al. astrocyte subtypes were defined from pairwise differential expression analyses as genes significantly and uniquely upregulated in each subtype relative to all other astrocyte subtypes (see section *Identification of cell state signatures*). For subtypes with fewer than 10 pairwise-unique upregulated genes using this criterion (Ast1, Ast2 and Ast6), published Green et al. one-versus-all markers were used as a fallback, restricted to genes with positive average log2 fold change. To limit the influence of highly gene-rich marker sets, Ast.7 markers were capped at the top 200 genes ranked by average log2 fold change.

Published astrocyte state signatures were compiled from prior single-cell, single-nucleus and bulk transcriptomic studies, including Zamanian et al.^40^, Liddelow et al.^16^, Habib et al.^8^, Prakash et al. ^104^ and Serrano-Pozo et al.^80^ Gene sets from mouse studies were mapped to human orthologues using the *nichenetr* package, and genes without a mapped human orthologue were excluded. Literature-defined signatures containing fewer than 10 genes after orthologue mapping and filtering were excluded from downstream analyses.

The background gene universe was defined as the set of genes that passed expression filtering with filterByExpr in the pseudobulk differential expression analysis, thereby restricting enrichment tests to genes that were detectable and tested in the Green et al. dataset. For each Green et al. astrocyte subtype marker set and each literature-defined astrocyte state signature, both gene sets were intersected with this background universe. We then computed the observed overlap and tested for enrichment using a one-sided hypergeometric test, corresponding to probability P(X *≥ k*), where *k* is the observed number of overlapping genes. *P* values from all pairwise comparisons were adjusted using the Benjamini-Hochberg procedure.

### Structural equation modelling of the AD cellular cascade

We used structural equation modelling (SEM) to test whether the AD cellular cascade proposed by Green et al^1^ applies to new datasets. This framework builds on the widely accepted sequence of AD progression in which Aβ accumulation precedes tau pathology, which in turn is linked to cognitive decline, and positions disease-associated glial subpopulations along this cascade^1^.

We focused on the microglial, astrocyte and oligodendrocyte subpopulations implicated in the Green et al. AD cascade: Mic.12, Mic.13, Ast.10 and Oli.7. Cell-state abundances were represented as square-root-transformed subpopulation frequencies. Aβ and tau pathology were measured in the mid-frontal cortex and square-root transformed. Cognitive decline was represented by the demographic-adjusted longitudinal cognitive slope. Analyses were restricted to participants from the CUIMC 1, CUIMC 2 and MIT cohorts with complete data for all model variables.

Before SEM fitting, we residualized variables within each cohort to remove demographic and technical confounding. Aβ and tau pathology were residualized for age at death and sex. Mic.12, Mic.13, Ast.10 and Oli.7 frequencies were residualized for age at death, sex, post-mortem interval, median genes detected per nucleus and total cell count. Cognitive decline was not further residualized because the cognitive slope was already adjusted for demographic covariates. Residualized values were then pooled across cohorts.

The SEM was specified to reflect the Green et al. cascade (**Extended Data Figure 1I**)^1^, with paths from Mic.12 to Aβ, from Aβ and Mic.12 to Mic.13, from Aβ and Mic.13 to tau, from tau and Mic.13 to Ast.10, from Ast.10 to Oli.7 and from both tau and Ast.10 to cognitive decline. Models were fit using the *lavaan* package (v0.6) in R with maximum likelihood estimation. Standard errors and confidence intervals were estimated using 10,000 bootstrap iterations. Standardized path coefficients, bootstrap confidence intervals and nominal *P* values were extracted for each path. Model fit was evaluated using the comparative fit index, root mean square error of approximation and standardized root mean square residual. SEMs were fit separately in the CUIMC 1 Discovery cohort, in the pooled replication cohorts (CUIMC 2 and MIT) and in all cohorts combined. To test alternative models, we applied the same approach except swapping the Ast10 node with Mic13 or with tau (**Extended Data Figure 1J**).

### Inference of cell-cell signaling drivers of glial states

To identify the cell-cell signaling that drives the transcriptional programs underlying a specific glial state (Ast10), we applied NicheNet^41^ using its R implementation nichenetr (v2.0.0). NicheNet prioritizes ligand-receptor pairs according to their downstream signaling activities. Briefly, NicheNet models ligand activities by utilizing three types of prior knowledge from several complementary resources, including ligand-receptor, signaling and gene-regulatory interactions. NicheNet integrates such prior knowledge into an expansive intracellular signaling network where each ligand-to-target link is given a score, termed *regulatory potential*. Such prior information is then used to infer potentially active ligand-receptor interactions by searching for highly differentially expressed ligand-receptor pairs with high regulatory potential scores for the given set of target genes in receiver cells.

NicheNet requires three primary inputs: (1) the target gene set, (2) the background gene set, and (3) expression data for sender and receiver cell populations. For our analysis, we defined the Ast10 transcriptional signature (see *Identification of cell state signatures*) as the target gene set and all genes available in the NicheNet ligand-target prior model as the background. Ast10 was designated as the receiver, while the 10 astrocyte subsets (including Ast10 itself) and 16 microglial subsets were set as sender populations.

Ligand activities were quantified using the NicheNet function *predict_ligand_activities* with default parameters, with activity measured as the area under the precision-recall curve (AUPR). This metric reflects the concordance between a ligand’s predicted targets and the observed transcriptional signature relative to background genes. Ligands were then prioritized using weighted criteria, including scaled ligand activity (*activity_scaled* = 2), differential upregulation of sender ligands (*de_ligand* = 1) and receiver receptors (*de_receptor* = 1), and the average expression of sender ligands (*exprs_ligand* = 1) and receiver receptors (*exprs_receptor* = 1).

Based on this prioritization, we selected the top 100 sender-ligand-receptor combinations, resulting in 49 unique ligand-receptor interactions comprising 41 ligands and 27 receptors for further analysis.

### Partial least squares regression (PLSR) modeling

To identify cell-cell signaling associated with Ast10 emergence, we applied partial least squares regression (PLSR)^105^ to model the relationship between astrocyte receptor or sender-specific ligand expression and donor-level Ast10 frequency. We use donor-level Ast10 frequency as a quantitative readout of the extent to which an individual’s astrocyte population is shifted toward the Ast10 state. Within this framework, sender-specific ligands and astrocyte receptors whose expression covaries with high or low Ast10 frequency were prioritized as candidate signaling cues associated with the emergence or depletion of the Ast10 state. Unlike principal component analysis (PCA), which identifies orthogonal components explaining the maximum variance in the data without regard to an outcome, PLSR is a supervised regression technique, identifying latent components that maximize covariance between the predictors and the outcome. This property makes PLSR well suited for linking high dimensional, coordinated expression patterns to Ast10 frequency across donors. However, we emphasize that PLSR identifies associative relationships and serves to prioritize candidate regulators; causal inference requires convergent evidence from complementary analyses, including ligand-receptor pathway inference (NicheNet), spatial transcriptomics (MERFISH), and targeted perturbation experiments.

#### Model construction

**1. Receptor Model:** The input matrix for the receptor model consisted of the pseudobulk expression (log-transformed mean expression) of NicheNet-prioritized receptors in all astrocytes across donors. The output vector consisted of the log-transformed percentage of Ast10 in each donor with an offset of 1.
**2. Cell-state Specific Ligand Model:** The input matrix consisted of pseudobulk expression of NicheNet-prioritized ligands in specific sender cell subsets (microglial or astrocytic states). To reduce noise and address sparsity in glial state representations across donors, sender-ligand pairs were filtered to retain only those where ligand expression was significantly correlated with Ast10 frequency across donors (|Pearson’s r| ≥ threshold, *P* < 0.05). The optimal correlation threshold was determined by parameter sweep (|r| = 0.10, 0.15, 0.20, 0.25; see below). For ligands with at least one significant sender, a summary score was calculated by averaging the scaled expression (zero-centered) across those senders. In cases where a ligand (e.g., NRG3) exhibited opposing correlations with Ast10 frequency across different sender subsets (i.e., positive in some and negative in others) separate summary scores were calculated for each direction. Notably, during each iteration of model training (see below), this correlation-based filtering and sender-cell-ligand aggregation rules were derived using only the donors in the training set to ensure no data leakage. The output vector for this model was identical to that used in the receptor model.

#### Bagged PLSR Training

All modeling steps, including training, cross-validation, and testing with external datasets, were conducted in Python (v3.9.2) using the scikit-learn library (v0.24.1)^106^. PLSR was implemented using the *PLSRegression* algorithm with repeated random cross-validation splits and bootstrap aggregation (bagging) to improve prediction stability and robustness of feature importance estimates (**Extended Figure S8A**). Across 2,000 iterations, donors were randomly partitioned into training (80%) and held-out validation (20%) sets. Within each iteration, 50 bootstrap samples were drawn with replacement from the training set, and individual PLSR models were fit on each bootstrap sample.

To prevent data leakage, predictor features and response variables were standardized to zero mean and unit variance based on statistics derived from the training data only, using the StandardScaler() function. For each bootstrap model, the optimal number of PLSR components was determined through 5-fold cross-validation within the bootstrap sample, minimizing overfitting by balancing predicted variance in cross-validation (Q^2^) against variance explained (R^2^). R^2^ and Q^2^ were calculated using the *r2_score* function to assess the concordance between the empirical Ast10 frequency estimated from the data and model prediction. For a given iteration, predictions for held-out samples were obtained by averaging predictions across the optimal bootstrap models.

Specific to the ligand model, different correlation thresholds (|r| = 0.10, 0.15, 0.20, 0.25) were tested using the aforementioned procedure and the optimal threshold was selected by maximizing mean predicted variance (Q^2^) on the held-out validation sets across iterations. The receptor model, which uses all NicheNet-prioritized receptors in astrocytes without correlation-based filtering, proceeded directly to variable importance quantification.

#### Variable Importance in Projection (VIP)

To assess the relative importance of individual ligands or receptors in the PLSR models, we calculated the Variable Importance in Projection (VIP) scores at the optimal number of PLSR components for each model. VIP scores summarize the contribution of each input variable (ligand or receptor) to the model, weighted by the variance explained by each PLS component. Mathematically, the VIP score for a given variable *x* is calculated as:

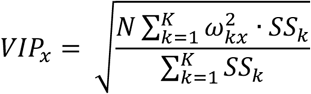

where *N* is the total number of input variables (28 ligands or 26 receptors), *wkx* is the weight of the *x^th^* variable for the *k^th^* PLS components, and *SSk* is the sum of squares explained by the *k^th^* PLS component. To help interpret the directionality of the associations, the VIP scores for each ligand or receptor were multiplied by the sign of the Pearson’s correlation coefficient between the predictor (ligand or receptor expression) and Ast10 frequency. Within each bootstrap model, |VIP| ≥ 1 indicates significant contribution. To obtain robust importance estimates, VIP scores were averaged across the 50 bootstrap models within each iteration, then summarized across all 2,000 iterations. Stability for each feature was defined as the proportion of iterations in which |VIP| ≥ 1.

#### Final Model Selection

To determine the final optimal model, we ranked each feature by their VIP stability, and performed a forward selection procedure: starting with the most stable feature, features were sequentially added to the model in order of decreasing stability. Model performance (Q^2^) was evaluated at each step using leave-one-out cross-validation (LOOCV) on *CUIMC1*, and the final model was trained using only features up to the point where additional features no longer improved predictive performance. This final model trained on *CUIMC1* is then used for *independent testing* with *CUIMC2* and *MIT* datasets which were not used in any way during model construction.

### Gene-gene co-expression assessment

To examine gene-gene co-expression relationships that may be obscured by transcript capture sparsity in single-cell and single-nucleus RNA-seq data, we applied MAGIC^107^ (Markov affinity-based graph imputation of cells), which uses data diffusion across transcriptionally similar cells to smooth gene expression measurements.

We implemented MAGIC in R using the Rmagic package (v2.0.3). For the Discovery snRNA-seq dataset, we randomly sampled 2,500 cells per astrocytic state (25,000 astrocytes total) to ensure balanced representation. Raw counts were normalized using the library.size.normalize function from phateR (v1.0.7), followed by square-root transformation. MAGIC was applied to genes expressed in at least 10 cells, using k-nearest neighbors (*knn*) = 10 and Markov matrix power (*t*) = 5.

For the iAst scRNA-seq dataset, MAGIC was applied separately to wild-type cells from each of the two donor lines to examine Ast10 gene-gene co-expression structure. Following gene filtering, library-size normalization and square-root transformation, MAGIC was performed using *knn* = 5 and *t*=3. Pairwise Pearson correlations were calculated across Ast10 signature genes and averaged between donor lines to identify the two co-expression modules used in subsequent gene-set enrichment analyses.

MAGIC-imputed expression was used for exploratory gene-gene co-expression analyses and the DREMI analysis of ligand-associated signaling pathways in the Discovery dataset (see DREMI Methods). Importantly, MAGIC-imputed expression was not used for differential gene expression testing or the calculation of gene-ranking statistics for gene-set enrichment analyses. For PLXNB1-knockout experiments in both iAst and mouse astrocytes, differential expression was assessed using Nebula on raw counts (See *Differential expression analysis to assess PLXNB1-knockout effect in scRNAseq data*).

### Signature summary score

To evaluate the overall expression of a specific gene signature or gene set in individual cells, we utilized the *AddModuleScore* function in Seurat to derive a summary measure for that signature. The function computes the summary score for each cell as the average expression across the signature genes subtracted by the aggregated expression of a set of randomly selected control features^108^.

For DREMI analysis, signature scores were calculated using the MAGIC-derived expression of differentially *upregulated* genes in the Ast10 signature, as identified in the differential expression analysis. For the spatial analysis of the Visium data, we assessed correlations at the spot level in the between two summary measures, 1) the prioritized Ast10-predicting ligand signature and 2) astrocyte-state signatures. Here, we computed the ligand signature score using normalized expression values (See *Spatial Correlation Analysis* section below) of the set of Ast.10 ligands with significant and positive VIP scores (VIP ≥ 1 in at least 50% of training iterations). Similarly, signature scores for other astrocytic states were computed using the normalized expression of their respective differentially upregulated marker genes, except for Ast.1, which lacked uniquely upregulated markers.

To avoid confounding the correlations, any genes in question were excluded from the cell-state signature score calculations in all association analyses described above.

### Identify ligand-associated pathway genes

For the prioritized ligand set in the final PLSR model (*JAM2, NCAN, GNAS, SEMA4D, RGMA, DSCAML1, PTPRK, EFEMP1, LGALS3, NRXN1, SEMA4B, FGF1, and A2M*), we focused on reconstructing its pathway leading to the Ast10 differentially expressed genes that are among the top 300 targets of the ligand according to the NicheNet database. To determine the signaling pathways through which a ligand is predicted to regulate the Ast10 signature genes of interest, we used the *get_ligand_signaling_path_with_receptor* function from NicheNet to identify the intermediate genes.

This function first identifies the top 5 transcription factors that best regulate the target genes and are closest to the upstream ligand, based on prior knowledge of the ligand-signaling and gene regulatory networks which are represented by a set of weights. These weights were then used to determine the shortest paths between these top transcription factors and the ligands. Genes along these shortest paths are considered as significant nodes within the pathway. To classify their roles within the pathway, these pathway nodes were annotated as one of the following categories: ligands, receptors, signaling mediators or transcriptional regulators, based on the OmniPath (R package OmnipathR, v3.9.3) *intercell* resources^109^ (for ligand, receptor annotations) as well as the *TFcensus*^110^ and *DoRothEA* resources^111^ (for transcriptional regulator annotations).

### DREMI to quantify pathway information flow

To quantify the relationships between a given ligand-associated pathway gene and Ast10 signature, we used a metric called DREMI (conditional-Density Resampled Estimate of Mutual Information)^52^. DREMI leverages the cell-to-cell variation in single-cell data to quantify the amount of information transmitted from gene X (i.e. pathway gene of interest) to gene Y (i.e. Ast10 signature score) in signaling networks. Notably, DREMI is a directional measure, as it quantifies the information flow using conditional density (i.e. probability of Y given X) rather than the joint density of X and Y. In estimating the conditional densities we adopted the k-nearest neighbor based-DREMI that was previously applied in scRNA-seq data following MAGIC implementation^107^.

To set the baseline for the DREMI score, we randomly permutated the gene expression vector X, then calculated a DREMI score between the permuted X and the Ast10 signature score. We repeated this process for each pathway gene in question, then average their DREMI scores to set the baseline. To evaluate the significance of each pathway gene’s DREMI score, we first derived DREMI scores for all genes available in each of these resources from OmniPath : *intercell* resources^109^ (for the ligand, receptor gene categories), the *TFcensus*^110^ and *DoRothEA* resources^111^ (for the transcriptional regulator gene category), as well as 10779 randomly selected genes that do not belong to any of the above categories (for the signaling mediator gene category). We then calculated an empirical *P*-value (*Pemp*) *for* gene *x*, based on all genes within the same signaling node category:

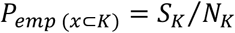

where *Sk* is the number of genes with a higher DREMI score than gene *x* within the signaling node category *K* (i.e., ligand, receptor, signaling mediator or transcriptional regulator) and *Nk* is the number of genes in the signaling category *K*.

### Visium data generation

The spatial transcriptomic data with Visium combined with immunofluorescence was generated as previously described^1,55^. Fresh-frozen dorsolateral prefrontal cortex (DLPFC) samples from ROSMAP participants were processed using the Visium Spatial Transcriptomics (ST) platform, coupled with immunofluorescence. Tissue sections encompassing all cortical layers and some white matter were selected based on RNA quality (RNA Integrity Number > 6). RNA was purified using the RNeasy Micro Kit (Qiagen, #74004) and RNA integrity was assessed using TapeStation 4150 and Bioanalyzer 2100 (Agilent Genomics).

Tissue blocks were prepared into ∼6 × 6 mm^2^, 10 μm thick sections embedded in Optimal Cutting Temperature (OCT; Sakura, 4583) and were mounted on Visium Tissue Optimization Slides and Gene Expression Slides (10x Genomics) in duplicates. Permeabilization time was optimized based on the Visium Spatial Tissue Optimization protocol (10x Genomics, CG000238 Rev D) with modifications. After fixation in cold 100% ethanol (–20°C for 30 minutes), sections were stained with Thioflavin S (ThioS; Sigma, T1892) to label neuritic plaques and an anti-GFAP Cy3-conjugated antibody (Millipore Sigma, MAB3402C3, dilution 1/50) to visualize astrocytes. Autofluorescence from lipofuscin was quenched using TrueBlack (Biotinum, 23007). Sections were then briefly immersed in 3X SSC buffer (Millipore Sigma, S6639L) and mounted with a solution containing 85% glycerol, 2 U/μL RNase inhibitor, and DAPI (Thermo Fisher, #62248).

Sections were imaged at 10X magnification using a Nikon Eclipse Ni-E immunofluorescence microscope at 10X magnification (Plan Apo λ, NA = 0.45). Following imaging, coverslips were removed, and sections were permeabilized to optimize mRNA release. A 3-minute permeabilization time was identified as optimal for the Visium workflow, balancing effective transcript release and minimal diffusion. Subsequent cDNA synthesis, library preparation, and sequencing were performed following manufacturer protocols (10x Genomics, Visium Spatial Gene Expression Reagent Kits, CG000239 Rev D), with sequencing conducted on a NovaSeq 6000 (Illumina). ST spots (diameter = 55µm) were aligned with brightfield images.

### Visium data preprocessing and cortical layer annotations

Visium sequencing reads were aligned to the human reference genome (GRCh38) and were assigned to spots using Space Ranger (v2.0.0, 10x Genomics). The resulting h5 files for each sample were imported into R (v4.2.2) using the SpatialExperiment (v1.16.0). For quality control, mitochondrial and ribosomal genes, as well as genes expressed in fewer than 10 spots, were excluded. Additionally, spots located outside the tissue or those with ≤ 1,000 UMIs or with ≤ 500 expressed genes were removed.

Cortical layers and white matter were identified as previously described^55^, by first selecting the top 10,000 most variable genes using the *modelGeneVar* function from the scran package (v1.26.2)^112^. To focus on layer-specific features, we extracted 3,003 marker genes by taking the top 429 marker genes for each of the six cortical layers and the white matter, as reported in a previous study^113^. We then intersected these marker genes with our highly variable genes, resulting in a final set of 1,357 highly variable marker genes. Principal component analysis (PCA) was then performed on this gene set, and the first 50 principal components (PCs) were used to mitigate batch effects using Harmony (v1.2.0), treating each tissue section as a separate batch and applying parameters *theta* = 3, *sigma* = 0.1, and *lambda* = 1. To identify spatial clusters, we applied BayesSpace (v1.8.2)^114^ to the first 35 Harmony-corrected PCs, using a spatial smoothing parameter of *gamma* = 2. This approach partitioned the spots into seven clusters, corresponding to six cortical layers and white matter. White matter spots were excluded from all subsequent analyses.

### Visium spatial correlation analysis of nearby Ast10-predicting ligands

For each tissue slide, we assessed Pearson’s correlations between two summary measures at the spot level in the Visium data: 1) the averaged ligand signature predicting the Ast.10 state within the neighborhood, and 2) a specific astrocyte-state signature. Both signature scores were computed using normalized expression values obtained via Seurat’s *SCTransform* function with default parameters, adjusting for the number of nuclei per spot and the percentage of mitochondrial reads.

To compute the ligand signature, we derived a summary score (See *Signature summary score* section above) focusing on the prioritized Ast.10 ligands with significant and positive VIP scores (VIP ≥ 1): *JAM2, NCAN, SEMA4D, GNAS, LRRC4B, RGMA* (**Figure 3C**). For each target spot, we averaged the ligand signature scores across neighboring spots within radii of 100-500 μm (1-5 inter-spot distances), excluding the target spot itself to avoid confounding the correlation assessment. To establish a control for baseline comparison, we also averaged ligand signature scores across the same number of spots randomly selected from within the tissue for each target spot. We selected the range of neighorbood radii of 100-500 μm to encompass immediate cell-cell contact (< ∼150 μm, based on an average human astrocyte territory size diameter of 142.6 μm^57^) and broader microenvironments potentially influenced by diffusion-mediated signaling of secreted factors (∼100-300 μm^58^).

The astrocyte-state signature score for the target spot was computed similarly, using each astrocytic state’s respective marker genes (see *Identification of cell-state signatures* section). Since we focused on linking the ligands to enrichment of specific astrocyte states, we limited the analysis to states with uniquely *upregulated* markers for most astrocyte states.

Finally, all Pearson’s correlation results were corrected for multiple hypothesis testing using the Benjamini-Hochberg method to calculate false discovery rates for each astrocytic state and neighborhood size **(Extended Data Figure S7A-C)**.

### Estimation of the effects of nearby ligands on Astrocyte 10 signature genes using MISTy

To assess the importance of individual ligands to the expression of Astrocyte 10 (Ast10) signature genes in spatial contexts, we utilized the explainable machine learning framework MISTy^59^, implemented via the R package mistyR (v1.10.0). This approach allowed us to decompose the contributions of intrinsic and spatial factors to Ast10 signature gene expression at the spot level using a multi-view predictive model. The model incorporated two distinct spatial contexts as predictors:

1. **Intrinsic view:** Captures the relationships among Ast10 signature genes within the same spot. The intrinsic view included Ast10 signature genes identified from the differential expression analysis (see *Identification of receiver glial state signature* section).
2. **Juxta view:** Reflects the cumulative expression of individual ligands from immediate neighboring spots, defined as spots within a three-spot radius. In the juxta view, all potential ligands, including those prioritized for their predictive relevance to Ast10 signature from the PLSR analysis, were included as candidate predictors.

The multi-view model was fitted independently for each of the 32 slides, generating a *standardized importance score* for each ligand. This score reflects the variance reduction in the ligand’s prediction of the expression of a given Ast10 signature gene. For each ligand-signature gene pair, importance scores were averaged across all 32 slides to obtain an aggregated score (**Extended Data Figure S7D**), representing the spatial association between a ligand and an Ast10 signature gene. To evaluate each ligand’s overall contribution to the collective Ast10 signature, we first z-score transformed the aggregated importance scores within each Ast10 signature gene and then summed these z-scores across all Ast10 signature genes to obtain the total importance score. To assess the significance of the PLSR-prioritized ligand set (|VIP|≥ 1) in predicting the overall Ast10 signature, we randomly sampled ligand sets of the same size 100,000 times, computing the mean total importance score for each set. The average total importance of the PLSR-prioritized ligand set was then compared against the distribution of randomly sampled sets to derive an empirical *P*-value (**Extended Data Figure S7E**).

### MERFISH gene panel design

We curated an initial set of 500 target genes designed to capture four major categories: (1) differentially up-/down-regulated signature genes of Ast10 based on snRNAseq^1^, (2) signature marking other relevant astrocyte and microglial subsets, including homeostatic glia (astrocytic states Ast1 and Ast2; microglial states Mic2 and Mic3), reactive astrocytes (Ast4 and Ast5), and additional astrocyte or microglial states within the same cellular community as Ast10 (Ast3, Ast9, Mic11, Mic12, and Mic13)^1^, (3) downstream signaling pathways of the prioritized Ast10-associated ligand-receptor pairs, curated from NicheNet^41^ and Omnipath^109^ databases, and (4) general cell type and layer markers. We evaluated this gene panel using the MERSCOPE Gene Panel Design Portal available at Vizgen (portal.vizgen.com) to ensure that each gene is sufficiently long to allow enough encoding probes to bind, and that the entire gene panel meets the abundance threshold to avoid optical crowding for MERSCOPE imaging. Genes with fewer than 30 target regions per transcript were removed from the list. To avoid including highly expressed genes in the panel, higher than an abundance threshold of 1000 FPKM (Fragments Per Kilobase of transcript per Million mapped reads) were put in the sequential gene panel. The final 477 targetable genes possessed a total abundance of ∼9000 FPKM. Full gene panel is listed in (**Supplementary Table S8)**. To serve as a control for unspecific binding of probes, we included 50 blank barcodes. All the probes were designed and constructed by Vizgen.

### MERFISH 2.0 Sample Preparation

Human postmortem frozen brain tissue was embedded in Optimum Cutting Temperature medium (VWR, 25608-930) and sectioned on a Leica cryostat at −20 °C at 10 um onto MERSCOPE Large Slide V 2.0 slides (Vizgen, 20400118). These sections were then processed for MERSCOPE Sample Preparation according to the manufacturer’s instructions (MERFISH 2.0 Sample Preparation User Guide for Sectioned Tissue Samples, 91600132 Rev A). Once adhered to the coverslip the tissue was fixed followed by three washes with 1× PBS. After aspiration, 70% ethanol was added to permeabilize the tissue for at least 24 h. After a wash with Formamide Wash Buffer (Vizgen, 20300002), the sample was proceeded to anchoring pretreatment, RNA anchoring, and then the tissue was embedded in a polyacrylamide gel followed by incubation with tissue clearing solution overnight at 47 °C. The sample remains in Clearing Solution for autofluorescence quenching. After the tissue became transparent, sample was washed with the wash buffer (Vizgen, 20300001) and then incubated with the custom MERSCOPE 500 Gene Panel V 2.0, and left to hybridize at 47°C overnight. The sample was then washed and incubated at 47 °C with Formamide Wash Buffer twice, and incubated with Enhancer Wash Buffer (PN 20300192) at 47 °C. The samples were stored in the Clearing Premix at 37°C for up to 7 days.

### MERSCOPE Ultra imaging

Clearing solution was removed, and samples were washed for 10 minutes with Sample Prep Wash Buffer. Samples with incubated with 6mL of DAPI and Poly T Reagent for 15 minutes at room temperature, washed for 10 minutes with 10 mL of Formamide Wash Buffer and transferred to 10 mL of Sample Prep Wash Buffer. In the meantime, 100 µl of RNase Inhibitor (New England BioLabs M0314L) was added to 250 µl of Imaging Buffer Activator and this mixture was added via the cartridge activation port to a pre-thawed and mixed MERSCOPE Large 500 Gene Imaging Kit V 2.0 (VIZGEN, #10400173). 15 ml mineral oil (Millipore-Sigma m5904-6X500ML) was added to the activation port and the MERSCOPE Ultra fluidics system was primed according to VIZGEN instructions. The sample was placed in the MERSCOPE Ultra Large Flow Chamber (FCX-L) and connected to the fluidics system of the Vizgen microscope, taking care to disperse air bubbles. A low-resolution mosaic was acquired using a 10X objective, and the regions of interest were selected for high-resolution imaging with a 60x lens. Detected transcripts were decoded on the MERSCOPE Ultra Instrument using a Codebook generated by Vizgen. Nucleus segmentation based on DAPI staining was performed using the build-in MERSCOPE Ultra Instrument Software to generate the output cell-by-gene matrix for downstream analysis.

### MERFISH Data Preprocessing and Annotations

#### Data Loading

The cell-by-gene matrix (.csv format) was loaded and processed as a SeuratObject using a custom function adapted from Seurat (v5.1.0) *ReadVizgen* function, with a minimum cell volume threshold of 1 μm^3^ to exclude segmentation artifacts, and filtering of blank control probes (prefix “Blank-”). Nucleus segmentations were read from parquet files containing polygon boundaries. Only cells with both centroid coordinates and complete segmentation polygons were retained. Cell-level metadata including cell volume (estimated from linear interpolation across 7-z plane measurements), solidity and perimeter-to-area ratio were added from the cell_metadata.csv file. Blank probe counts were calculated separately for quality assessment.

#### Quality control and integration

Cell-level quality control was performed independently for each tissue region on three metrics using quantile-based thresholds with hard cutoffs: (1) transcript count: 5^th^ to 99^th^ percentile with minimum 0 and maximum 4000 transcripts, (2) cell volume: above 5^th^ percentile with minimum 1 μm^3^ and maximum 5000 μm^3^, (3) gene count: 5^th^ to 99^th^ percentile with minimum 30 genes and no upper limit. Cells failing any threshold were excluded from downstream analysis. After these preprocessing steps, 754,760 cells were retained for subsequent analysis.

Quality-filtered cells from all samples were merged and normalized using Seurat’s *SCTransform* with *glmGamPoi* method, regressing out cell volume to correct for transcript count difference due to differences in nuclei size. All genes on the panel were included as variable features. Principal component analysis (PCA) was performed on the SCTransformed data using 100 principal components. To correct for batch effects across samples, data were integrated using Harmony (v) with sample ID as the batch variable, using 100 principal components and default parameters.

#### De novo clustering and cell type annotation

Graph-based clustering was performed on harmony-corrected embeddings using the Leiden algorithm (*FindClusters*, algorithm = 1). To identify optimal clustering resolution, we tested multiple values (0.1, 0.3, 0.5, 0.7, 1.0) and evaluated cluster stability using *clustree* (v0.5.0), which visualizes cluster relationships across resolution parameters. Resolution 0.7 was selected based on minimal cluster instability (few splits/merges between adjacent resolutions) while providing sufficient granularity to separate known cell types based on marker gene expression.

Cluster-specific marker genes were identified using pseudobulk differential expression analysis with *DESeq2* (v1.42.0) on raw transcript counts. For each cluster, single-cell counts were aggregated by cluster and sample to create pseudobulk profiles, with each sample treated as a biological replicate (using Seurat *AggregrateExpression* function). Each cluster was tested against all others using a one-versus-rest design: cluster of interest versus all other cells. Sample variation was modeled through DESeq2’s dispersion estimation across biological replicates. Low-expression genes were filtered (minimum 10 counts in ≥3 pseudobulk samples). Significance was defined as adjusted p-value (*padj*) <0.05 (Benjamini-Hochberg correction) with positive log2 fold-change.

Cell type annotations were assigned using a marker-based scoring system. Each cluster was scored against curated marker gene panels for 13 brain cell types: astrocytes (*AQP4, ALDH1L1, GJA1, FGFR3*), oligodendrocytes (*MOG, OPALIN, OLIG1, OLIG2*), oligodendrocyte precursor cells (*PDGFRA*), microglia (*P2RY12, TMEM119, HEXB, CX3CR1*), macrophages (*CD163, MRC1, LYVE1*), monocytes (*CD14, LYZ*), endothelial cells (*CLDN5, PECAM1, VWF*), pericytes (*PDGFRB, RGS5*), smooth muscle cells (*ACTA2*), fibroblasts (*COL1A1*), excitatory neurons (*SLC17A7, RBFOX3*), inhibitory neurons (*GAD1, GAD2, PVALB, SST, VIP*), and T cells (*CCR7, CD4*). To summarize cluster enrichment of cell-type specific markers, for each cluster-cell-type combination, a weighted enrichment score was calculated based on the differential gene expression statistics: score = ∑(*log*₂*FC* × −*log*₁₀ *p_ad_*_j_) for significant upregulated markers (*padj* < 0.05, log2FC > 0). Scores were normalized (z-scores) across cell types. Clusters were assigned to the top-scoring cell type. Clusters with no significant marker upregulated were labeled “Unassigned”.

#### White matter and gray matter niche annotation

Each sample was separately annotated for white matter (WM) and gray matter (GM) regions based on neighborhood analysis on the cell-type composition, using Seurat function *BuildNicheAssay*. For each cell, the local cellular neighborhood was defined using *k*-nearest neighbors (*neighbors.k* = 200). Niche identity was determined by *k*-means clustering (*niches.k* = 2 for two niches) on neighborhood cell type compositions. Due to variability in tissue morphologies, three of the tissues required specific parameter adjustment (*neighbors.k* = 400 and *niches.k = 6* and then merged into two niches). Niches assigned as WM or GM based on oligodendrocyte enrichment (*reference.cell.type = “Oligodendrocytes”*). WM regions showed high expression of myelin markers (*MOG, OPALIN*), while GM regions showed enrichment for neuronal markers (*SLC17A7, RBFOX3, RASGRF2, RORB*). Annotations were validated by spatial visualization of marker gene expression.

### Ast10 *in situ* identification and hotspot analysis of MERFISH Data

To identify Ast10 *in situ*, individual astrocytes from all tissues were scored by averaging the expression across five Ast10 marker genes (*SLC38A2, SMTN, SLC39A11, CACNA1D*, *MT1E*). In snRNAseq data, these five markers together achieved high sensitivity and specificity in distinguishing Ast10 (area under receiver operating characteristic curve AUC score of 0.88, **Extended Data Figure 8D, E**). For cross-modality comparison of Ast10 frequencies only, we applied a stringent threshold and classified astrocytes within the top 1% of the Ast10 score distribution across all MERFISH tissues as Ast10.

However, all subsequent spatial analyses were performed using the *continuous* Ast10 score rather than this discrete classification to maximize statistical power.

To assess the degree to which astrocytes with high or low Ast10 score cluster spatially, local *Getis-Ord Gi\** statistics^61^ were computed on the continuous Ast10 score using *k*-nearest astrocyte neighbors (*k* = 8) for each tissue section independently. Local *Getis-Ord Gi\** statistics for focal astrocyte *i* (*Gi\**) was calculated as follows:

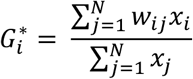

where *xi* is the *continuous* Ast10 score for astrocyte *i* and *wij* is the spatial weight matrix which indicates whether *j* is within the *k^th^* nearest neighbor of *i*. Since *Gi\** is a z-score metric, significantly positive *Gi\** values *(Gi* ≥* 1.96, equivalent to *P-*value ≤ 0.05) indicate hotspots, which are clusters of astrocytes with high Ast10 scores; significantly negative *Gi\** values *(Gi* ≤ -*1.96, equivalent to *P-*value ≤ 0.05) indicate coldspots (clustering of astrocytes with low Ast10 scores).

### Spatial Lag Model

To validate prioritized ligand-receptor pairs that predict the Ast10 state in a spatial context, we applied a multi-sample Spatial Lag of X (SLX) mixed-effects modeling framework to the MERSCOPE data. We restricted the analysis to ligand-receptor pairs in which the ligand was retained as a stable predictor in the final PLSR model and for which both the ligand and its cognate receptor(s) were present in the MERSCOPE gene panel, yielding a final set of 21 pairs. Cognate ligand-receptor pairs were obtained from the NicheNet human ligand-receptor network. For each pair, the model estimated the Ast10 score of a focal astrocyte as a function of four effects: (1) spatially weighted ligand expression in the *k* nearest neighboring microglia or astrocytes (i.e., the spatial lag), (2) receptor expression within the focal astrocyte itself, and (3) focal astrocyte cell volume and the neighborhood total transcript counts as fixed-effect technical covariates, with (4) inter-sample variability modeled as a random effect. Ligand and receptor expression values were log1p-transformed and z-scored per gene within each tissue section prior to spatial lag computation or model fitting. Neighborhood total transcript count was similarly log1p-transformed and spatially lagged, and both technical covariates were z-scored within tissue section. To assess robustness to neighborhood definition, the analysis was repeated across multiple values of *k*.

#### Spatial Weight Matrix Construction

For each tissue section, we used the spatial coordinates of individual astrocytes and microglia to construct row-standardized *k*-nearest-neighbor (KNN) spatial weight matrices in which the weights across the focal cell’s neighbors summed to one. KNN graphs were generated using the *knearneigh* function from the R package spdep (v1.4.1) and converted to row-standardized spatial weights lists using *nb2listw* (*style = “W”, zero.policy = TRUE*).

#### Spatial Lag Computation

For each candidate ligand, the spatially lagged expression was computed as the weighted average of normalized ligand expression across the focal astrocyte’s *k*-nearest neighboring microglia or astrocytes using the *lag.listw* function from spdep. This spatial lag represents the mean ligand expression in the local glial microenvironment of each astrocyte. The spatial lag of total transcript count was computed in the same manner to serve as a fixed-effect technical covariate.

#### Model Specification

For each ligand-receptor pair (*l, r*), a linear mixed-effects model with Gaussian errors was fitted jointly across all samples using the R package glmmTMB (v1.1.14). The model was specified as:

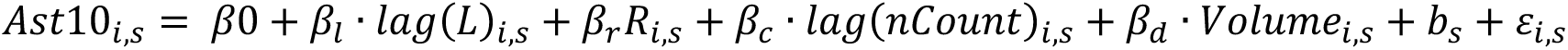

where *lag(L)i,s* is the spatially lagged normalized expression of ligand *l* in the neighborhood of focal astrocyte *i* in sample *s*; *Ri,s* is the normalized expression of receptor *r* in the focal astrocyte *i* of sample *s*; *lag(nCount)i,*s is the spatially lagged total transcript count and *Volumei,s* is cell volume, included as technical covariates; *bs* ∼ *N(0,σi,s)* is a sample-specific random intercept; and *εi,s* is the residual error. Only astrocytes were included as observations (i.e., the response variable was defined for astrocytes only), while spatial lag predictors were computed over the combined astrocyte-microglia pool. Models were fitted using the *glmmTMBControl* function.

#### Sensitivity Analysis

To assess robustness to neighborhood size, the full modeling pipeline was repeated across four values of *k* (k = 3, 6, 9, 12), corresponding to a median interaction radii ranged approximately 59 µm-126 µm. This range was selected to capture biologically distinct signaling scales, from direct cell contact to medium-range paracrine signaling. All other model parameters were held constant across configurations. Multiple testing correction was applied using the Benjamini-Hochberg (BH) procedure, separately within ligand and receptor predictor terms, and independently for each *k*.

### Immunofluorescence microscopy

A six μm formalin-fixed paraffin-embedded (FFPE) tissue section from the dorsolateral prefrontal cortex (Brodmann Area 9) was de-paraffinized with CitriSolv (xylene substitute, Decon Laboratories, 1601H) for 20 min. Heat-induced epitope retrieval was performed with citrate buffer (pH=6) using a microwave (800W, 30% power setting) for 25 minutes. The section was washed with phosphate-buffered saline (PBS) and blocked with a bovine serum albumin (BSA) medium (Sigma-Aldrich, A7906) for 30 minutes at room temperature (RT). The primary antibodies, anti-PLXNB1 (Bio-Techne, MAB3749, 1:100) and ZIP11 (Thermo Fisher, PA5-120717, 1:100), were incubated overnight at 4°C. The tissue section was washed three times with PBS and incubated with fluorochrome-conjugated secondary antibodies (Thermo Fisher, A21202, A31573, 1:500) for 1 hour at RT. After washing with PBS, GFAP-conjugated Cy3 (Millipore Sigma, MAB3402, 1:100) was incubated for 2 hours at RT. After washing, the section was treated with True Black Lipofuscin Autofluorescence Quencher (Invitrogen, P3693) to minimize autofluorescence. An anti-fading reagent with DAPI (Life technology, P36931) was used for tissue mounting.

### Immunofluorescence image acquisition and analysis

Imaging was performed using the Nikon Eclipse Ni-E epifluorescence microscope at 20X magnification with the NIS-Elements Advanced Research software (v5.21.03). A Hamamatsu Orca-Fusion Digital Camera (C14440) was utilized to capture 40 images in a systematic zigzag pattern, ensuring coverage of all cortical layers for post-acquisition analysis. The Plan Apochromatic λ objective lens with a numerical aperture of 0.75 was used for high-performance resolution imaging across a range of wavelengths. The camera was set to a 1X1 binning configuration to enhance signal sensitivity and improve signal-to-noise ratio while maintaining a fast image acquisition. The epifluorescence microscope was equipped with a high color rendering LED light source from a Lumencor system, providing stable and efficient illumination for fluorescence imaging. For multi-channel fluorescence, a Multilaser Laser Induced Detection Area (LIDA) system was employed, integrating multiple lasers, each assigned to specific fluorophores, enabling simultaneous excitation and detection of their emissions.

The excitation lasers used were 450 nanometers (nm), 550 nm, and 640 nm, with individual detection channels for the emitted light. The emission wavelengths were set as follows: DAPI (365 nm), FITC (488 nm), TRITC (561 nm), and Cy5 (640 nm). The exposure time for all images acquired at 20X magnification was 40 milliseconds.

To extract single-cell level protein measurements from the images, astrocytes were segmented using CellProfiler (v4.2.5)^115^ based on DAPI+ and *GFAP*+ objects. First, nuclei (DAPI) were segmented using the “IdentifyPrimaryObjects” module with advanced settings. The object diameter was set between 18 and 80 pixels, and thresholding was performed with the “Robust Background” method, without adjustment. The lower and upper bounds of the threshold ranged from 0 to 1. For *GFAP* segmentation, the “EnhanceOrSuppressFeatures” module was used to augment the ramified features of astrocytes by applying the “Line Structures” method. *GFAP*+ objects were then segmented using the “IdentifyPrimaryObjects” module, with object diameters ranging from 10 to 300 pixels. Thresholding was performed again with the “Robust Background” method without adjustment. The lower and upper bounds of the threshold ranged between 0 to 1. The “SplitOrMergeObjects” module was used to group smaller objects, such as ramified structures, with larger nearby objects. To exclude *GFAP*+DAPI-objects, the “RelateObjects” module defined DAPI as the parent object and *GFAP* as the child object.

The mean intensities of *SLC39A11*, *GFAP*, and *PLXNB1* were measured for each *GFAP*+DAPI+ object. All intensity measurements from CellProfiler were rescaled to a range between 0 to 16 bits by multiplying a factor of 65535 (2^16^ – 1), followed by log10 transformation.

To statistically test for the association between *PLXNB1* expression and Ast10 marker *SLC39A11* expression at the single-cell level, we fit a mixed effects linear regression model. The model predicts *SLC39A11* mean intensity based on *PLXNB1* mean intensity using all astrocytes (*GFAP*+DAPI+ cells). *GFAP* mean intensity was included as a covariate, while random effects accounted for variability in both intercepts and slopes for *GFAP* and *PLXNB1* mean intensities, as well as variability across images. All continuous variables were z-score normalized prior to model fitting. The model was implemented using the *lmer* function from the lme4 package (v1.1-35.5) in R. The structure of the model is as followed:

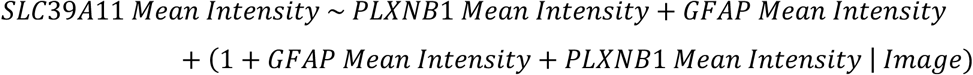

Here, *PLXNB1* mean intensity and *GFAP* mean intensity were modeled as fixed effects; the terms in parentheses represent random effects, allowing intercepts and slopes for *GFAP* and *PLXNB1* to vary across images. This mixed-effects modeling approach accounts for both single-cell-level relationships and image-level variability, providing robust estimates of the association between *PLXNB1* and *SLC39A11* expression.

### Generation of isogenic hiPSC pairs with or without *PLXNB1* knockout

Two isogenic pairs of wild-type (WT) and *PLXNB1*-knockout (*PLXNB1-KO*) human induced pluripotent stem cell (hiPSC) from two control donors (CD1: 553, CD2: 2607) were generated using CRISPR/Cas9, as previously described^64^. Three guide RNAs (gRNAs) targeting *PLXNB1* exon 3 were obtained through the Mount Sinai CRISPR Initiative Program. **Table 1** provides details on the gRNA sequences.

**Table 1.**
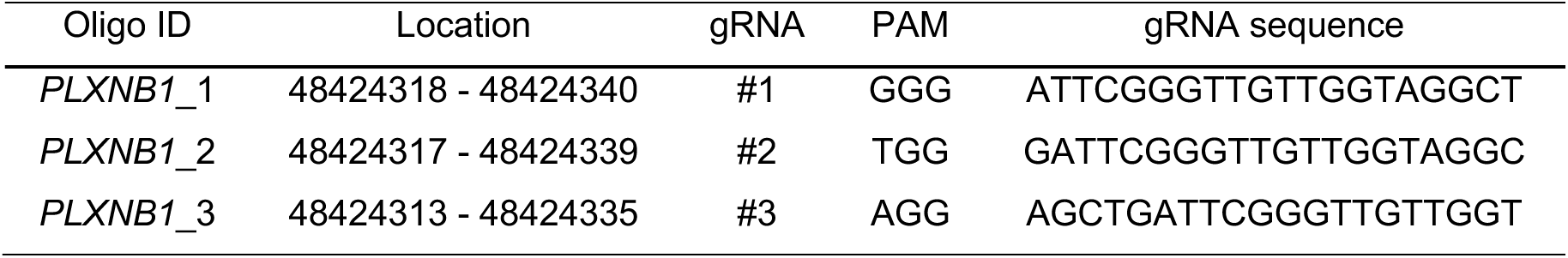
Guide RNAs (gRNAs) used for generating *PLXNB1* KO hiPSCs.

**Table 2.** Cell inclusion criteria for quality control of scRNAseq data from hiPSC-derived astrocytes.

| iAst | UMI count range | Gene count range | MT% threshold |
| --- | --- | --- | --- |
| Line 553 | 2,000 – 50,000 | 2,300 – 7,000 | ≤10% |
| Line 2607 | 2,500 – 50,000 | 2,300 – 7,000 | ≤10% |

### Generation of hiPSC-derived astrocytes

The isogenic pairs of WT and *PLXNB1*-KO hiPSCs (from single clones) were induced into astrocytes by expressing FUW-M2rtTA (Addgene, 20342), TetO.Sox9.puro (Addgene, 117269), and TetO.NfiB.Hygro (Addgene, 117271), as previously described^64,66,116^. On day 0, add doxycycline (Dox) to induce *SOX9* and *NFIB* expression. On day 1, add dual antibiotics (puromycin and hygromycin) to enrich transduced hiPSCs, then expand cells in Expansion Medium (DMEM/F-12, HEPES, N2, 10% FBS, GlutaMAX). On day 3, switch to Astrocyte FGF Medium (Neurobasal, B27, GlutaMAX, CNTF, BMP4, FGF2) with Dox. On day 8, transition to Astrocyte Maturation Medium (DMEM/F-12, Neurobasal, N2, sodium pyruvate, EGF-like growth factor, CNTF, BMP4, cAMP, NAC) with Dox. On day 11, fully withdraw Dox. Collect hiPSC-derived astrocytes between days 21-24.

### scRNA-seq of hiPSC-derived astrocytes

iAst were collected and filtered through a 70 µm cell strainer into a fresh falcon tube. The viability of single cells was assessed using Acridine Orange/Propidium Iodide viability staining reagent (Nexcelom), and debris-free suspensions of >80% viability were deemed suitable for the experiments. scRNA-seq was performed using the Chromium platform (10x Genomics) with the 5’ gene expression (5’ GEX) v2 kit, with a targeted recovery of 10,000 cells. Briefly, Gel-Bead in Emulsions (GEMs) were generated on the sample chip in the Chromium X system. Barcoded cDNA was extracted from the GEMs after Post-GEM RT-cleanup and amplified for 12 cycles. Full-length cDNA from poly-A mRNA transcripts underwent enzymatic fragmentation and size selection to optimize cDNA amplicon size (∼400 bp) for library construction. The cDNA fragments were then subjected to end-repair, adapter ligation, and 10x-specific sample indexing according to the manufacturer’s protocol (10x Genomics). The concentration of the single-cell library was accurately quantified using qPCR (Kapa Biosystems) to achieve appropriate cluster counts for paired-end sequencing on the NovaSeq 6000 (Illumina), aiming for a sequencing depth of 25,000 reads per cell.

### Preprocessing scRNA-seq data from hiPSC-derived astrocytes

Sequencing reads were aligned to the human reference genome (GRCh38) using Cell Ranger (v 5.0.1). Starting from the raw UMI count matrices generated by Cell Ranger, we performed quality control separately for each iAst line by retaining cells that met line-specific thresholds for UMI count and gene count, and had a mitochondrial transcript percentage of ≤ 10%:

### Ast10 co-expression structure in hiPSC-derived astrocytes

To determine whether the Ast10 signature gene co-expression structure observed in human brain tissue is preserved in iAst, we computed Pearson correlation matrices across Ast10 signature genes (upregulated genes only) using MAGIC-derived expression values from wild-type cells only, separately for each iAst line. For each line, pairwise Pearson correlations were calculated for all available Ast10 genes; genes absent from either line were excluded. The resulting per-line correlation matrices were then averaged element-wise to generate a final co-expression matrix for clustering. Gene-gene co-expression modules were identified by hierarchical clustering of the averaged correlation matrix using Pearson distance (1-*Pearson’s R*) and complete linkage (*hclust*, *method = “complete”*). The dendrogram was cut at *k* = 2 to define two gene sets. For comparison with post-mortem astrocytes, we similarly computed Pearson correlation for each gene pair from the available list of Ast10 upregulated genes, using MAGIC-derived expression values of the *CUIMC1* snRNAseq dataset (See *Gene-gene co-expression assessment)*. Overall, MAGIC here is only used for understanding the underly gene-gene correlation structures; however, evaluation of the knockout effect using differential gene expression analysis was performed on raw counts.

### Preprocessing scRNA-seq dataset of mice with or without Plxnb1 knockout

We used Seurat to preprocess the raw UMI count matrices generated by Huang et al using single-cell RNA sequencing of cortices from mice with or without *Plxnb1* deletion^64^. For quality control, we removed cells with ≤200 or ≥ 5000 genes, UMI count ≥ 50,000, as well as those with mitochondrial reads > 10%. We then removed genes that were expressed in less than 2 cells. After quality control, we retained 32,874 cells and 20,557 genes in total.

### Annotating cell-types to identify astrocytes in mouse scRNA-seq data

To annotate cell types and identify astrocytes in the scRNA-seq datasets, we used the Discovery dataset as reference and applied the same Seurat mapping approach^39^ similar to the cell-state annotation process mentioned above. To map the mouse data onto the reference, which is based on human transcriptomes, we first converted all mouse genes to human genes using the *convert_mouse_to_human_symbols* function from the nichenetr package. To annotate the major cell types, we normalized each dataset using *SCTransform* (with *variable.features.n* = 4000 and 1000 for the mouse data and iAst data, respectively). Transfer anchors were identified using the *FindTransferAnchors* function with the 10 nearest neighbors (*k.anchor* = 10) and the first 30 principal components (*npcs* = 30), while keeping the other parameters as default. To transfer the cell-type labels, the anchors were then input to the *TransferData* function, using the 10 nearest neighbors for weighting anchors (*k.weight* = 10) and *npcs* = 30. Together, the preprocessing and annotation pipelines result in 3688 astrocytes, with 1,240 wild-type astrocytes and 2,448 with *Plxnb1* knock-out. These annotated astrocytes were then selected for down-stream analysis.

### Differential expression analysis to assess *PLXNB1*-knockout effect in scRNAseq data

#### Sequencing Depth Equalization

To minimize systematic differences in sequencing depth between WT and *PLXNB1*-knockout conditions within each line or species, we compared median UMI counts between conditions. For each comparison, the condition with the higher median UMI count was probabilistically downsampled on a per-cell basis to match the target depth, defined as the median UMI count of the shallower condition, using the *downsampleMatrix* function from the DropletUtils package (v1.30.0). After downsampling, the median UMI counts were 12,246 for line 553 and 9,654 for line 2607. We applied the same procedure to all astrocytes in the mouse dataset, resulting in median UMI counts of 2400. Because the cell count imbalance between conditions (2:1) was substantially larger in the mouse dataset than in the iAst dataset, mouse astrocytes were additionally randomly downsampled to equal cell counts per condition (1,240 cells each) prior to differential expression analysis. No such cell-count balancing was applied to the iAst dataset, where WT and KO cell counts were comparable across lines. These depth-equalized count matrices were used as input for downstream differential expression analyses.

#### Differential Gene Expression

To test whether the Ast10 transcriptional program is systematically downregulated in *PLXNB1*-knockout astrocytes, we performed single-cell differential expression analysis on the raw count data using Nebula^117^ (v1.5.6). Negative binomial mixed-effects models were fitted to depth-equalized counts with library size offsets defined as the log of total UMI counts per cell, computed from the full transcriptome. Models were fitted using *method = “LN”* and gene filter based on counts per cell <0.1% (*cpc* = 0.001).

For iAst, we jointly analyzed two independent isogenic lines derived from distinct genetic backgrounds, using *∼ condition + line*, with line as a fixed effect and sample as the subject-level random effect (*id = sample_id*). This model provided the primary estimates of gene-level log2 fold changes (PLXNB1-KO versus WT) and *P*-values while accounting for within-sample cell-level correlation. We additionally performed separate analyses within each line (∼ condition) to assess the consistency of effect directions across genetic backgrounds. Because each line contained only one sample per condition, these per-line analyses were used to assess directional concordance rather than statistical significance. We additionally performed separate analyses within each line (∼ condition) to assess the consistency of effect directions across genetic backgrounds. Because each line contained only one sample per condition, these per-line analyses were used to assess directional concordance rather than statistical significance.

For murine astrocytes, which included only one sample per condition, we performed differential expression analysis using Nebula with the same modeling framework as in the per-line iAst analysis, specifying *sample* as the subject-level variable and using the formula *∼ condition* with the same parameter settings.

### Gene set enrichment analysis to assess *PLXNB1*-knockout effect

Since our primary hypothesis concerned a coordinated shift in the Ast10 transcriptional program following *PLXNB1* knockout, rather than differential expression of individual genes, we based the main inference on gene-set enrichment rather than on standalone per-gene significance. In the mouse astrocytes and iAst analyses, genes were ranked by the Wald statistic estimated from each Nebula model, providing a signed measure of differential expression that reflects both the estimated effect and its standard error. Genes upregulated in *PLXNB1*-knockout cells were placed at the top of the ranked list and genes downregulated in *PLXNB1*-knockout cells were placed at the bottom. Gene set enrichment analysis was performed separately for Ast10 Gene Set 1 and Ast10 Gene Set 2, using the *fgseaMultilevel* function (*eps* = 0 and default parameters) from the *fgsea*^118^ R package (v1.36.2).

### Ligand stimulation experiment

One million human primary astrocytes (ScienCell #1800) were thawed onto a T75 flask that was pre-treated with 2 µg/cm^2^ poly-L-lysine in complete astrocyte medium (ScienCell #1801) according to the manufacturer’s instructions, and media replaced once every 2-3 days. After reaching ∼90% confluence (2-4 days post-thaw) cells were dissociated using 0.05% Trypsin/EDTA solution (Gibco) and plated at 50,000 cells/cm^2^ on a 12-well plate that was pre-treated with 2 µg/cm^2^ poly-L-lysine. Once reaching 90% confluence (2-3 days post-plating) cells were treated for 24 hours with SEMA4D (1 µg/mL, R&D Systems 7470-S4-050), JAM2 (5 µg/mL, R&D Systems 1074-VJ-050), a TNF/IL1B cocktail (TNF = 1 ng/mL, R&D Systems #210TA020; IL1B = 5 ng/mL, R&D Systems 201LB005), or treatment/vehicle combinations thereof (see Figure 6H-I). Vehicles for the treatments were as follows: water (SEMA4D vehicle), IgG1 (JAM2 vehicle, 5 µg/mL, R&D Systems #110-HG-100), 0.1% Bovine Serum Albumin (BSA) + PBS (TNF/IL1B vehicle). Cells were lysed in 200 µL RNA lysis buffer (Invitrogen) + 1% B-mercaptomethanol for 2 minutes on ice, followed by neutralization with 200 µL 70% ethanol. RNA was extracted using PureLink RNA Mini Kit (Invitrogen) for downstream bulk RNA sequencing.

### Bulk RNA sequencing of primary astrocytes

#### Library preparation and sequencing

Total RNA was extracted from primary human astrocytes and enriched for polyadenylated mRNA by poly-A pulldown. Strand-specific RNA-seq libraries were prepared using the Illumina TruSeq Stranded mRNA library preparation kit with KAPA HiFi HotStart Ready Mix for the final PCR amplification step. Libraries were sequenced on the Element AVITI platform at the Columbia Genome Center as paired-end 75 bp reads, targeting 40 million read pairs per sample. Base calling and adapter trimming were performed using bases2fastq (v2.2.0).

#### Transcript quantification

Raw sequencing reads were pseudoaligned to a kallisto index (v0.50.1) built from human protein-coding transcriptome sequences (Ensembl release 113, GRCh38.p14, cDNA only) using default parameters. All 24 samples achieved pseudoalignment rates between 82.5% and 86.0% (mean 84.5%). Transcript-level estimated counts were imported and summarized to gene-level counts using the *tximport* R package with the ‘lengthScaledTPM’ method.

### Differential expression analysis to assess ligand stimulation effect in primary astrocytes

To identify genes whose expression changed in response to ligand stimulation, we performed differential expression analysis using DESeq2 (v1.38.0). DESeq2 operates on raw estimated counts, modeling count data using a negative binomial distribution to account for the overdispersion typical of RNA-seq data. We performed three sets of pairwise comparisons (n = 3 per group): cytokine cocktail stimulation alone versus vehicle control (TNF /IL1B vs. vehicle); ligand treatment versus vehicle control (ligand vs. vehicle); and ligand treatment in combination with cytokine cocktail versus cytokine cocktail alone (TNF /IL1B + ligand vs. TNF /IL1B), the latter designed to isolate ligand-specific transcriptional effects in the context of cytokine-induced astrocyte reactivity. Size factors were estimated to normalize for differences in sequencing depth across samples, and gene-wise dispersion was estimated by sharing information across genes using DESeq2’s empirical Bayes shrinkage procedure. Differential expression was tested using the Wald test, which evaluates whether the log2 fold change between groups differs significantly from zero. Raw p-values were adjusted for multiple comparisons using the Benjamini-Hochberg method to control the false discovery rate.

### Gene set enrichment analysis to assess ligand stimulation effect

For each differential expression comparison, genes were ranked using a signed composite statistic calculated based on DESeq2 Wald statistic × -log10(*P*-value), which prioritizes genes according to both the direction of differential expression and the strength of statistical support. Genes with missing *P-*values or Wald statistics due to low counts or outlier filtering were removed before ranking. Gene set enrichment analysis was then performed using the fgsea package (v1.36.2), with the Ast.10 upregulated signature (see *Identification of cell state signatures*) as the gene set of interest. For each comparison, normalized enrichment scores (NES) and Benjamini-Hochberg-adjusted p-values (*FDR*) reported by fgsea were obtained.

### Association between Ast10 frequency and cognitive decline adjusting for neuropathologies

To evaluate the extent to which the association between Ast10 frequency and cognitive decline can be explained by other factors, we conducted a linear regression analysis. Cognitive slope was modeled as a function of Ast10 frequency alongside neuropathological and demographic covariates available in either the Discovery or Replication cohorts (n = 867 participants). Included covariates were age, sex, education, study cohort, and neuropathological measures: amyloid, tau, Lewy bodies, TAR DNA-binding protein 43 (TDP-43), gross chronic infarcts, chronic microinfarcts, cerebral atherosclerosis, arteriolosclerosis, cerebral amyloid angiopathy, hippocampal sclerosis, and neuron proportion estimated from snRNA-seq (see Methods: *Other neuropathological measures* for details).

Participants with complete data across all variables (n = 687) were included in the final model. Continuous variables were z-score normalized, and categorical variables were dichotomized for model parsimony. Linear regression was performed using the *lm* function from the stats package (v4.4.2) in R.

### Mediation analysis of synaptic proteins in the association between Ast10 frequency and cognitive decline

Synaptic protein levels were measured using Tandem Mass Tag (TMT) multiplexed mass spectrometry in a subset of participants (n = 687) from the CUIMC1, CUIMC2 and MIT datasets. TMT data were preprocessed and normalized as previously described^119^. To assess whether synaptic proteins mediate the relationship between Ast10 frequency and cognitive decline, we performed mediation analysis across proteins that are part of the SNARE-complex (CPLX1, CPLX2, CPLX3, CPLX4, SNAP25, STX1A, STX1B, STX3, VAMP1, VAMP2). For a given mediator protein, we fit three linear models, each representing an “edge” of the mediation triad:

1. Total effect: Association between Ast10 frequency and cognitive decline without adjusting for the mediator protein.
2. Mediator model: Association between Ast10 frequency and the mediator protein levels.
3. Outcome model: Association between the mediator protein levels and cognitive decline, adjusting for Ast10 frequency.

All models were adjusted for sex, postmortem interval, dataset, age at death, education, cell count, and median gene count per participant. The *mediate* function from the mediation package (v4.5.1) in R was used with non-parametric bootstrapping (5,000 simulations) to estimate the proportion of the effect mediated by each synaptic protein.

### Direction Dependence Analysis of Synaptic protein – Ast10 association

Since mediation analysis assumes a directional relationship between the exposure and mediator, we used direction dependence analysis (DDA) to test whether the association between Ast10 frequency and each synaptic protein was better supported in the Ast10 → protein direction or the reverse protein

◊ Ast10 direction. DDA was performed using the *dda.indep* function in the dda R package. Ast10 frequency was square-root transformed, consistent with the primary cell-state association analyses.

For each protein, the target model specified Ast10 → protein and the alternative model specified protein → Ast10. Both variables were adjusted for the same covariates used in the mediation analysis. Directional evidence was assessed using distance correlation (dCor), which quantifies dependence between the predictor and regression residuals. We calculated the difference in dCor between the alternative and target models (alternative minus target) and estimated its 95% confidence interval using 1,000 bootstrap resamples. A positive difference indicates lower predictor-residual dependence in the Ast10 → protein model and supports this direction, whereas a negative difference supports protein → Ast10. Direction was assigned when the 95% confidence interval excluded zero; intervals spanning zero were considered inconclusive.

Proteins were considered consistent with the proposed pathway when DDA supported Ast10 → protein and the protein showed a significant mediation effect (*P* < 0.05) on the association between Ast10 frequency and cognitive decline.

### Pepinemab SIGNAL-AD Trial Design and cognitive measure

The double-blind, placebo-controlled SIGNAL-AD (NCT04381468) study enrolled 50 individuals with MCI or mild AD dementia in two parallel treatment cohorts conducted at 16 medical research centers and hospitals in the United States of America. Key inclusion criteria: Mini-Mental State Examination (MMSE) ranged between 17-26, Global Clinical Dementia Rating of 0.5 or 1.0 and amyloid positive status, as determined by positron-emission tomography (PET) scan or cerebrospinal fluid (CSF) measurements; both male and female participants were eligible for enrollment. Key exclusion criteria: electrocardiogram abnormalities, prior treatment with monoclonal antibody within 180 days or 5 half-lives, > 4 cerebral microhemorrhages or superficial siderosis. Subjects who satisfied all eligibility criteria were participants and were randomized to one of two treatment arms through an interactive web response system. The subjects, site investigators, site personnel and study statisticians, as well as representatives of these organizations and staff at Vaccinex, were blinded as to treatment assignments until database lock. The investigational agent and placebo were in vials of identical appearance^93^.

Treatment was randomized 1:1 to pepinemab (40 mg/kg) or placebo (every four weeks, 60 minute intravenous infusion) for 12 months. The primary objective of the study is safety and tolerability. Additional secondary and exploratory assessments include pharmacokinetics and pharmacodynamics (PK/PD), fluid biomarkers including multiplex O-link proteomics of CSF, Fluorodeoxyglucose Positron Emission Tomography (FDG-PET) to measure brain metabolism, and change in standard individual cognitive scales, Clinical Dementia Rating Scale-Sum of Boxes (CDR-SB), Alzheimer’s Disease Assessment Scale-Cognitive Subscale 13-item (ADAS-Cog13), Integrated Alzheimer’s Disease Rating Scale (iADRS) and MMSE. By their definitions, higher CDR-SB and ADAS-Cog13 scores indicate greater cognitive impairment, whereas higher iADRS and MMSE indicate better cognitive function. CSF was collected by lumbar puncture at baseline and end of treatment. Cognitive assessments were evaluated at baseline, 12, 24, 36, and 48 weeks.

Institutional review board (IRB) approvals for the study protocol, amendments and informed consent documents were obtained before use in the study; written informed consent was obtained from study participants before the initiation of study procedures. This study utilized both a central IRB – Western Institutional Review Board - as well as local IRBs at sites that did not utilize the central IRB. This study was conducted in accordance with the Declaration of Helsinki, International Conference on Harmonization guidelines and applicable portions of the United States Code of Federal Regulations.

The ClinicalTrials.gov identifier (no. NCT04381468) was obtained before study initiation. All experiments including human specimens were performed in compliance with the relevant ethical regulations. An independent unblinded Data Safety Monitoring Board (DSMB) met with Vaccinex and study medical monitors regularly throughout the trial; no serious adverse events (SAEs) related to treatment were observed and no safety trends emerged. For sample size determinations, for the primary objective of safety, sample size was not determined based on power calculations but instead was chosen based on empirical evidence from other Phase 1 AD trials. The study protocol did not enroll nor discriminate on the basis of sex or gender. Funding for SIGNAL-AD was provided by Vaccinex, Inc., Alzheimer’s Association, and Alzheimer’s Drug Discovery Foundation.

### Olink CSF Measurements

CSF and plasma samples were processed and stored in accordance with international consensus guidelines. For each sample, a fresh aliquot was used, i.e. without previous freeze-thaw cycles. Sample size at baseline included 26 patients in placebo group and 23 participants in pepinemab group, with 16 matched pairs containing baseline and end of treatment CSF samples in placebo group and 17 matched pairs in pepinemab group. Proteomics analyses were performed using validated multiplex panels based on Proximity Extension Assay (PEA) technology (Olink Proteomics Inc.). Briefly, target proteins are bound by unique pairs of oligonucleotide-labelled antibodies. When in close proximity to one another, the oligonucleotides hybridize to form a Polymerase Chain Reaction (PCR) target sequence. The resulting DNA amplicon is quantified on an Illumina NovaSeq platform using Next Generation Sequencing (Olink Explore 5000). Quality control samples and negative controls were supplied by the manufacturer. For each protein, the lower limit of detection (LOD) was defined as the sample of blanks plus 3 standard deviations. Proteins were included in further analyses if they were above the LOD in >85% of samples and passed quality control. Extreme outliers, defined as Normalized Protein eXpression (NPX) values above or below a *z*-score of 5, were capped at a *z*-score of ±5. Data were evaluated with PCA plots of the participants; no obvious participant sample outliers were identified or removed. Of the 5420 measured proteins, 940 were above the LOD in >85% of CSF samples and were included in the analyses. Only CSF biomarker data relevant to the Ast10-associated protein signature are analyzed in this manuscript; clinical efficacy and full trial outcomes are not the primary focus of this study.

### Calculation of Ast10 z-scores using CSF proteomic measurements

To derive a CSF proteomic measure as a proxy for Ast10 enrichment, we intersect the 940 proteins measured in the Olink CSF panel with the Ast10-associated protein set, which includes genes uniquely upregulated in Ast10 from single-nucleus gene expression data (see *Identification of cell state signatures*), along with prioritized ligands associated with Ast10 enrichment from the final PLSR model (**Figure 3I**). Six overlapping proteins were retained for score calculation: JAM2, BCAN, NCAN, GNAS, RGMA and EDNRB. These included prioritized ligands predictive of Ast10 enrichment (JAM2, BCAN, NCAN, GNAS and RGMA) and Ast10-specific markers (JAM2, EDNRB and BCAN).

For each retained protein, expression (NPX) values were normalized relative to the baseline reference distribution across all participants at Week 0. Specifically, for each protein, we calculated the median and median absolute deviation (MAD) of its Week 0 values and used these to compute a robust standardized value of the given protein for each sample:

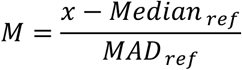

where *x* is the observed expression value for a given protein in a given sample, *Medianref* and *MADref* are the median and MAD, respectively, of the Week 0 reference distribution for that protein. In the implementation, MAD was calculated using the *mad* function in R with the normal-consistency scaling (*constant* = 1.4826).

The Ast10 CSF z-score for each sample was defined as the mean of these protein-level standardized values across all retained Ast10-associated proteins, thereby assigning equal weights to each protein in the composite score. Protein selection, score construction and downstream analyses were performed blinded to treatment assignment.

### Assessment of treatment effects on individual Ast10-associated proteins in the CSF

To assess treatment effects on individual Ast10-associated proteins in the CSF, we analyzed participants with paired Week 0 and Week 48 samples. For each protein, we calculated a change score (Week 48 minus Week 0) using the same MAD-standardized values used to construct the composite Ast10 CSF score. These within-subject change scores were then compared between treatment groups using two-sample t-tests, and the magnitude of the between-group difference in change was summarized using Cohen’s *d*. *P-*values were adjusted across proteins using the Benjamini–Hochberg false discovery rate procedure.

### Associations between baseline Ast10 and subsequent cognitive change

To examine whether the baseline CSF Ast10 z-score was associated with subsequent cognitive decline, and whether this association differed by treatment group, we calculated Spearman’s correlation between baseline Ast10 z-score (Week 0) and change in cognitive performance (Week 48 − Week 0), separately within each treatment group. Four pre-specified cognitive outcomes were evaluated: CDR-SB, ADAS-Cog13, iADRS, and MMSE. To allow a consistent directional interpretation across measures, change scores were oriented such that a negative Spearman’s coefficient indicated greater cognitive worsening. For iADRS and MMSE, where lower scores reflect greater impairment, change scores were used as calculated. For CDR-SB and ADAS-Cog13, where higher scores reflect worse performance in their original definitions, change scores were multiplied by -1 before analysis. 95% confidence intervals were estimated using percentile bootstrap with 2,000 resamples.

### Statistical information

Statistical methods, covariates, and procedures for multiple-testing correction are described in the relevant Methods sections. Statistical significance was assessed at *P* < 0.05, with Benjamini-Hochberg false discovery rate (FDR) correction applied where indicated. Empirical *P* values were calculated using reference distributions or permutation tests, as described for the corresponding analyses. All boxplots highlight the median, first and third quartiles, with whiskers extending to 1.5 times the interquartile range. Sample sizes and are provided in the figure legend.

## Author contributions

P.L.D. and N.C.-L. designed the study; N.C.-L. performed the computational- and statistical analyses, with the guidance from P.L.D., V.M., M.F., H.U.K.; S.E.H. performed the ligand stimulation experiment, with the supervision of T.Y.P.; Y.Z. prepared the single nucleus libraries, performed sequencing and generated the MERSCOPE data; M.F., K.C. performed the sequence alignment and demultiplexing analysis; J.Z., B.H., S.L. and M.W. generated isogenic pairs of *PLXNB1* knockout hiPSCs-derived astrocytes and performed the single-cell RNA-seq, with the supervision from K.J.B., B.Z., H.Z., R.H.F., and A.L.; M.T, T.L. performed immunohistochemistry experiment and image analysis; M.T., H.U.K. generated the ST data; E.E. and M.Z. conceived and supervised the SIGNAL-AD trial; Olink data collection, curation, and replication of statistical analysis were performed by T.L.F., V.M.. and L.B..; E.E. and T.F. had unrestricted access to all clinical trial data. N.C.-L., H.U.K analyzed the ST data. N.C.-L. and P.L.D. wrote the manuscript, and all co-authors edited it for critical comments. All authors agreed to submit the manuscript, read and approved the final draft and take full responsibility of its content, including the accuracy of the data and the fidelity of the trial to the registered protocol and its statistical analysis. D.A.B. is the PI of the parent ROS, and MAP studies and obtained funding and performed study supervision. J.S. is the PI of the parent LATC and Rush Clinical Core studies and obtained funding and performed study supervision. L.L.B is the PI of the parent study MARS and obtained funding and performed study supervision. All authors critically reviewed the manuscript. P.L.D. obtained funding for the project.

## Data Availability

All data and analysis output are available via the AD Knowledge Portal (https://doi.org/10.7303/9618104). The AD Knowledge Portal is a platform for accessing data, analyses, and tools generated by the Accelerating Medicines Partnership (AMP-AD) Target Discovery Program and other National Institute on Aging (NIA)-supported programs to enable open-science practices and accelerate translational learning. The data, analyses and tools are shared early in the research cycle without a publication embargo on secondary use. Data is available for general research use according to the following requirements for data access and data attribution (https://adknowledgeportal.org/DataAccess/Instructions). Data can be access from the Synapse database (https://www.synapse.org): for *CUIMC1* snRNA-seq data (syn31512863); for *MIT* snRNA-seq data (syn52293417); for AMP-AD 2.0 *Diversity* snRNA-seq (syn52339332); for *CUIMC2* snRNA-seq data (syn64769520); for 10x Visium spatial transcriptomic data (syn62109935); for *PLXNB1*-KO mice single-cell RNA-seq data (syn26448163); for ROSMAP proteomic data (syn51150434); for all other data, including MERSCOPE raw data, *PLXNB1*-KO iAst single-cell RNA-seq data, ligand stimulation bulk RNA-seq, and immunofluorescence raw images (syn64778723). RADC resources can be requested at https://www.radc.rush.edu.

## Code Availability

All code used in this study are available on GitHub: https://github.com/cu-ctcn/Ast10_Communication

## Declaration of generative AI in the writing process

During the preparation of this work, the authors used ChatGPT-5 to assist with language editing, grammar and readability. After using this tool or service, the authors reviewed and edited the content as needed and take full responsibility for the content of the manuscript.

## Sex as a biological variable

Donor sex was obtained from cohort metadata and is reported in the demographic summaries. Sex was included as a covariate in primary association models where applicable. This study was designed to identify disease-associated cellular states and multicellular signaling programs rather than sex-specific mechanisms and was not powered for definitive sex-stratified inference. We therefore account for sex in relevant models but do not draw conclusions about sex-specific effects.

## Acknowledgements

We thank the individuals who have generously donated their brains to research through the Rush University Alzheimer’s Disease Center. The five RADC cohorts are supported by P30AG10161, P30AG72975, R01AG15819, R01AG17917, U01AG46152, U01AG61356, R01AG067482 and R01AG22018. This work is supported by the National Institutes of Health’s National Institute on Aging (NIH-NIA), through the following grants: Accelerating Medicines Partnership -Alzheimer’s Disease Target Discovery and Preclinical Validation (AMP-AD, U01AG061356), the Alzheimer’s Disease Sequencing Project (ADSP) Functional Genomics Consortium (FunGen-AD, U01AG072572), R21AG077168 (to M.W. and A.L.), RF1AG077828 (to R.H.F., H.Z., and M.W.), U01AG046170, RF1AG057440, R01AG068030, RF1AG074010 (to B.Z.). This work is also supported by the Alzheimer’s Association, through the following grants: AARF-24-1313800 (to. N.C.-L.), AARG-22-928419 (to A.L. and M.W.), ADSF-21-816675 (to H.U.K. and M.K.) and by the New York State Department of Health DOH01-C38330GG and DOH01-C39068GG (to H.Z.). Vaccinex sponsored the SIGNAL-AD clinical trial with partial funding provided by the Alzheimer’s Association and Alzheimer’s Drug Discovery Foundation. This research was funded in part through the NIH/NCI Cancer Center Support Grant P30CA013696 and used the Genomics and High Throughput Screening Shared Resource. We also thank Drs. Zhihong Chen, Seunghee Kim-Schulze, and other members of the Human Immune Monitoring Center (HIMC) at Icahn School of Medicine at Mount Sinai for single-cell RNA sequencing.

## Supplementary Tables

Supplementary Table S1. – Dataset demographics, Cell-state frequencies across studies, Meta-analysis results

Supplementary Table S2 – Astrocytic state signature genes, Pairwise differential gene expression results

Supplementary Table S3 – NicheNet results

Supplementary Table S4. – PLSR ligand importance, model performance, loadings and scores

Supplementary Table S5 – PLSR receptor importance, model performance, loadings and scores

Supplementary Table S6 – Prioritized ligand pathway genes and DREMI scores

Supplementary Table S7. – Visium data demographics, Spatial correlation results and distance sensitivity analysis, MISTy results

Supplementary Table S8. – MERSCOPE donor demographics and Ast10 Getis-Ord G* summary statistics. Supplementary Table S9 – Raw CellProfiler analysis output

Supplementary Table S10. – Differential gene expression results from iAst scRNA-seq data

Supplementary Table S11 – Differential gene expression results from bulk RNA-seq of human primary astrocytes.

Supplementary Table S12. – Cognition model data and results; mediation analysis results

## Supplemental Figures

**Extended Data Figure 1.**
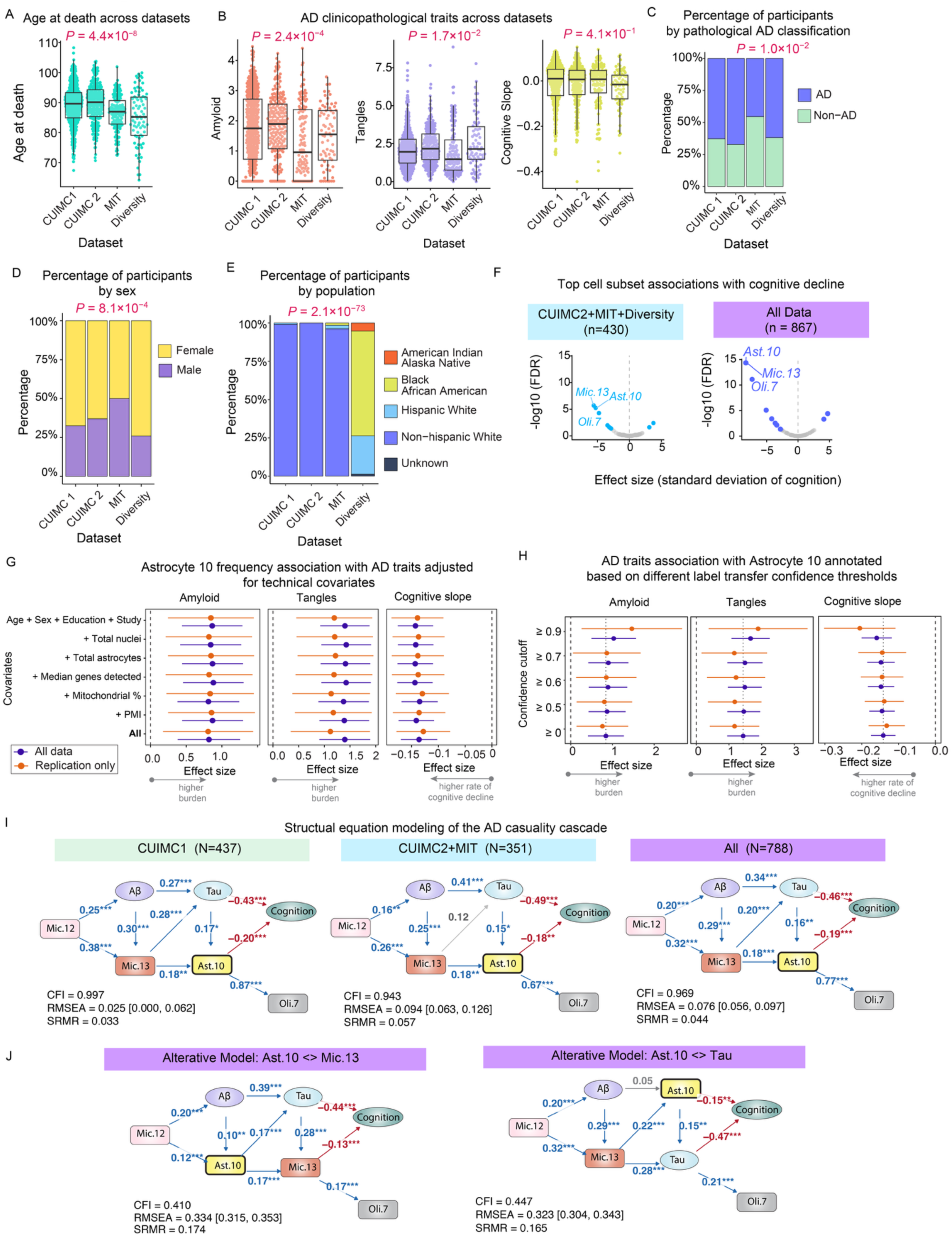
Participant characteristics and meta-analysis of cell subset associations with cognitive decline. **(A–E)** Clinicopathologic characteristics of the 867 participants across the four snRNA-seq datasets. Additional details are provided in Table 1. **(F)** Effect sizes and statistical significance of associations between 91 cell subpopulations and cognitive decline, assessed via meta-analysis of the Replication set (Replication, n = 430) and all datasets combined (All, n = 867). Significant cell subpopulations (False Discovery Rate < 0.05) are highlighted, with the three most significant subpopulations indicated. **(G)** Sensitivity of Ast10 associations with AD traits to technical and demographic covariate adjustment. Forest plots show meta-analysis effect sizes for associations between Ast10 frequency and amyloid burden, tau tangles, and cognitive slope in the full dataset (purple) and Replication datasets only (orange). Models were adjusted for age, sex, education, and study, with additional adjustment for total nuclei, total astrocytes, median genes detected, mitochondrial read percentage, or post-mortem interval (PMI), individually or jointly (“All”). Dots indicate effect-size estimates and error bars indicate 95% confidence intervals. Positive effects for amyloid and tangles indicate higher pathology burden; negative effects for cognitive slope indicate faster cognitive decline. **(H)** Sensitivity of Ast10 associations with AD traits to cell-state label-transfer confidence score. Ast10 annotations in the replication datasets were restricted using progressively more stringent label-transfer confidence thresholds (≥0, ≥0.5, ≥0.6, ≥0.7, and ≥0.9), and associations with amyloid burden, tau tangles, and cognitive slope were re-estimated with meta-analysis. Results are shown for the full dataset (purple) and Replication datasets only (orange). Dots indicate effect-size estimates and error bars indicate 95% confidence intervals. Positive effects for amyloid and tangles indicate higher pathology burden; negative effects for cognitive slope indicate faster cognitive decline. **(I)** Structural equation model of the AD cellular cascade proposed by Green et al, applied across CUIMC1, CUIMC2+MIT and the combined cohort. The model places Mic.12 upstream of Aβ, Mic.13 between Aβ and tau, Ast.10 downstream of tau and upstream of cognitive decline, and Oli.7 downstream of Ast.10. Arrows denote modeled directional paths; blue and red arrows indicate positive and negative standardized path coefficients, respectively. Path labels show standardized coefficients, with significance indicated by \**P* < 0.05, \*\**P* < 0.01 and \*\*\**P* < 0.001. Model fit statistics are shown beneath each panel: Comparative Fit Index (CFI), Root Mean Square Error of Approximation (RMSEA), and Standardized Root Mean Square Residual (SRMR). **(J)** Alternative structural equation models testing the positional specificity of Ast.10 within the proposed cascade in the combined cohort. The original model was modified by exchanging the position of Ast.10 with Mic.13 (left) or with tau (right), while retaining the remaining model structure. Blue and red arrows denote significant positive and negative standardized path coefficients, respectively; grey arrows denote non-significant paths. Path labels show standardized coefficients, with significance indicated by \**P* < 0.05, \*\**P* < 0.01 and \*\*\**P* < 0.001. Model fit statistics are shown below each model as in (I).

**Extended Data Figure 2.**
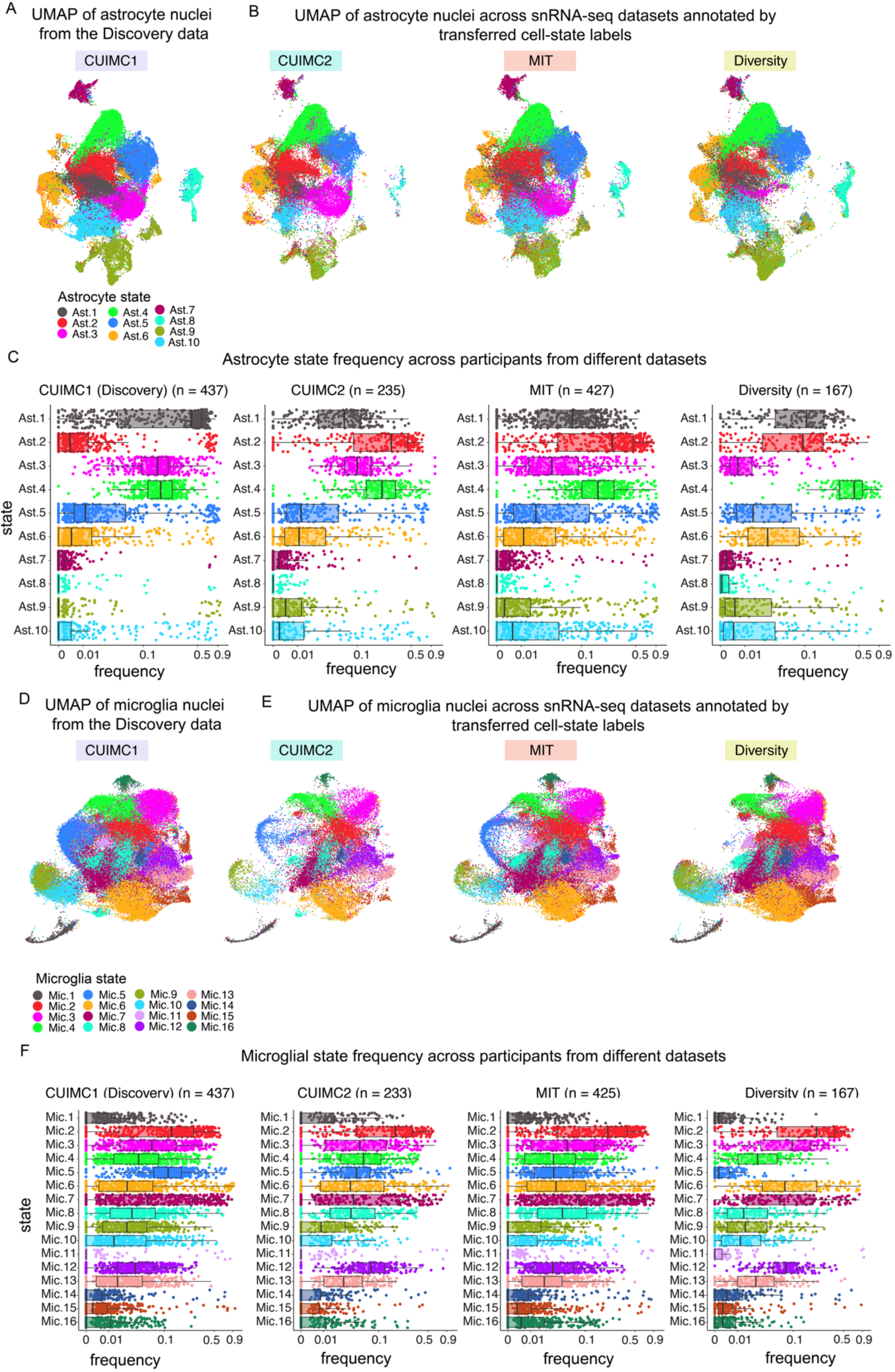
Uniform taxonomies of astrocytic and microglial states applied across four snRNA-seq datasets. **(A)** Uniform Manifold Approximation and Projection (UMAP) of individual astrocyte nuclei from the Discovery CUIMC1 dataset, colored by the astrocyte-state taxonomy defined in Green et al^1^. **(B)** UMAP embeddings of individual astrocyte nuclei from each dataset in the Replication set, mapped onto the reference UMAP space of CUIMC1 (shown in A). Each cell is colored by the predicted cell-state label, assigned using CUIMC1 as the reference (See Methods for details). The Replication set includes three snRNA-seq datasets: Columbia University Irving Medical Center dataset 2 (CUIMC2), Massachusetts Institute of Technology dataset (MIT)^3^ and the Accelerating Medicines Partnership for AD Diversity dataset (Diversity). **(C)** Frequency distributions of astrocytic states across participants in the four snRNA-seq datasets. **(D)** Uniform Manifold Approximation and Projection (UMAP) of individual microglia nuclei from the Discovery CUIMC1 dataset, colored by the microglial state taxonomy defined in Green et al^1^. **(E)** UMAP embeddings of individual microglia nuclei from each dataset in the Replication set, mapped onto the reference UMAP space of CUIMC1 (shown in A). Each cell is colored by the predicted cell-state label, assigned using CUIMC1 as the reference (See Methods for detail). The Replication set includes three snRNA-seq datasets: Columbia University Irving Medical Center dataset 2 (CUIMC2), Massachusetts Institute of Technology dataset (MIT)^3^ and the Accelerating Medicines Partnership for AD Diversity dataset (Diversity). **(F)** Frequency distributions of microglial states across participants in the four snRNA-seq datasets.

**Extended Data Figure 3.**
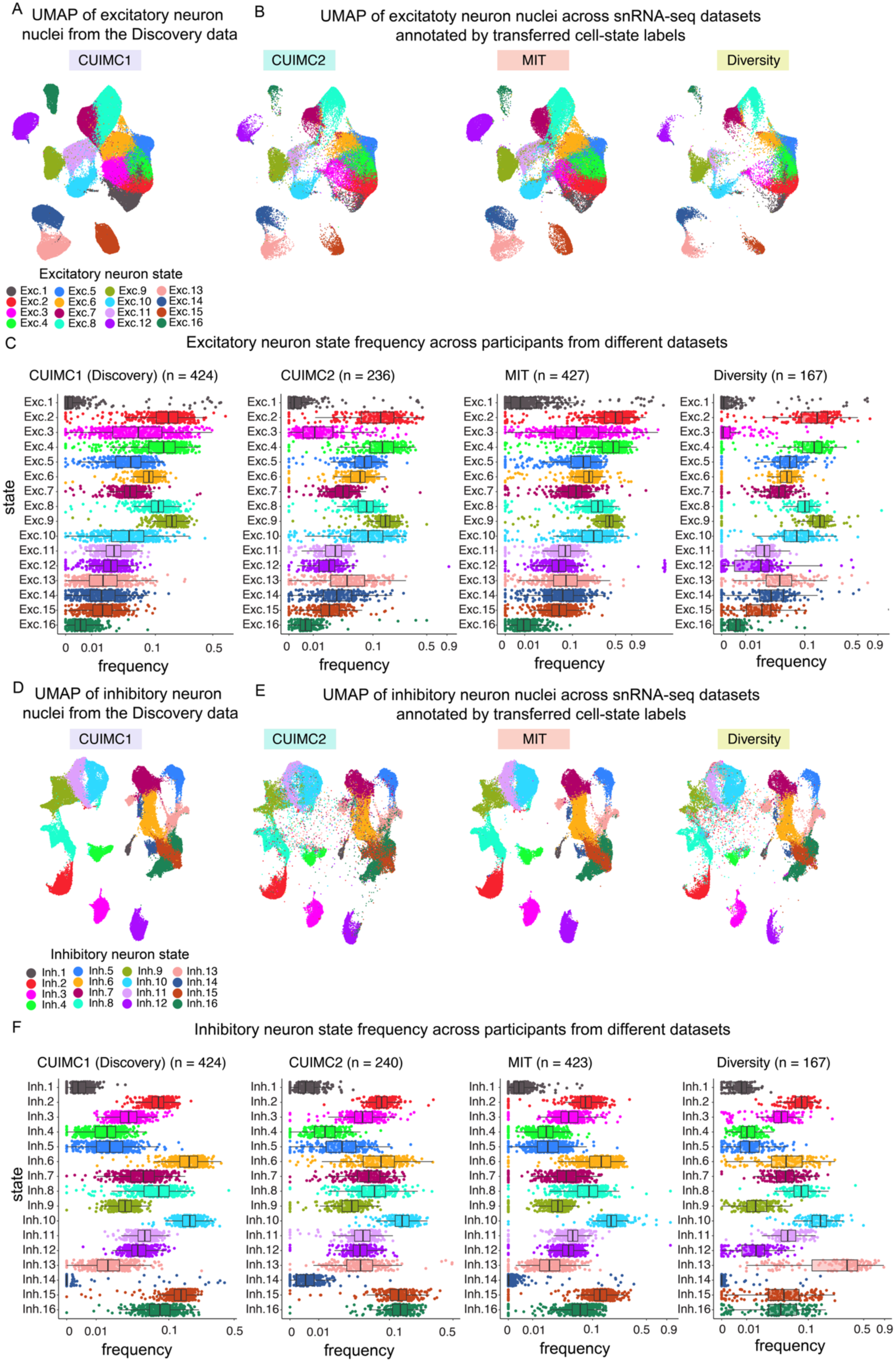
Uniform neuronal taxonomy applied across four snRNA-seq datasets. **(A)** Uniform Manifold Approximation and Projection (UMAP) of individual excitatory neuron nuclei from the Discovery CUIMC1 dataset, colored by the excitatory neuron cell-state taxonomy defined in Green et al^1^. **(B)** UMAP embeddings of individual excitatory neuron nuclei from each dataset in the Replication set, mapped onto the reference UMAP space of CUIMC1 (shown in A). Each cell is colored by the predicted cell-state label, assigned using CUIMC1 as the reference (See Methods for details). The Replication set includes three snRNA-seq datasets: Columbia University Irving Medical Center dataset 2 (CUIMC2), Massachusetts Institute of Technology dataset (MIT)^3^ and the Accelerating Medicines Partnership for AD Diversity dataset (Diversity). **(C)** Frequency distributions of excitatory neuron states across participants in the four snRNA-seq datasets. **(D)** Uniform Manifold Approximation and Projection (UMAP) of individual inhibitory neuron nuclei from the Discovery CUIMC1 dataset, colored by the inhibitory neuron cell-state taxonomy defined in Green et al^1^. **(E)** UMAP embeddings of individual inhibitory neuron nuclei from each dataset in the Replication set, mapped onto the reference UMAP space of CUIMC1 (shown in A). Each cell is colored by the predicted cell-state label, assigned using CUIMC1 as the reference (See Methods for details). The Replication set includes three snRNA-seq datasets: Columbia University Irving Medical Center dataset 2 (CUIMC2), Massachusetts Institute of Technology dataset (MIT)^3^ and the Accelerating Medicines Partnership for AD Diversity dataset (Diversity). **(F)** Frequency distributions of inhibitory neuron states across participants in the four snRNA-seq datasets.

**Extended Data Figure 4.**
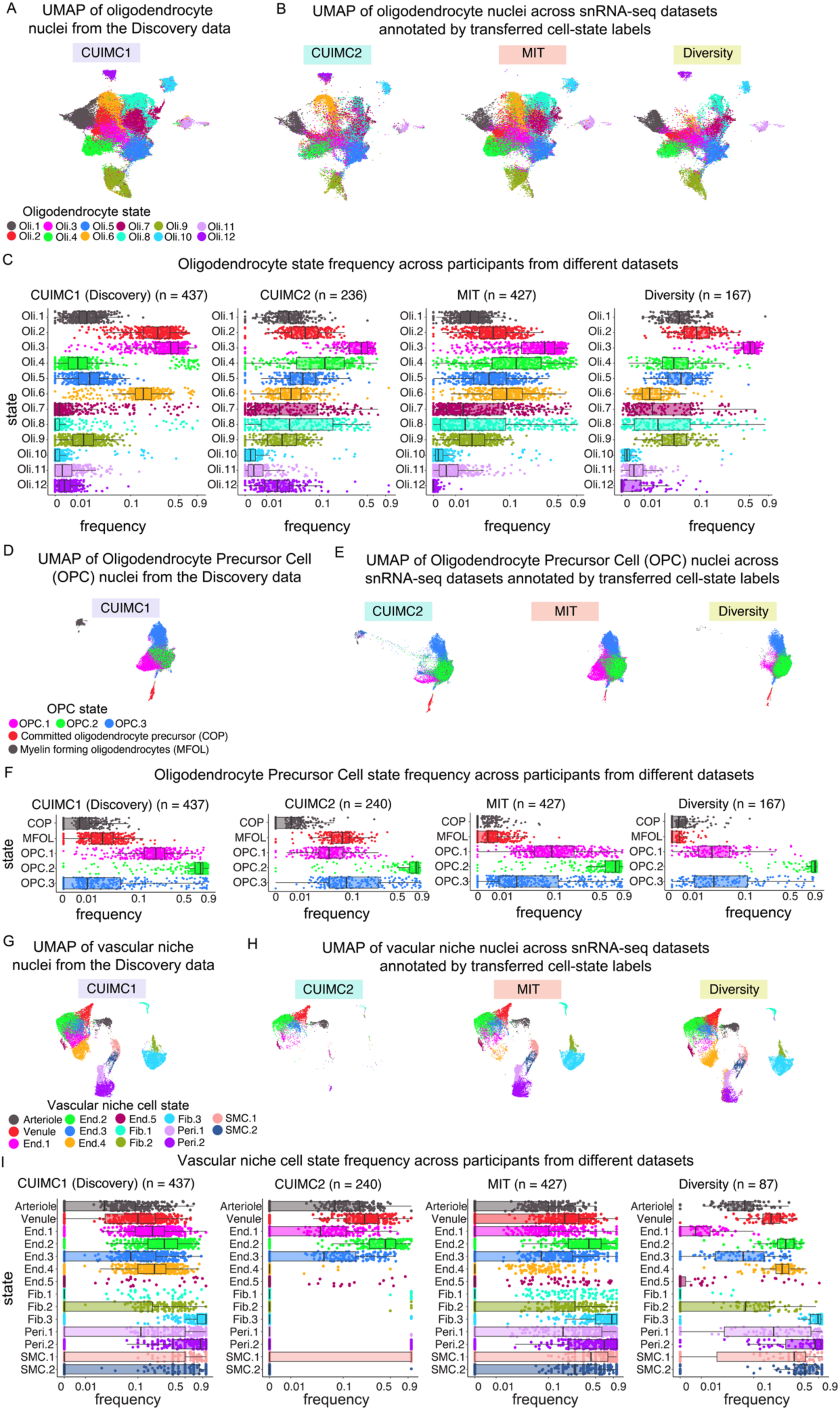
Uniform cell-state taxonomy for oligodendrocyte lineage and vascular niche population across four snRNA-seq datasets. **(A,D,G)** Uniform Manifold Approximation and Projection (UMAP) of oligodendrocyte nuclei (A), oligodendrocyte precursor cell (OPC) nuclei (D), and vascular niche nuclei (G) from the Discovery CUIMC1 dataset, colored by the cell-state taxonomy defined in Green et al^1^. **(B,E,H)** UMAP embeddings of oligodendrocyte nuclei (B), OPC nuclei (E), and vascular niche nuclei (H) from each dataset in the Replication set, projected onto the CUIMC1 reference UMAP space shown in A, D, and G. Cells are colored by predicted cell-state labels assigned using CUIMC1 as the reference (see Methods). The Replication set includes three snRNA-seq datasets: Columbia University Irving Medical Center dataset 2 (CUIMC2), Massachusetts Institute of Technology dataset (MIT)^3^ and the Accelerating Medicines Partnership for AD Diversity dataset (Diversity). **(C,F,I)** Participant-level frequencies of oligodendrocyte states (C), OPC states (F), and vascular niche cell states (I) across the four snRNA-seq datasets.

**Extended Data Figure 5.**
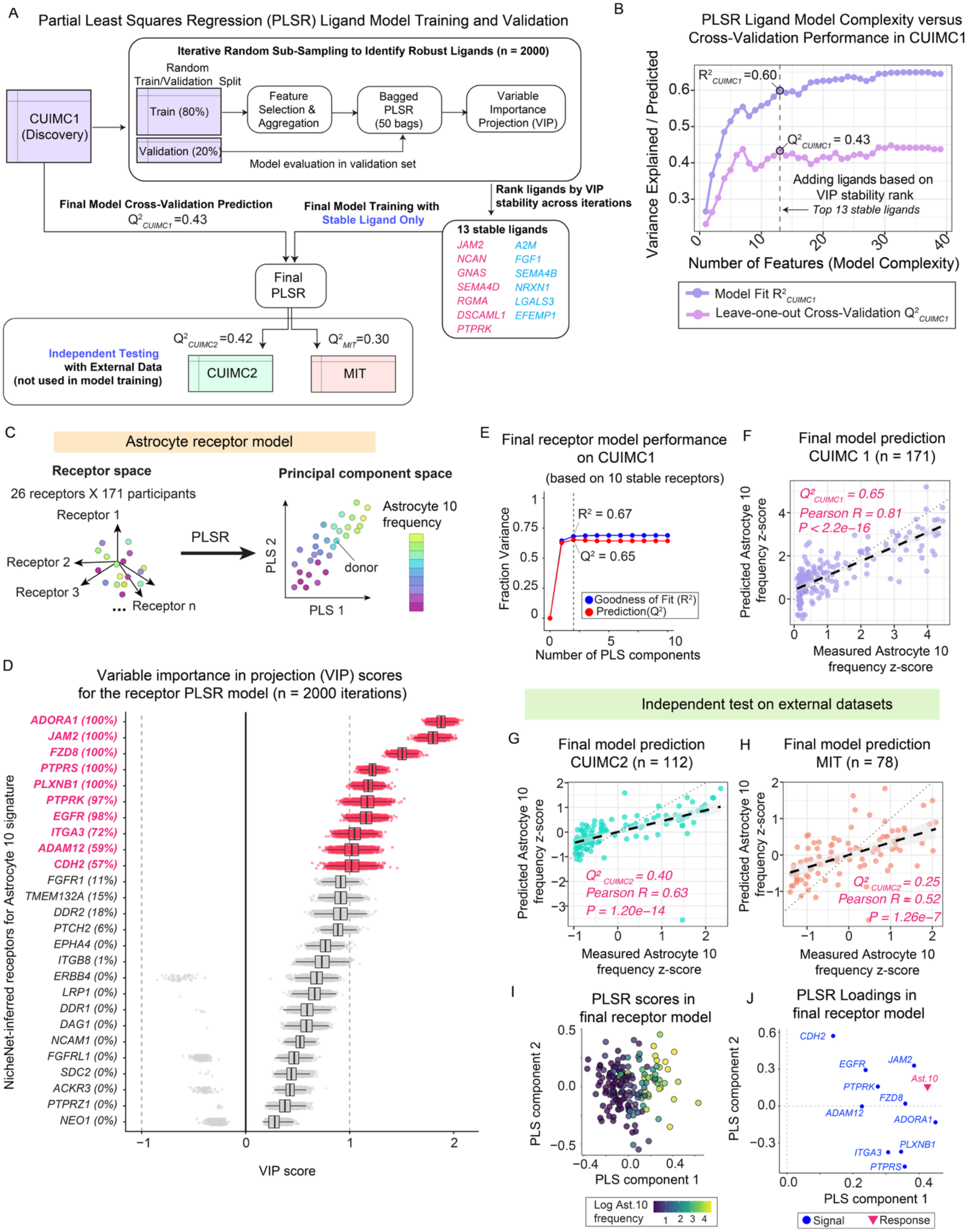
Partial Least Squares Regression modeling links ligands and receptors to Astrocyte state 10 frequency across participants. **(A)** Schematic of the ligand partial least squares regression (PLSR) model-training workflow. Using the CUIMC1 discovery cohort, PLSR models were trained in an iterative random sub-sampling framework (n = 2,000 iterations) to identify robust ligands associated with Astrocyte 10 (Ast10) frequency. In each iteration, samples were randomly split into training (80%) and validation (20%) sets. Feature selection and scaling were performed using the training set only, after which a bagged PLSR model was fit and evaluated on the validation set; ligands were then ranked by variable importance in projection (VIP). Ligands showing stable high VIP rankings across iterations were selected for the final model. A final PLSR model using these stable ligands was then trained in CUIMC1, with performance assessed by internal cross-validation, and subsequently evaluated in independent external test datasets (CUIMC2 and MIT), which were not used for model training. **(B)** Model complexity analysis in CUIMC1 to determine final model. Cumulative variance explained by the fitted model (R^2^CUIMC1) and leave-one-out cross-validation performance (Q^2^CUIMC1) are shown as ligands were incrementally added according to VIP stability rank. Cross-validation performance plateaued after inclusion of the top stable ligands, defining the final model complexity. **(C)** PLSR modeling predicts Ast10 frequency based on the NicheNet-prioritized receptors. For each donor, the input vector includes the pseudobulk astrocyte expression of 26 receptors. PLSR reduces the high-dimensional input receptor space to a principal component space, which captures covarying receptor patterns that predict Ast10 variation. Representative principal components (PLS 1 and PLS 2) are shown. **(D)** Stability and VIP scores of receptors in the PLSR model, quantified across 2,000 iterations of repeated random partitioning of training and validation sets. The magnitude of the VIP score reflects each receptor’s contribution to Ast10 frequency prediction, while its sign indicates positive or negative association with Ast10 frequency. Receptor stability is defined as the proportion of iterations in which |VIP≥ 1. Ten receptors selected for the final model based on stability ranking are highlighted. **(E)** Performance of the final PLSR model based on the 10 stable receptors, evaluated by variance explained (R^2^) and variance predicted (Q^2^) using leave-one-out cross-validation, plotted against the number of PLS components. The first two PLS components which achieve the optimal performance was highlighted. **(F)** Agreement between measured Ast10 frequency in CUIMC1 (n = 171) and leave-one-out cross-validation prediction by the final receptor PLSR model. Each point represents one participant. **(G-H)** Independent tests of the final receptor model with external datasets not used in model training. Predicted versus empirical Ast10 frequency z-scores in the CUIMC2 (G) and MIT (H) snRNAseq datasets. Predictions were made using the receptor PLSR model trained on the CUIMC1 Discovery dataset. **(I)** Donor-level PLSR scores for the first two components of the final receptor model, colored by log-transformed Ast10 frequency. **(J)** PLSR loadings (the first two PLS components) of the final receptor PLSR model for each receptor predictor and the response variable (Ast10 frequency).

**Extended Data Figure 6.**
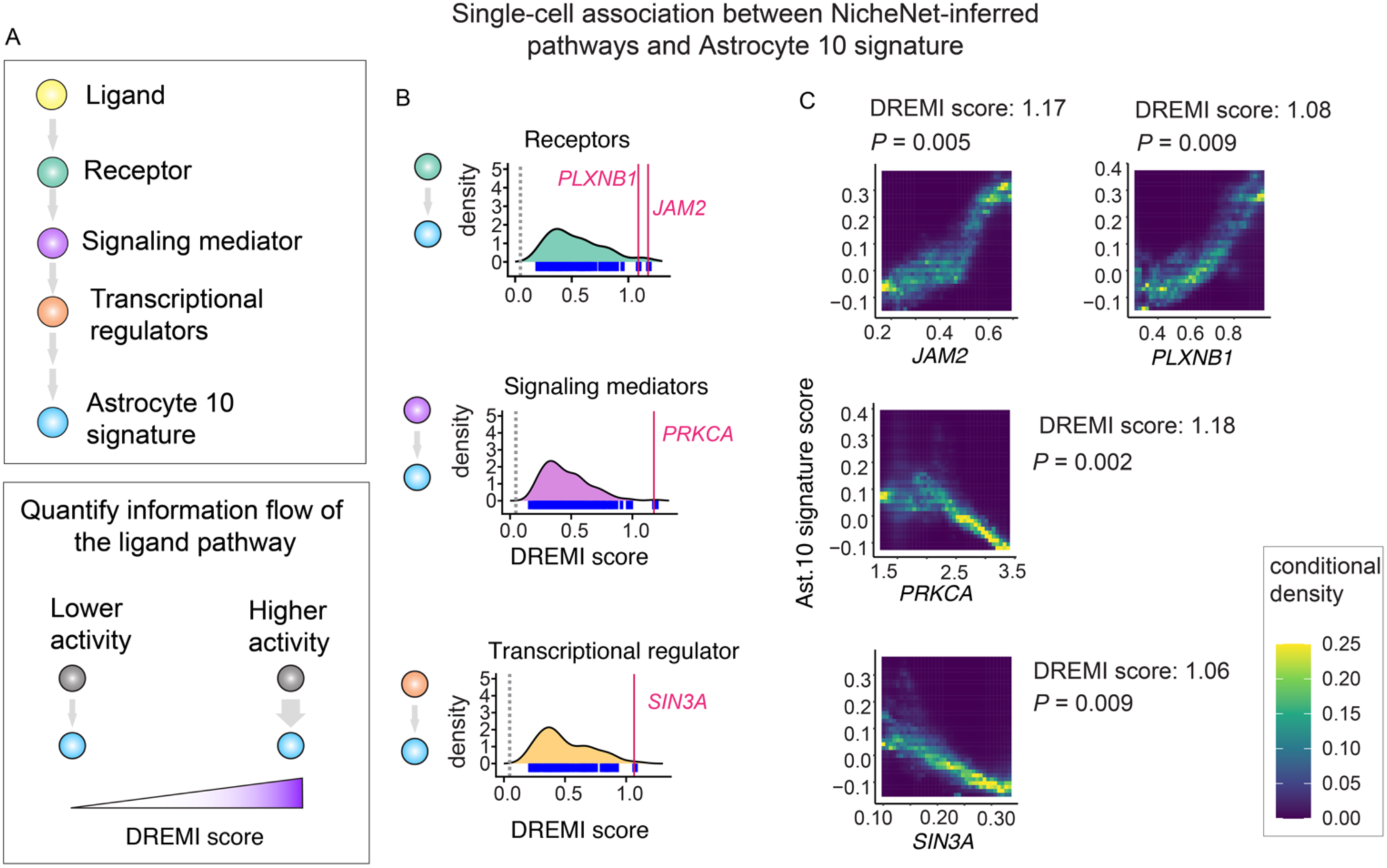
Pathway genes of the prioritized ligands share significant mutual information with the Astrocyte 10 signature at the single-cell level. **(A)** Schematic of the analysis quantifying the associations between prioritized ligand pathway genes and the Astrocyte 10 (Ast10) signature at the single-cell level. A simplified signaling cascade illustrates four signaling node categories: ligands, receptors, signaling mediators, and transcriptional regulators. To assess signal transfer activity, we applied Density Rescaled Mutual Information (DREMI)^52^ to quantify the mutual information between each signaling node and the Ast10 signature score across individual astrocytes in the Discovery snRNAseq dataset. A higher DREMI score indicates stronger signal transfer activity from the pathway node to the Ast10 transcriptional signature. **(B)** DREMI score distributions for each signaling node category: ligands, receptors, transcriptional regulators, and signaling mediators. Representative top-ranked pathway genes in each category are highlighted in red. Grey lines indicate the mean DREMI scores from shuffled data, serving as a baseline for comparison. **(C)** Density Rescaled Visualization of Information (DREVI)^52^ plots showing the relationship between representative pathway genes and the Ast10 signature score across individual astrocytes. Color intensity represents the conditional density distribution of astrocytes, illustrating how the Ast10 signature score (y-axis) varies with expression level of the pathway gene of interest (x-axis). DREMI scores and empirical *P*-values for each signaling node are displayed. *P*-values were derived from permutation tests across all possible genes in each category: receptors (n = 658), signaling mediators (n = 11,124), and transcriptional regulators (n = 1,079) (see Methods for details).

**Extended Data Figure 7.**
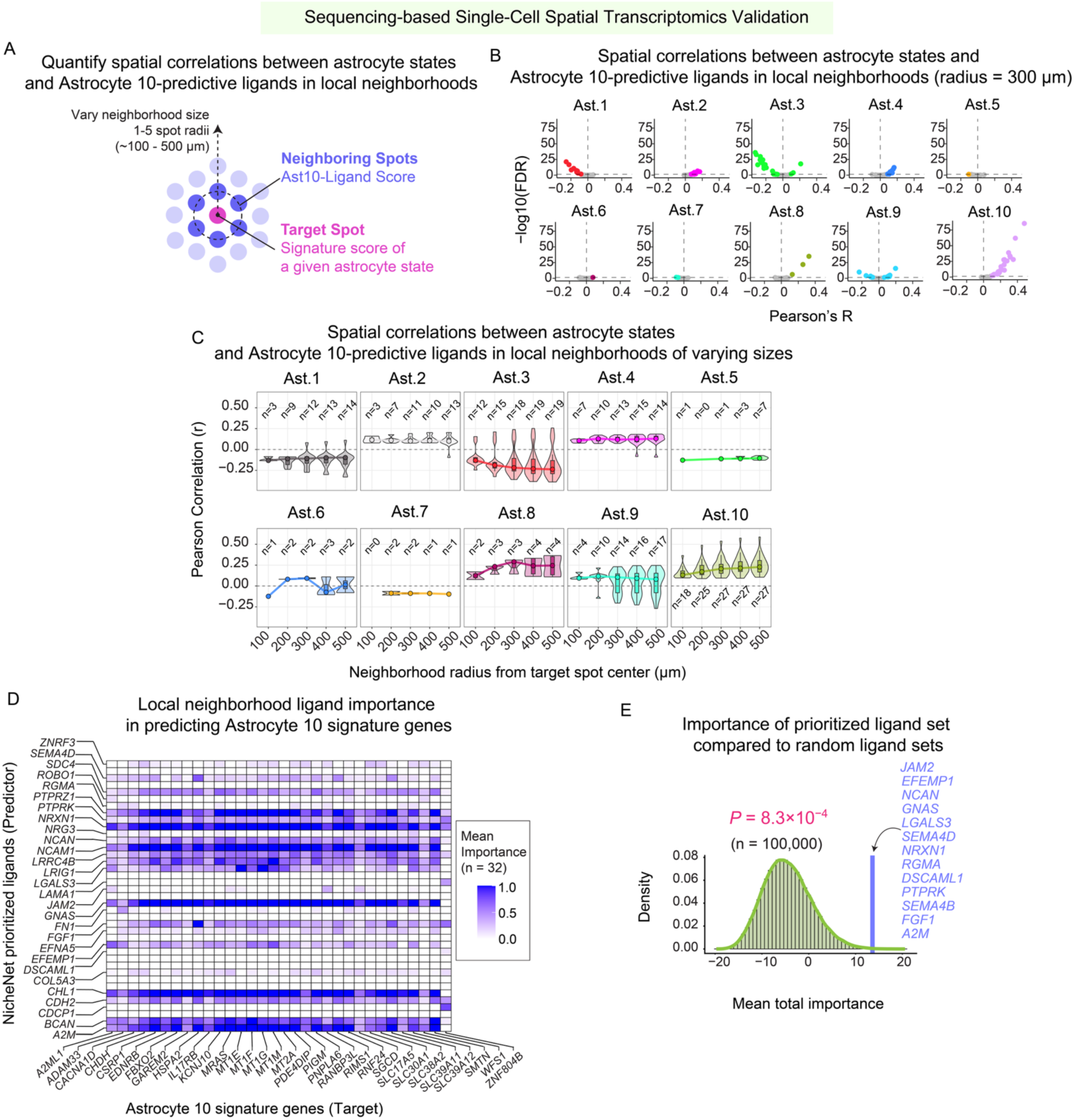
Prioritized ligands preferentially predict nearby Ast10 signature relative to other astrocytic states in spatial transcriptomic data. **(A)** Schematic of the Visium spatial correlation analysis. For each focal spot (pink), the astrocyte-state signature score at the focal spot was correlated with the neighborhood ligand signature score computed from neighboring spots (purple), excluding the focal spot itself. The neighborhood ligand signature was defined using prioritized Ast10-predictive ligands. Neighborhood size was varied from 1 to 5 spots (approximately 100-500 µm). **(B)** Volcano plots summarizing within-slide Pearson’s correlations in Visium data, as described in (A). Each point represents one tissue sample and shows the Pearson’s correlation statistics computed across spots between the focal-spot astrocyte-state signature and the neighborhood ligand signature score. The ligand signature was defined using prioritized Ast.10-predictive ligands with significantly positive VIP scores in at least 50% of training iterations. For each focal spot, the ligand signature score was averaged across neighboring spots within a 300 µm radius, excluding the focal spot itself. Astrocyte state signatures were derived from differentially upregulated marker genes, except for Ast1, for which downregulated markers were used because no upregulated markers were identified. For interpretability, Pearson’s R values for Ast1 were inverted so that negative correlations indicate depletion of the Ast1 signature (see Methods). Each point represents one tissue sample; significant correlations (false discovery rate, *FDR* < 0.05) are highlighted. **(C)** Sensitivity analysis of spatial correlations between the prioritized Ast10-predictive ligand signature and astrocyte-state signatures (Ast1-Ast10) across neighborhood sizes of approximately 100-500 µm. Violin plots show the distribution of Pearson’s correlation coefficients across tissue samples with significant spatial correlations at each neighborhood size; the number of significant samples (out of 32 total samples) is indicated. State signatures were derived from differentially upregulated marker genes, except for Ast1, for which downregulated markers were used because no upregulated markers were identified. For interpretability, Pearson’s R values for Ast1 were inverted so that negative correlations indicate depletion of the Ast1 signature (see Methods). **(D)** MISTy-derived importance scores quantifying the local contribution of NicheNet-prioritized ligands within a 300 µm radius to prediction of individual Ast10 signature genes, averaged across 32 spatial transcriptomic slides from 17 donors. Importance scores > 0 indicate significant influence. (see Methods for metric details). **(E)** Total MISTy-derived importance scores of the 13 PLSR-prioritized ligands compared to randomly sampled ligand sets of the same size (see Methods). Empirical *P*-value was calculated from 100,000 random samplings. The prioritized ligand set is indicated.

**Extended Data Figure 8.**
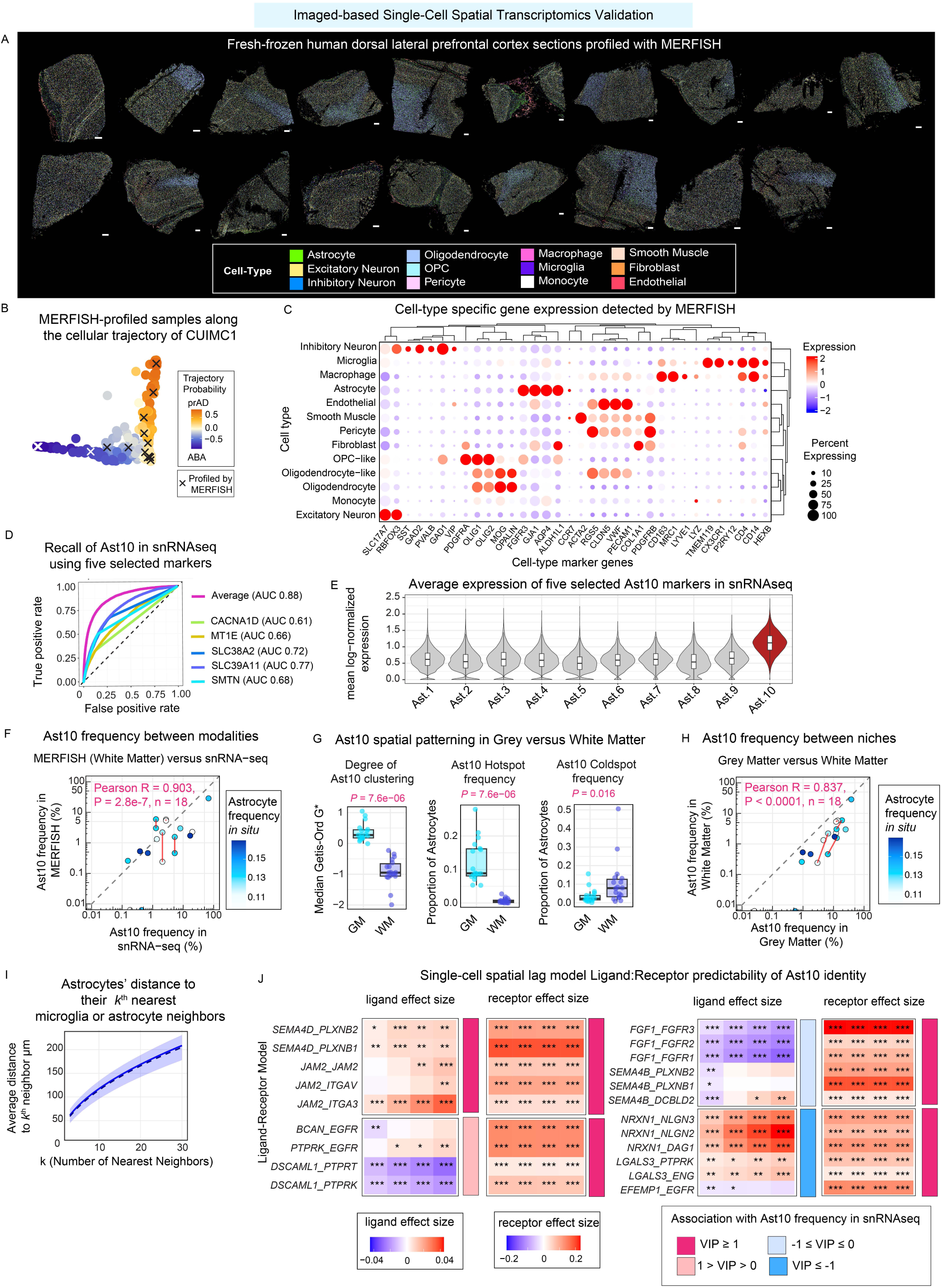
Prioritized ligand and receptor pairs predict Ast10 cells at the single-cell level in DLPFC tissue profiled using MERFISH spatial transcriptomics. (A) 19 fresh-frozen human dorsal lateral prefrontal cortex (DLPFC) tissue sections profiled by MERFISH, visualized in physical coordinates, with individual segmented nuclei colored by their cell type annotation. Scale bar: 500µm (B) Distribution of CUIMC1 donors with matched snRNA-seq and MERFISH profiling data along the cellular trajectories defined by Green et al. (C) Expression of canonical cell-type marker genes across major cell types annotated in the 19 MERFISH-profiled tissue sections. Dot size indicates the proportion of cells expressing each gene, and color indicates scaled expression level. (D) Receiver operating characteristic (ROC) curves evaluating the ability of five selected Ast10 marker genes (*SLC38A2, SMTN, SLC39A11, CACNA1D* and *MT1E*), individually or as a combined mean-expression score, to distinguish Ast10 from other astrocyte states in the *CUIMC1* snRNA-seq dataset. AUC: Area under the ROC curves.The dashed diagonal indicates random classification (AUC = 0.5). (E) Distribution of the average log-normalized expression of the five selected markers across the ten astrocyte states (Ast1-Ast10) in the *CUIMC1* snRNA-seq dataset. Violin plots show the expression distribution across individual nuclei, with embedded boxplots indicating the median and interquartile range; whiskers extend to 1.5 times the interquartile range. Ast10 is highlighted in red. (F) Pearson’s correlation between donor-level Astrocyte 10 (Ast10) frequency estimated from snRNA-seq (CUMIC1) and Ast10 frequency measured in white matter regions by MERFISH. Red lines connect tissue sections from the same donor. Points are colored by the relative abundance of astrocytes among all annotated cell types. (G) Local spatial statistics comparing Ast10 spatial patterning between grey (GM) versus white matter (WM) across 19 tissue sections. Left: median Getis-Ord G* scores, quantifying local clustering of astrocytes with relatively high or low Ast10 score; Middle: proportion of astrocytes with significantly positive Getis-Ord G* scores (hotspots); Right: proportion of astrocytes with significantly negative Getis-Ord G* scores (coldspots). *P*-values were calculated using paired Wilcoxon signed-rank test. (H) Pearson’s correlation in Ast10 frequency between grey and white matters across 19 MERFISH-profiled tissues. Red lines connect tissue sections from the same donor. Each data point is colored by the relative abundance of astrocytes among all annotated cell types in a given sample. (I) Average distance from each astrocyte to its *k^th^* nearest neighboring microglia or astrocytes, shown as a function of neighborhood size (*k*). Shaded regions indicate 95% confidence intervals. (J) Spatial lag modeling effect sizes for prioritized ligand-receptor pairs. Models quantify their spatial colocalization with Ast10 scores across multiple neighborhood scales. For each ligand-receptor pair, Ast10 scores in a target astrocyte are modeled as a function of the average ligand expression in the *k*-nearest microglia or astrocytes and receptor expression in the target astrocyte, adjusting for other technical covariates (see Methods). *k* ranges from 3 to 12 neighbors (approximately 50-126 µm, as shown in G). For visualization, ligand-receptor pairs are grouped by whether they are positively or negatively correlated in Ast10 frequency in snRNAseq data analysis (Figure 3, Extended Data Figure 5). Asterisks: false discovery rate (*FDR*). ***, *FDR* < 0.001; **, *FDR* < 0.01; *, *FDR* < 0.05)

**Extended Data Figure 9.**
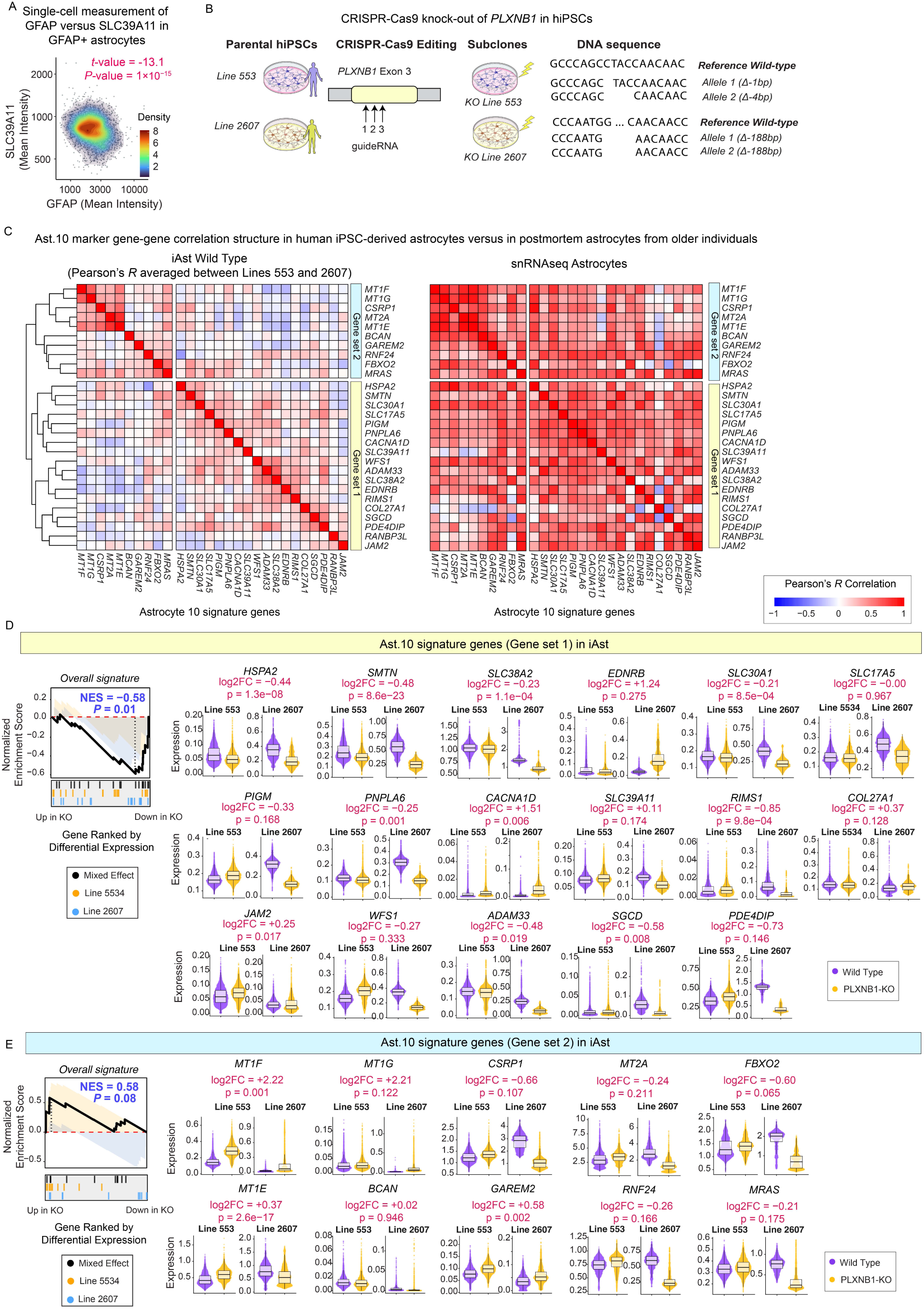
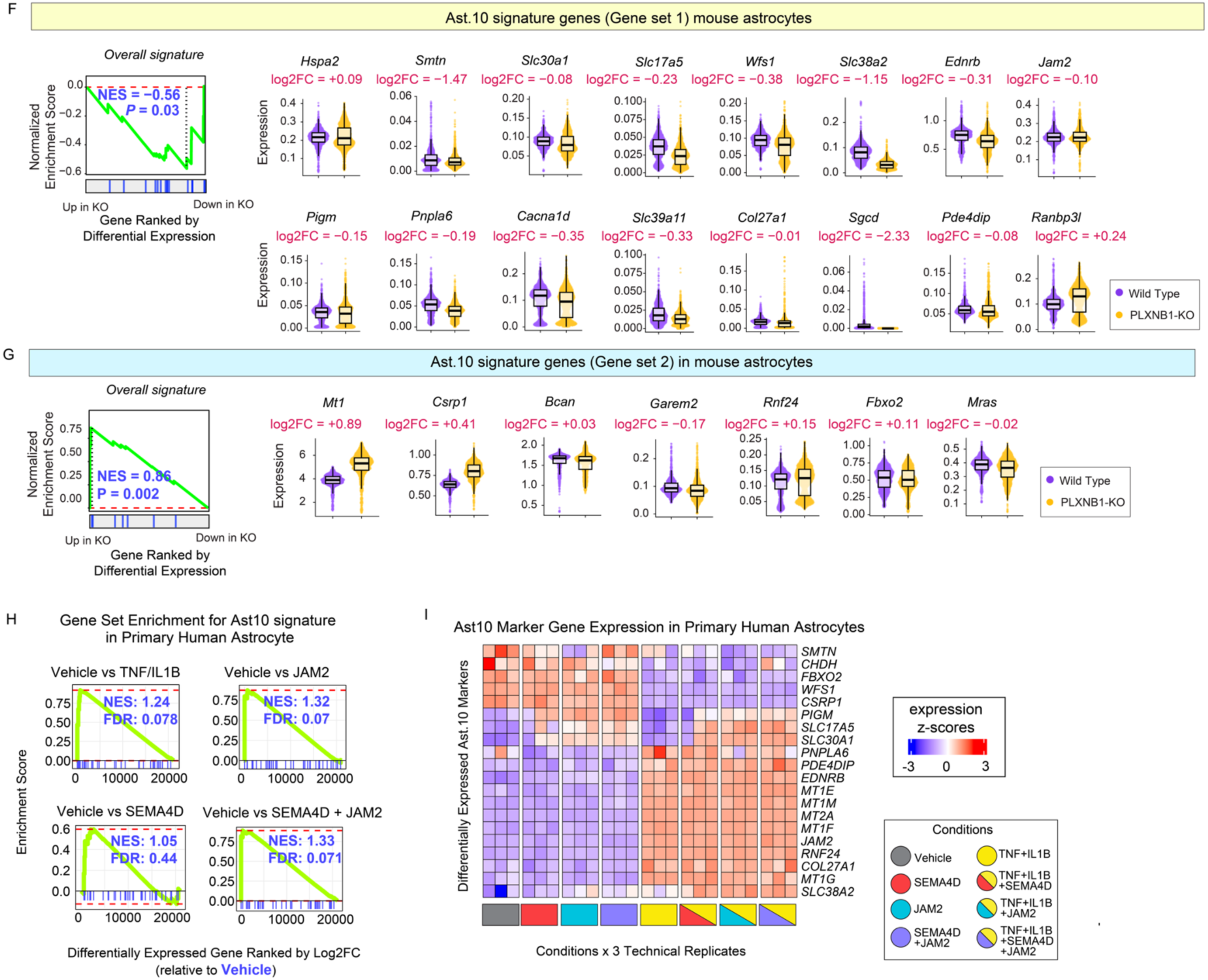
Validation of the role of PLXNB1 on Ast10 program. **(A)** Single-cell density plot showing the covariation between SLC39A11 and GFAP expression, quantified based on mean fluorescence intensity measured from immunofluorescence imaging of an AD donor. Each data point represents an astrocyte identified by GFAP-based segmentation across 40 images. Statistics indicate the effect size and significance of GFAP expression for predicting SLC39A11 levels, adjusting for SLC39A11 expression and image variability, derived from fitting a mixed effect model (See Methods for details). **(B)** Schematic of *PLXNB1* gene editing in two parental human induced pluripotent stem cell (hiPSC) donor lines (Lines 553 and 2607). Three guide RNAs targeting exon 3 of *PLXNB1* were used to generate clonal knockout lines by CRISPR-Cas9, as previously described^64^. Sanger sequencing confirmed frameshift deletions in the resulting PLXNB1-knockout subclones. **(C)** Gene-gene correlation structure of Ast10 marker genes in hiPSC-derived astrocytes (iAst, wild type) and postmortem astrocytes from older individuals in the discovery snRNA-seq dataset. For each gene pair, Pearson’s correlation was computed separately in each wild-type iAst line using MAGIC-imputed expression values across all astrocytes, and correlations were averaged across Lines 553 and 2607. Heatmaps show the resulting correlation matrices for iAst and snRNA-seq astrocytes; genes were grouped by hierarchical clustering using complete linkage based on the averaged iAst wild type correlation structure. **(D,E)** Single-cell expression of Ast10 signature genes in wild-type and PLXNB1-knockout iAst derived from the two donor lines, shown separately for (D) Ast10 Gene Set 1 and (E) Ast10 Gene Set 2. Left, gene set enrichment analysis assessing enrichment of each Ast10 gene set in *PLXNB1*-knockout relative to wild-type cells, using on genes ranked by differential expression (Wald statistics) from the Nebula^117^ mixed- effects model; normalized enrichment score (NES) and *P*-value are indicated. Colored ticks denote gene positions in the ranked list, with line-specific ranking shown for Lines 553 and 2607 and the combined mixed-effects model indicated in black. Right, violin plots show single-cell expression distributions for individual Ast10 genes in each line, with log2 fold-change (log2FC) and *P-*value from differential expression analysis using the Nebula mixed-effect framework (see Methods). **(F,G)** Validation of the Ast10 program in mouse astrocytes with and without *Plxnb1* knockout. Left, gene set enrichment analysis for Ast10 Gene Set 1 (F) and Gene Set 2 (G), using genes ranked by differential expression (Wald statistics) in *Plxnb1*-knockout versus wild-type mouse astrocytes; normalized enrichment score (NES) and *P*-value are indicated. Violin plots show single-cell expression distributions for individual Ast10 signature genes; log2FC from differential gene expression analysis is indicated above each gene. **(H)** Gene set enrichment analysis of the Ast10 signature in primary human astrocytes under the indicated stimulation conditions relative to vehicle control. Cells were treated with Vehicle, SEMA4D, JAM2, SEMA4D + JAM2, TNF + IL1B, TNF + IL1B + SEMA4D, TNF + IL1B + JAM2, or TNF + IL1B + SEMA4D + JAM2. For each differential gene-expression comparison, genes were ranked using a composite score calculated as the DESeq2^121^ Wald statistics multiplied by the -log10 transformed *P* value. **(I)** Heatmap showing z-scored expression of differentially expressed Ast10 markers across stimulation conditions in primary human astrocytes, with three technical replicates per condition.

**Extended Data Figure 10.**
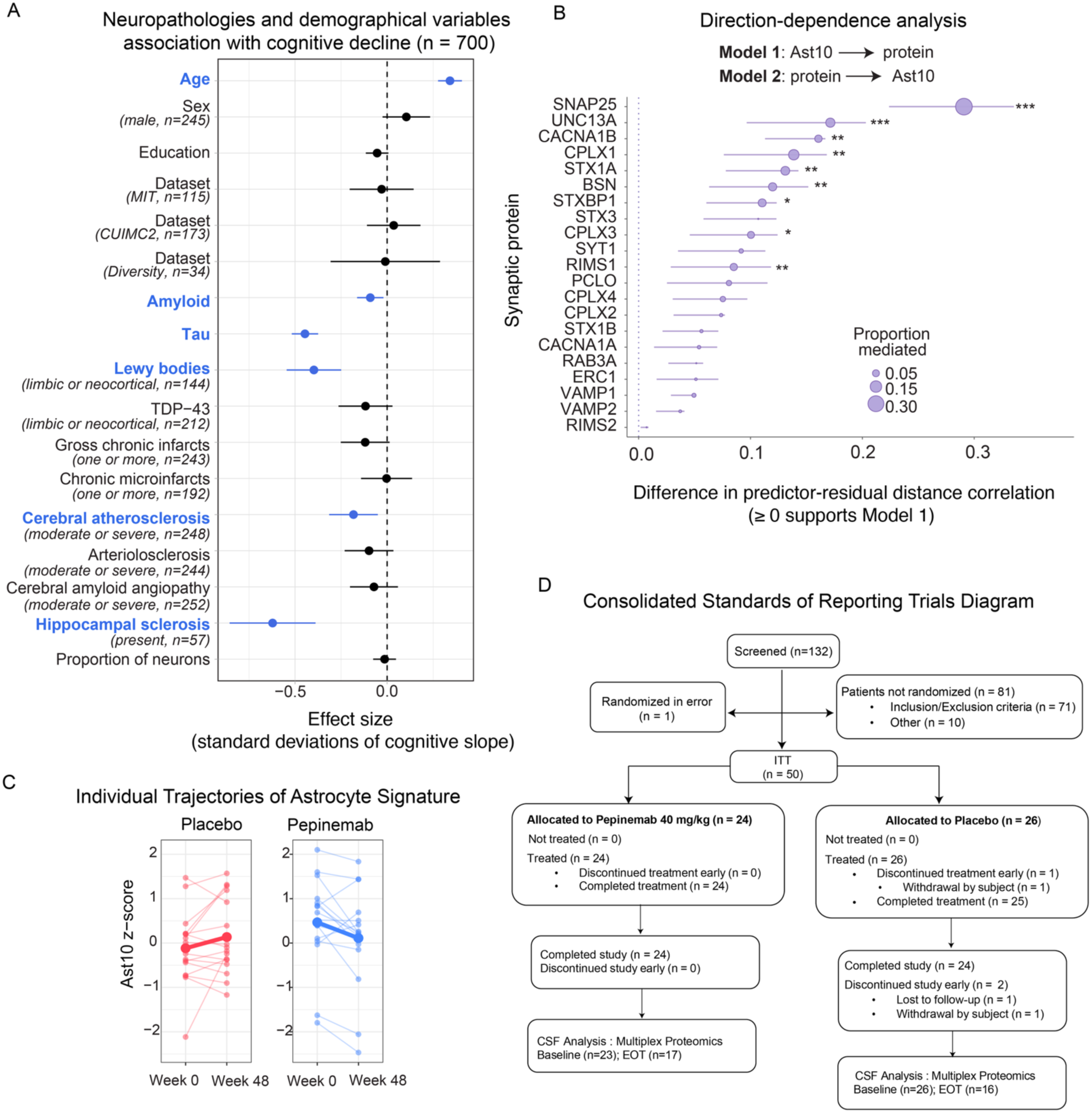
Neuropathological and demographic correlates of cognitive decline and longitudinal Ast10 proteomic signature trajectories in the pepinemab trial. **(A)** Forest plot from a multivariate regression model predicting cognitive decline based on pathologic variables and demographic variables (y-axis) in the Rush University cohorts. Effect sizes (dots) and the respective 95% confidence intervals (error bars) are shown. Significant variables are highlighted in blue. All continuous variables were z-scored. Categorical variables were dichotomized; their respective factor levels and case numbers are indicated in brackets. A total of n = 700 samples across all available snRNAseq datasets (Discovery + Replication) with complete measurements for all variables were analyzed. **(B)** Direction-dependence analysis assessing the directional ordering between Ast10 frequency and synaptic protein abundance. For each protein, two competing models were compared: Model 1, Ast10 → protein, and Model 2, protein → Ast10. Directional support was quantified as the difference in distance correlation between predictor and model residuals for Model 2 versus Model 1; positive values indicate lower residual dependence, and therefore greater support for the hypothesized Ast10 → protein direction. Points show the estimated difference in distance correlation and error bars indicate 95% confidence intervals. Point size denotes the proportion of the Ast10-cognitive decline association mediated by each protein in the corresponding mediation analysis. Models were adjusted for sex, post-mortem interval, dataset, age at death, education, total cell count, and median gene count per participant. Asterisks indicate significance of the mediation test: \**P <* 0.05*, **P <* 0.01, \*\*\**P <* 0.001. **(C)** Ast10 proteomic signature scores (Ast10 z-score) at week 0 and week 48 across for individual participants in the AD-SIGNAL trial treated with either placebo or pepinemab. Thin lines indicate individual trajectories and thick lines indicate group averages at the indicated timepoints. **(D)** Consolidated Standards of Reporting Trials (CONSORT) Diagram detailing the participant flow in the SIGNAL-AD trial.

